# Multi-Ancestry Meta-Analysis yields novel genetic discoveries and ancestry-specific associations

**DOI:** 10.1101/2021.04.23.441003

**Authors:** Patrick Turley, Alicia R. Martin, Grant Goldman, Hui Li, Masahiro Kanai, Raymond K. Walters, Jonathan B. Jala, Kuang Lin, Iona Y. Millwood, Caitlin E. Carey, Duncan S. Palmer, Meghan Zacher, Elizabeth G. Atkinson, Zhengming Chen, Liming Li, Masato Akiyama, Yukinori Okada, Yoichiro Kamatani, Robin G. Walters, Shawneequa Callier, David Laibson, Michelle N. Meyer, David Cesarini, Mark Daly, Daniel J. Benjamin, Benjamin M. Neale

## Abstract

We present a new method, Multi-Ancestry Meta-Analysis (MAMA), which combines genome-wide association study (GWAS) summary statistics from multiple populations to produce new summary statistics for each population, identifying novel loci that would not have been discovered in either set of GWAS summary statistics alone. In simulations, MAMA increases power with less bias and generally lower type-1 error rate than other multi-ancestry meta-analysis approaches. We apply MAMA to 23 phenotypes in East-Asian- and European-ancestry populations and find substantial gains in power. In an independent sample, novel genetic discoveries from MAMA replicate strongly.

## INTRODUCTION

The past decade has seen the discovery of hundreds of thousands of credible genetic associations for complex traits and diseases^1^. Many of these discoveries were identified by meta-analyzing genome-wide association studies (GWAS)^2–4^ conducted in multiple cohorts, boosting statistical power relative to GWAS in any one cohort. GWAS summary statistics are now facilitating progress in many areas, including novel drug development^5,6^ and disease prediction for potential clinical application^7^.

Unfortunately, GWAS research to date has overwhelmingly been conducted in European-ancestry samples. Consequently, GWAS estimates for European populations are substantially more precise than those for other populations, potentially generating unequal gains from the scientific advances. Clearly, correcting these imbalances requires intensifying data-collection efforts in non-European populations^8,9^, and a number of promising efforts are underway^10–14^. However, it will take years before levels of precision are reached comparable to those currently available for European-ancestry studies. Even then, for many traits, further increases in precision would almost certainly still be valuable.

To complement the current efforts to improve the precision of GWAS effect-size estimates for non-European-ancestry populations, we have developed a new method, Multi-Ancestry Meta-Analysis (MAMA), that efficiently shares information across populations via cross-ancestry GWAS meta-analysis. Currently, researchers who wish to meta-analyze summary statistics from different-ancestry cohorts must confront two well-known obstacles. First, the populations may differ in allele frequencies and linkage disequilibrium (LD) patterns. Second, the effects of SNPs may not be identical across the two populations. Both obstacles induce heterogeneity in marginal (i.e., GWAS) effect sizes between populations. Consequently, a naïve meta-analysis of summary statistics that ignores these obstacles would generally result in biased estimates for both populations.

In different contexts, the two obstacles above have each been addressed separately in previously developed methods. For example, POPCORN directly models allele-frequency and LD differences to estimate genetic correlation between populations^15^. MTAG accounts for heterogeneity of marginal effect sizes across GWAS summary statistics in a one-population, multi-trait setting^16^. MAMA adapts and combines strategies from both methods, accounting for differences between populations in conditional effects, allele frequencies, and LD.

Several methods have been developed for cross-ancestry meta-analysis^17–21^. Compared to these, MAMA has a unique combination of attractive features:

i. it only requires GWAS summary statistics and a reference panel for each population;
ii. under plausible assumptions discussed below, it is the best linear unbiased estimator, but also appears robust under alternative assumptions;
iii. it has a linear, closed-form solution, ensuring fast computation time.

For MAMA’s intuition, first consider an inverse-variance-weighted (IVW) meta-analysis^18^ applied to GWAS summary statistics with no heterogeneity (e.g., each GWAS sample is from the same population). In such a case, IVW is unbiased and efficient. However, if there is heterogeneity in the marginal effect sizes between populations, then the estimate from the alternate ancestry needs to be adjusted to avoid bias, and the relative weight applied to the alternate ancestry’s estimate must be reduced to be efficient^16^. Before meta-analyzing, MAMA estimates the heterogeneity across ancestries for each SNP and optimally adjusts the estimates and relative weights on the alternate ancestries.

To evaluate MAMA, we apply it to GWAS summary statistics for a wide range of anthropometric, health, and behavioral phenotypes in populations with European and East-Asian ancestries. MAMA identifies many new loci that were not found using either of the single-ancestry GWAS results. For some phenotypes, we calculate that MAMA generates increases in power approximately equivalent to an increase of hundreds of thousands of individuals in the East Asian discovery sample. In comparisons of the signs of estimates from MAMA-identified loci to the signs of GWAS estimates from independent samples, MAMA’s replication record is strong.

For a less technical description of the paper and of how MAMA results should— and should not—be interpreted, see the Frequently Asked Questions (FAQs in Supplementary Information).

## RESULTS

### The MAMA Framework

MAMA is an extension of the related method, MTAG^16^, adapting its assumptions to the multi-ancestry context and generalizing it to allow for differences across ancestries. To make the relationship clear, our derivation of MAMA is parallel to Turley et al.’s^16^ derivation of MTAG. We highlight where the derivations differ, due to modeling the conditional effects of SNPs (rather than marginal associations of SNPs) and differences in allele frequencies and LD, all of which are crucial for cross-ancestry meta-analysis. More details are in the supplementary materials.

#### Background

To begin, let *y*_*p,i*_ denote the phenotype of interest for individual *i* in population *p*, normalized to have mean zero within its population. Let ***x***_*p,i*_ denote the vector of genotypes for the set of all SNPs in the reference panel, including unmeasured SNPs. We assume that genotypes are mean centered within their population, but all of our results below generalize to any assumption about the genotype units used (e.g., raw allele counts, standard deviations of allele counts), but we assume that genotypes are mean centered within their population. Let ***r***_*p*_ ≡ Var(***x***_*p,i*_) denote the variance-covariance matrix (i.e., LD matrix) of the genotype vector in population *p*.

We use an additive model: *y*_*p,i*_ = ***x***_*p,i*_ ***b***_*p*_ + *ε*_*p,i*_, where ***b***_*p*_ is the vector of conditional effects in population *p*. We treat the effects across each of *P* populations {***b***_1_, …, ***b***_*P*_} as random and allow SNPs’ effects to be correlated across populations.

MAMA’s key assumption is that the effects have constant (co)variance across SNPs: Var(*b*_1,*j*_, …, *b*_*P,j*_) = ***ω*** for all SNPs *j*, where ***ω*** is a *P* × *P* positive-semidefinite matrix. We call this the “homogeneous-***ω***” assumption. A special case is perfect genetic correlation, where the conditional effects of SNPs are equal or proportional across populations. As we discuss below, perfect genetic correlation may be reasonable to assume in some cases (as we do in our real-data analysis). However, MAMA allows for the genetic correlation and variances to be any values, as long as they are the same across SNPs. MAMA’s key assumption differs from MTAG’s key assumption because MTAG assumes constant (co)variance of *marginal* effects, whereas MAMA assumes constant (co)variance of *conditional* effects (conditioning on all SNPs in the reference panel). MAMA’s assumption is more plausible in the multi-ancestry context because the distribution of marginal effects should depend on LD structure.

The GWAS estimate for SNP *j* in population 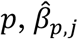, is an estimate of the marginal association. The estimand of MAMA is similarly the marginal association *β*_*p,j*_. As inputs, MAMA uses (i) the vector of GWAS effect-size estimates for SNP *j* across populations, 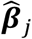, and their standard errors across multiple populations and (ii) an LD matrix ***r***_*p*_ for each population from a reference panel. For the reference panel, in our applications, we use data from the 1000 Genomes Project^22^, but any reference panel corresponding to the ancestries of the GWAS samples would also work.

#### Summary of MAMA Derivation

To define the two matrices needed for the estimator, note that the *p*^th^ element of 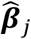 is a function of the population *p*’s true conditional SNP effects ***b***_*p*_, LD matrix ***r***_*p*_, and error:

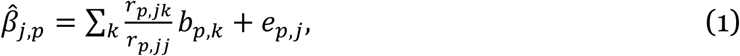

where *r*_*p,jk*_ is the (*j, k*)^th^ element of ***r***_*p*_, *b*_*p,k*_ is the *k*^th^ element of ***b***_*p*_, and *e*_*p,j*_ includes sampling variation and confounding biases. Thus, the vector 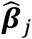 can be decomposed into the true marginal association ***β***_*j*_ plus error: 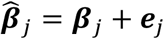. The two matrices we need are the variance-covariance matrix of the true marginal associations conditional on the LD matrices, denoted **Ω**_*j*_ ≡ Var(***β***_*j*_ |{***r***_1_, …, ***r***_*P*_}), and the variance-covariance matrix of the error, denoted **Σ**_*j*_ ≡ Var(***e***_*j*_|{***r***_1_, …, ***r***_*P*_}). Because samples drawn from different populations do not overlap, **Σ**_*j*_ is diagonal.

MAMA is a Generalized Method of Moments (GMM) estimator^23^. It is defined by a vector of *P* moment conditions, 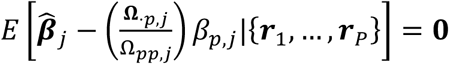, where **Ω**._*p,j*_is the *p*^th^ column of **Ω**_*j*_ and Ω_*pp,j*_ is the (*p, p*)^th^ element of **Ω**_*j*_. Each moment condition is a necessary condition for a best linear unbiased estimate of the marginal association for SNP *j* in population *p*. The MAMA estimator is the efficient GMM estimator based on these moment conditions:

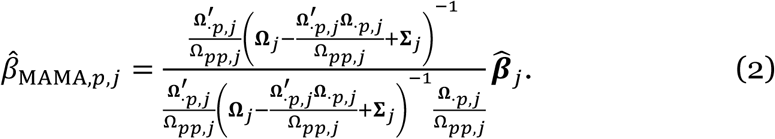

Two special cases may help with intuition. First, suppose conditional effect sizes are equal in all populations, and consider a SNP where LD patterns are identical between populations. Then all elements of **Ω**_*j*_ are equal, and equation (2) specializes to the formula for inverse-variance-weighted (IVW) meta-analysis: 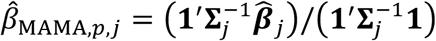, where **1** is a vector of ones. Second, suppose conditional effect sizes are uncorrelated across populations or LD patterns are uncorrelated at a particular SNP. Then **Ω**_*j*_ is diagonal, and equation (2) sets each population’s MAMA estimate equal to the population’s GWAS estimate: 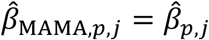.

More generally, when conditional effect sizes and/or LD patterns are imperfectly correlated, MAMA produces estimates in between these two special cases. MAMA is unbiased because it optimally modifies the GWAS estimates from other populations before meta-analyzing them with population *p*: MAMA deflates another population’s GWAS estimates when the correlation with population *p*’s GWAS estimates is smaller and inflates them when the other population’s heritability is smaller than *p*’s (this is the 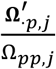 term in the numerator of equation (2)). MAMA has the minimum variance among linear unbiased estimators because, in addition to putting lower relative weight on estimates with greater sampling variance (as in IVW meta-analysis), it puts lower relative weight on estimates with greater heterogeneity in marginal effect sizes (the term in the inverse in the numerator).

MAMA’s moment conditions and estimator closely resemble MTAG’s. Like MTAG, MAMA is asymptotically unbiased and has lower expected mean squared error than that of the original GWAS summary statistics. Under the homogeneous-***ω*** assumption, MAMA is the best linear unbiased estimator (see Supplementary Note).

#### Estimating Ω_j_ and Σ_j_

Above, we assume that **Ω**_*j*_ and **Σ**_*j*_ are known. In practice, we estimate them using an approach similar to LD score regression (LDSC)^24^ and POPCORN^15^. From equation (1),

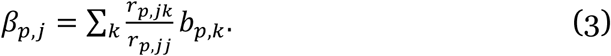

The *j*^th^ diagonal element of **Ω**_*j*_ is the variance of *β*_*p,j*_ : Ω_*pp,j*_ = ∑_*k*_ (*r*_*p,jk*_/*r*_*p,jj*_)^2^Var(*b*_*p,k*_). By the homogeneous-***ω*** assumption, *ω*_*pp*_ ≡ Var(*b*_*p,k*_). Thus, Ω_*pp,j*_ = *ω*_*pp*_*ℓ*_*j,pp*_, where *ℓ*_*j,pp*_ ≡ ∑_*k*_ (*r*_*p,jk*_/*r*_*p,jj*_)^2^ is closely related to the LD score used in LDSC.

The (*p, q*)^th^ off-diagonal entry of **Ω**_*j*_ is the covariance between *β*_*p,j*_ and *β*_*q,j*_ for populations *p* and *q*. Similarly as for the diagonal entries, Ω_*pq,j*_ = *ω*_*pq*_*ℓ*_*j,pq*_, where *ω*_*pq*_ ≡ Cov(*b*_*p,k*_, *b*_*q,k*_) is the covariance of conditional effect sizes between population *p* and *q* and *ℓ*_*j,pq*_ ≡ ∑_*k*_(*r*_*p,jk*_*r*_*q,jk*_)/(*r*_*p,jj*_ *r*_*q,jj*_) is a cross-ancestry generalization of an LD score. The LD score *ℓ*_*j,pq*_ is similar to that used in POPCORN^15^. Since the variance of the error, **Σ**_*j*_, is the residual variance in 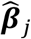 after accounting for **Ω**_*j*_, it can be estimated using the LDSC intercept.

Using these relationships, we estimate the elements of **Ω**_*j*_ and **Σ**_*j*_. First, we calculate LD scores *ℓ*_*j,pp*_ and cross-ancestry LD scores *ℓ*_*j,pq*_ using a reference panel. Then, we regress 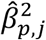 onto the vector of single-ancestry LD scores, *ℓ*_*j,pp*_, and we regress 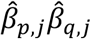 onto the vector of *ℓ*_*j,pq*_. The slope coefficients of these regressions are consistent estimates of *ω*_*pp*_ and *ω*_*pq*_, respectively, and the intercept is a consistent estimate of the corresponding element of **Σ**_*j*_. Using these estimates of *ℓ*_*j,pp,t*_, *ℓ*_*j,pq,t*_, *ω*_*pp*_, and *ω*_*pq*_, we construct the sample analog of **Ω**_*j*_, which is a consistent (method of moments) estimator for **Ω**_*j*_. Finally, we substitute our estimates of **Ω**_*j*_ and **Σ**_*j*_ into equation (1) to obtain MAMA estimates.

This procedure treats estimates of **Ω**_*j*_ and **Σ**_*j*_ as if they were estimated without error. This has two important implications. First, MAMA standard errors may be too small. Because the relevant aspects of MAMA are identical to those of MTAG, based on the simulations described in Turley et al.^16^, we anticipate this effect to be negligible as long as fewer than five populations are analyzed at once and as long as the reference panel used to construct LD scores is of sufficiently similar ancestry to the GWAS sample.

Second, when the GWAS for one or more populations is low powered, estimates of *ω*_*pq*_ may be very noisy, leading to bias and/or loss of statistical power. In such cases, it may increase precision to assume perfect genetic correlation between the populations (i.e., 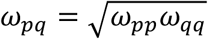.) When perfect genetic correlation is assumed, it is particularly important to verify the robustness of novel findings. Specifically, we recommend both replicating novel results and validating that the results are robust to assuming lower levels of genetic correlation.

#### LD Correlation

The LD scores used by MAMA can be combined to produce a metric of the similarity of LD patterns local to a certain SNP for a pair of populations. This metric, which we call the LD correlation, is

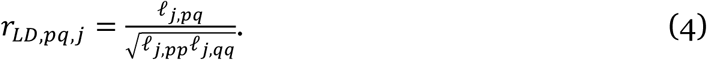

When *r*_*p,jk*_ = *r*_*q,jk*_ for all *k* for some SNP *j*, then *ℓ*_*j,pq*_ = *ℓ*_*j,pp*_ = *ℓ*_*j,qq*_ and therefore *r*_*LD,pq,j*_ = 1. Alternatively, if *r*_*p,j*(_ is uncorrelated with *r*_*q,j*(_ across SNPs *k* for some SNP *j*, then *r*_*LD,pq,j*_ = 0. As such, *r*_*LD,pq,j*_ may be thought of as a measure of the correlation of the LD patterns local to SNP *j*. As discussed below, the performance of MAMA and other cross-ancestry meta-analysis methods differs across SNPs with different LD correlations.

### Comparison to other methods and simulation

Several existing methods have been used to conduct GWAS meta-analysis across populations, including fixed-effect IVW meta-analysis (FE, e.g., ^17,18^), the modified random effects meta-analysis approach of Han and Eskin (RE2)^19^, Meta-Analysis of Transethnic Association studies (MANTRA)^20^, and Meta-Regression of Multi-Ethnic Genetic Association (MR-MEGA)^21^. These approaches rely on different assumptions and have varying computational intensity. For example, FE is computationally fast but assumes that the marginal effect of each SNP is the same across populations. RE2, MANTRA, and MR-MEGA model cross-population heterogeneity of marginal effects, but none incorporate information on LD differences. MANTRA is substantially more computationally intensive than any of these other methods, and MR-MEGA can only be run on three or more populations at once.

We conducted simulations to compare MAMA to (i) a standard single-ancestry GWAS, (ii) FE, (iii) MTAG, (iv) RE2, (v) MANTRA, and (vi) MR-MEGA. MTAG was not developed for cross-ancestry meta-analysis, but we include it because it accounts for heterogeneity across GWAS summary statistics.

We assessed three metrics of performance: bias, mean *χ*^2^ statistic, and type-1 error rate. To evaluate bias, we orient each SNP such that the true marginal effect is positive and report the mean difference between the estimated effect size and the true marginal effect. For the type-1 error rate, we report the fraction of SNPs with null marginal effects that have a *P* value of less than 0.05. MANTRA, which is a Bayesian method, does not report a standard error, so we cannot calculate a standard mean *χ*^2^ statistic or type-1 error rate. We therefore use the posterior standard deviation as an imperfect proxy for the standard error to calculate these performance metrics. Because the bias, mean *χ*^2^ statistic, and type-1 error rate are a function of the LD correlation, *r*_*LD,pq,j*_, we evaluate these performance metrics in five LD correlation bins.

We conducted two- and three-population simulations. The summary statistics in our simulations are based on estimated LD patterns of the AFR, EAS, and EUR subsamples from the 1000 Genome Project data^22^. We assess the bias and mean *χ*^2^ statistic of each method in data simulated under an infinitesimal genetic architecture that satisfies MAMA’s homogeneous-***ω*** assumption. We test the type-1 error rate with data simulated under a spike-and-slab model, where some SNPs are null in the EAS population but non-null in the other population(s). For the SNPs that are null only in the EAS population, this model violates the homogeneous-***ω*** assumption. We use these SNPs to assess the robustness of each method when this assumption is violated. For more details, see the Online Methods.

Figures 1-3 show the results of the three-population simulations for EAS; Supplementary Figures 1-3 show the full three-population results, and Supplementary Figures 4-6 show the results of the two-population simulations. Bias estimates for each method and LD correlation bin are in Figure 1. Across the three populations, only GWAS and MAMA have low bias for the entire LD correlation spectrum. MTAG estimates are biased at high and low LD correlation levels. All other methods have substantial bias toward zero, with the exception of MR-MEGA in the AFR population. These estimates have low bias because MR-MEGA puts little weight on the EAS and EUR summary statistics, so little bias can be introduced. MANTRA has the largest bias, but it is a Bayesian estimator so shrinkage toward zero is expected.

**Figure 1.**
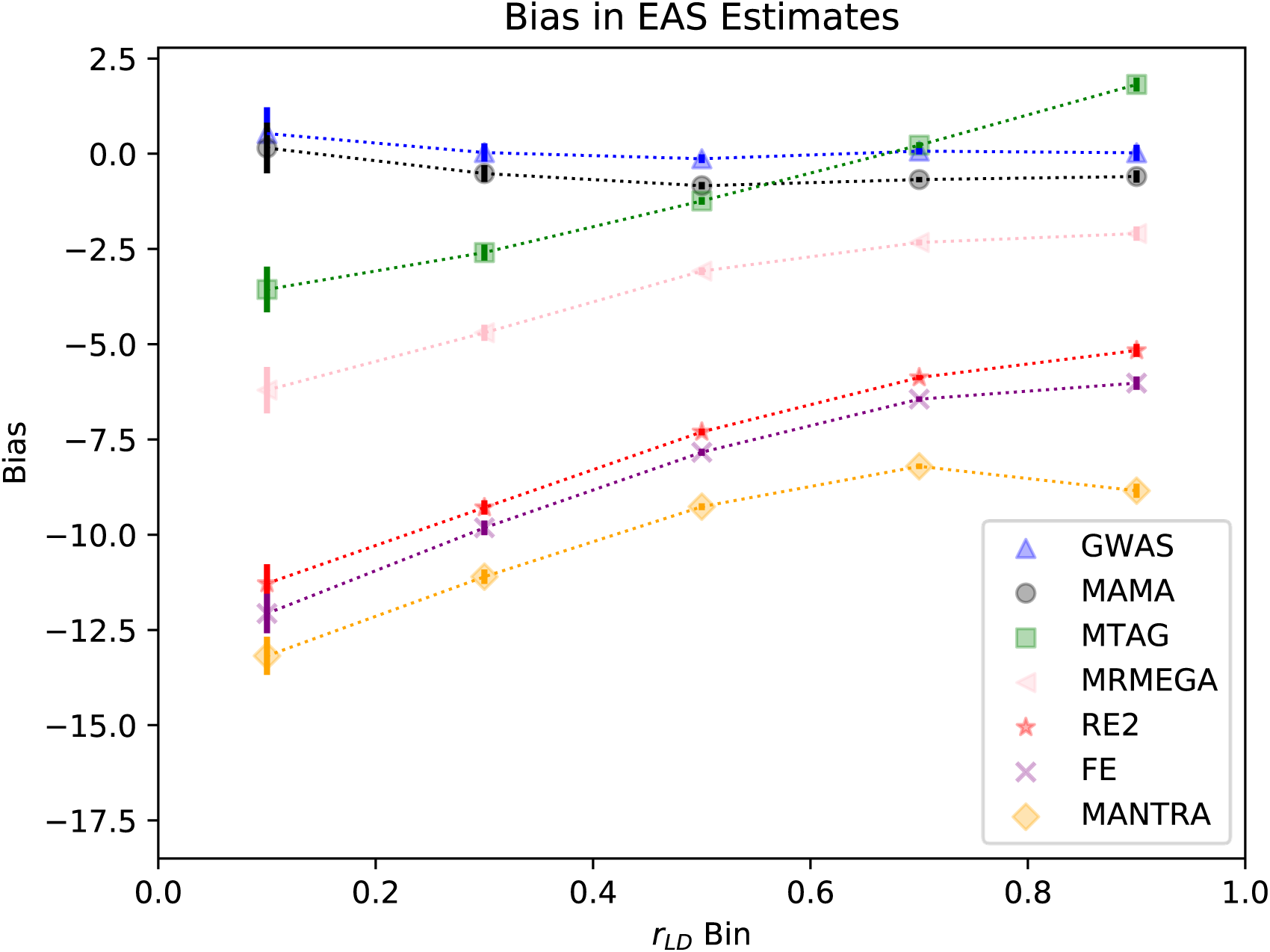
Bias for cross-ancestry meta-analysis methods by LD correlation bin. Note: Bias estimates in EAS defined as the difference between marginal SNP effects and meta-analyzed SNP effects averaged within LD correlation bins. All SNPs are oriented such that their marginal effects are positive. 95% confidence intervals are represented by vertical bars. In the three-population case, r_LD_ is calculated as the average r_LD_ value between the two population pairs that contain the target population (i.e., EAS-EUR and EAS-AFR for EAS). SNPs are binned into 5 groups of equal width between zero and one, and the bias is reported for SNPs within a bin.

**Figure 2.**
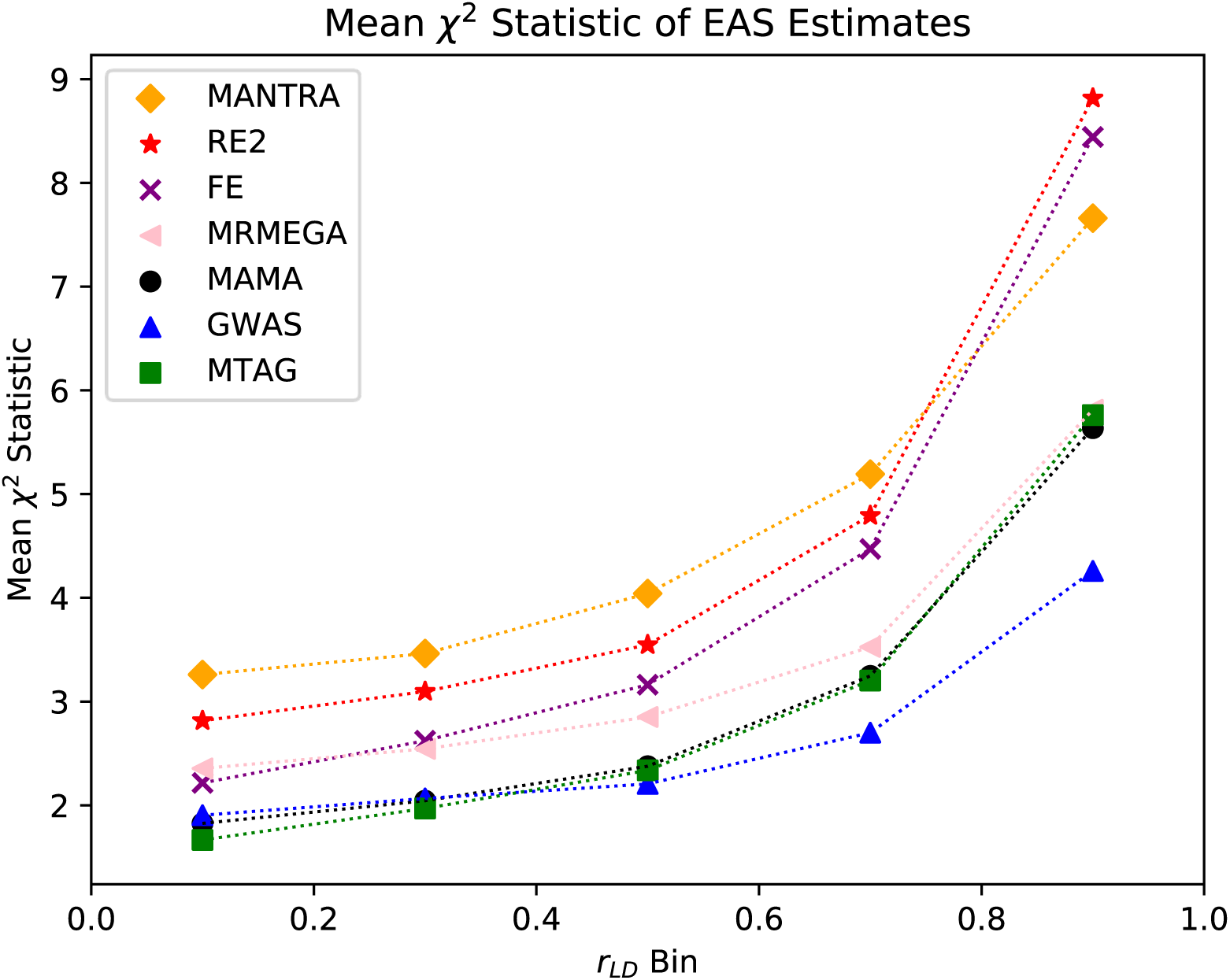
Mean *χ*^2^ statistic for cross-ancestry meta-analysis methods by LD correlation bin. Note: Mean ***χ***^**2**^ estimates in EAS defined as the squared ratio of the meta-analyzed SNP effect over the SNP’s standard error averaged within LD correlation bins. RE2 mean ***χ***^**2**^uses the reported RE2 P value and evaluates it on the inverse ***χ***^**2**^ distribution with one degree of freedom. In the three-population case, r_LD_ is calculated as the average r_LD_ value between the two population pairs that contain the target population (i.e., EAS-EUR and EAS-AFR for EAS). SNPs are binned into 5 groups of equal width between zero and one, and the mean ***χ***^**2**^ estimate is reported for SNPs within a bin.

**Figure 3.**
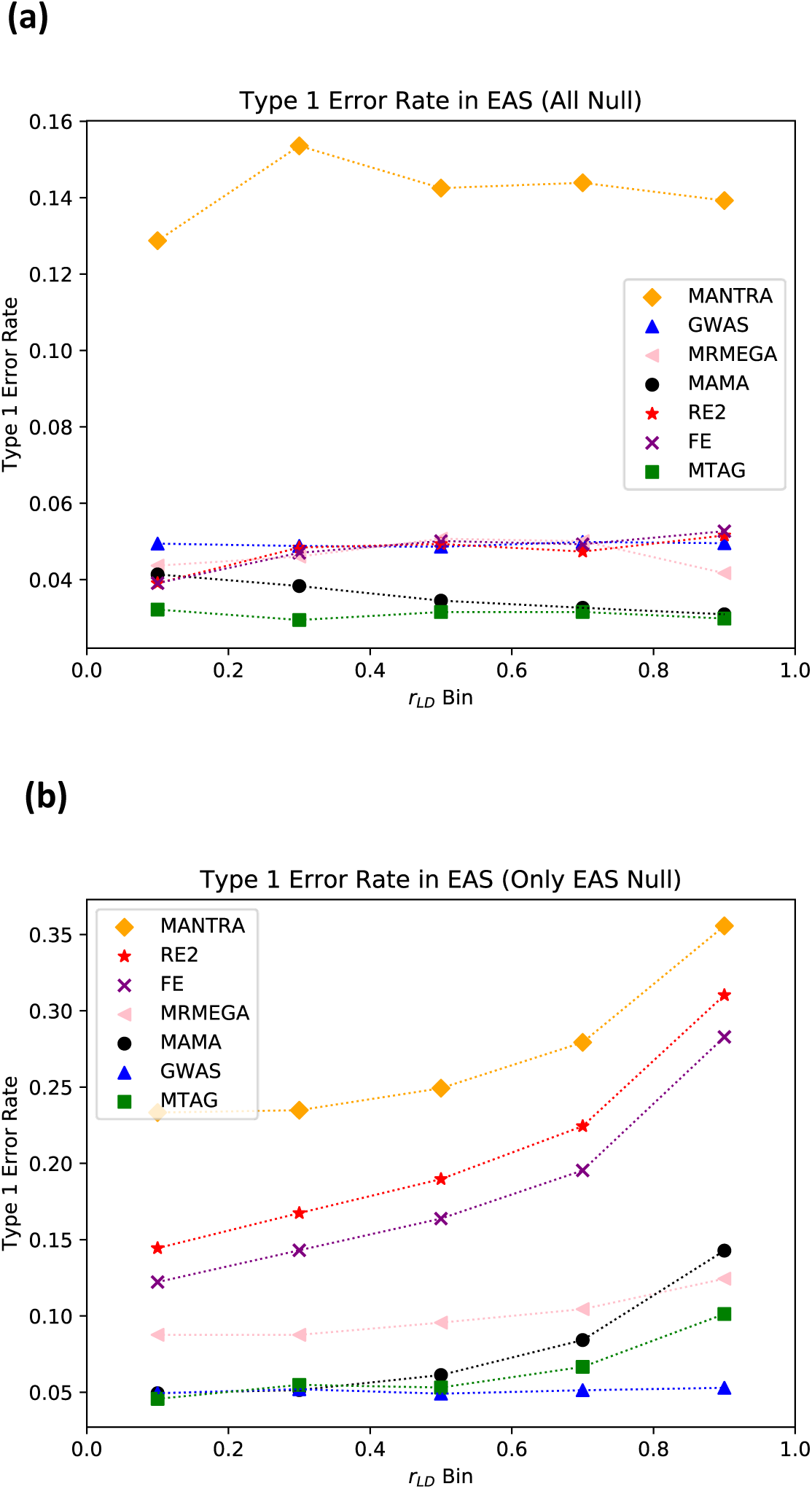
Type-1 error rate for cross-ancestry meta-analysis methods by LD correlation bin. Note: Type-1 error rate in (a) EAS for SNPs that are null in all three populations and (b) EAS for SNPs that are null in the EAS population but nonnull in the other populations. Type-1 error rate is defined as the fraction of null SNPs whose P value is less than 0.05. r_LD_ is calculated as the average r_LD_ value between the two population pairs that contain the target population (i.e., EAS-EUR and EAS-AFR for EAS). SNPs are binned into 5 groups of equal width between zero and one, and the type-1 error rate is reported for SNPs within a bin.

Figure 2 shows the mean *χ*^2^ statistic of each method. The patterns are the same for the EAS and EUR populations. In our simulations, MANTRA, RE2, FE, and MR-MEGA generally have a greater mean *χ*^2^ statistic than MAMA, and MTAG tends to have a comparable mean *χ*^2^ statistic. However, as shown in Figure 1, these gains in power come at the cost of substantial bias.

Figure 3a reports the type-1 error rate of SNPs that are null in all three populations. The type-1 error rate is controlled if it is less than 0.05. MANTRA is the only method that has an uncontrolled type-1 error rate, but this just means that the posterior standard deviation is not a good proxy for the standard error of MANTRA estimates. Both MAMA and MTAG have a type-1 error rate less than 0.05. That is because both methods include a stratification correction (even though there is no stratification in the simulation), which slightly inflates the standard errors.

Figure 3b reports the type-1 error rate of SNPs that were null in the EAS population but non-null in the AFR and EUR populations, a violation of the homogeneous-***ω*** assumption. For these SNPs, MAMA has an uncontrolled type-1 error rate, especially for SNPs with a high LD correlation. However, MANTRA, RE2, and FE each have even higher type-1 error rates. MR-MEGA has higher type-1 error rates than MAMA in all but the highest LD correlation bin. In this scenario, MTAG has lower type-1 error rate across most of the LD correlation spectrum.

In summary, although some methods have greater statistical power (as measured by the mean *χ*^2^ statistic) than MAMA, MAMA generally is the least biased across the LD correlation spectrum. Furthermore, although all methods considered except MANTRA control the type-1 error rate when the homogeneous-***ω*** assumption holds, MAMA is more robust to a plausible violation of this assumption than all other methods except MTAG.

### Application in Real Data

We applied MAMA to GWAS summary statistics for 23 phenotypes that were available from the China Kadoorie Biobank (CKB) and/or the Biobank Japan (BBJ) and for which we had access to comparable phenotypes in the UK Biobank (UKB). In the CKB and BBJ cohorts, we restricted our sample to those classified as having EAS ancestries; in the UKB cohort, we restricted our sample to those classified as having EUR ancestries. The complete list of phenotypes and cohorts we considered are in Supplementary Table 2. For each phenotype, each cohort is split into discovery and replication samples, and separate GWASs are run in each sample. Sample and SNP filters used by each cohort are in Supplementary Tables 3 and 4.

The LD score regression estimates, 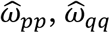, and 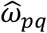 yield an estimator of the genetic correlation between populations *p* and 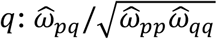. When heritability is low and sample size is small, the resulting estimate can be noisy and often nonsensical. For example, the estimated EAS-EUR genetic correlation for hematocrit is 3.4, and the average estimate across phenotypes is 1.4 (See Supplementary Table 1). Because the estimates are noisy (and not inconsistent with perfect genetic correlation), we impose perfect genetic correlation but allow for differences in heritability across populations (see Online Methods). Later, we explore the robustness of our results to alternative assumptions.

Before summarizing results across all phenotypes, we illustrate the performance of MAMA using two phenotypes: BMI, which has a smaller difference in sample sizes across populations, and educational attainment, which has a larger difference. For BMI, we use GWAS summary statistics from Biobank Japan and the China Kadoorie Biobank (combined *N* = 188,613) and the UK Biobank (*N* = 350,011). For educational attainment, we use summary statistics from the China Kadoorie Biobank (*N* = 42,435) and from the EUR-based GWAS meta-analysis reported in Lee et al.^25^ except omitting our UKB replication sample from the meta-analysis (combined *N* = 1,037,282). For brevity, we focus our discussion on the gains from MAMA in the EAS population; results for the EUR population are in the Supplementary Figures.

Figures 4-5 display Manhattan plots corresponding to the EAS GWAS and MAMA summary statistics. For both phenotypes, MAMA generates large increases in statistical power. In the GWAS of BMI in the EAS sample, we find 94 lead SNPs; using MAMA, we find 201. We categorize each MAMA lead SNP as (i) being the same as a genome-wide-significant SNP from any of the input GWAS, (ii) being in LD with a genome-wide-significant SNP from either of the input GWAS, or (iii) being (approximately) uncorrelated with any genome-wide significant SNP *in either input GWAS*. We refer to SNPs in group (iii) as “novel lead SNPs.” By this definition, of the 201 lead SNPs, 56 are novel. In order to obtain an increase in the mean *χ*^2^ statistic equivalent to that observed in the EAS MAMA summary statistics, we would have to increase the GWAS sample size from 188,613 to 282,048 individuals (see Online Methods). For educational attainment, where the sample-size difference is starker, the EAS GWAS contains no genome-wide significant SNPs, but the MAMA results identify 105 independent loci. Six of these loci are novel by our definition above. Attaining the same mean *χ*^2^ statistic in an EAS-ancestry GWAS as we observe in the MAMA results would require 542,873 individuals, compared to the 42,435 individuals available in this study.

**Figure 4.**
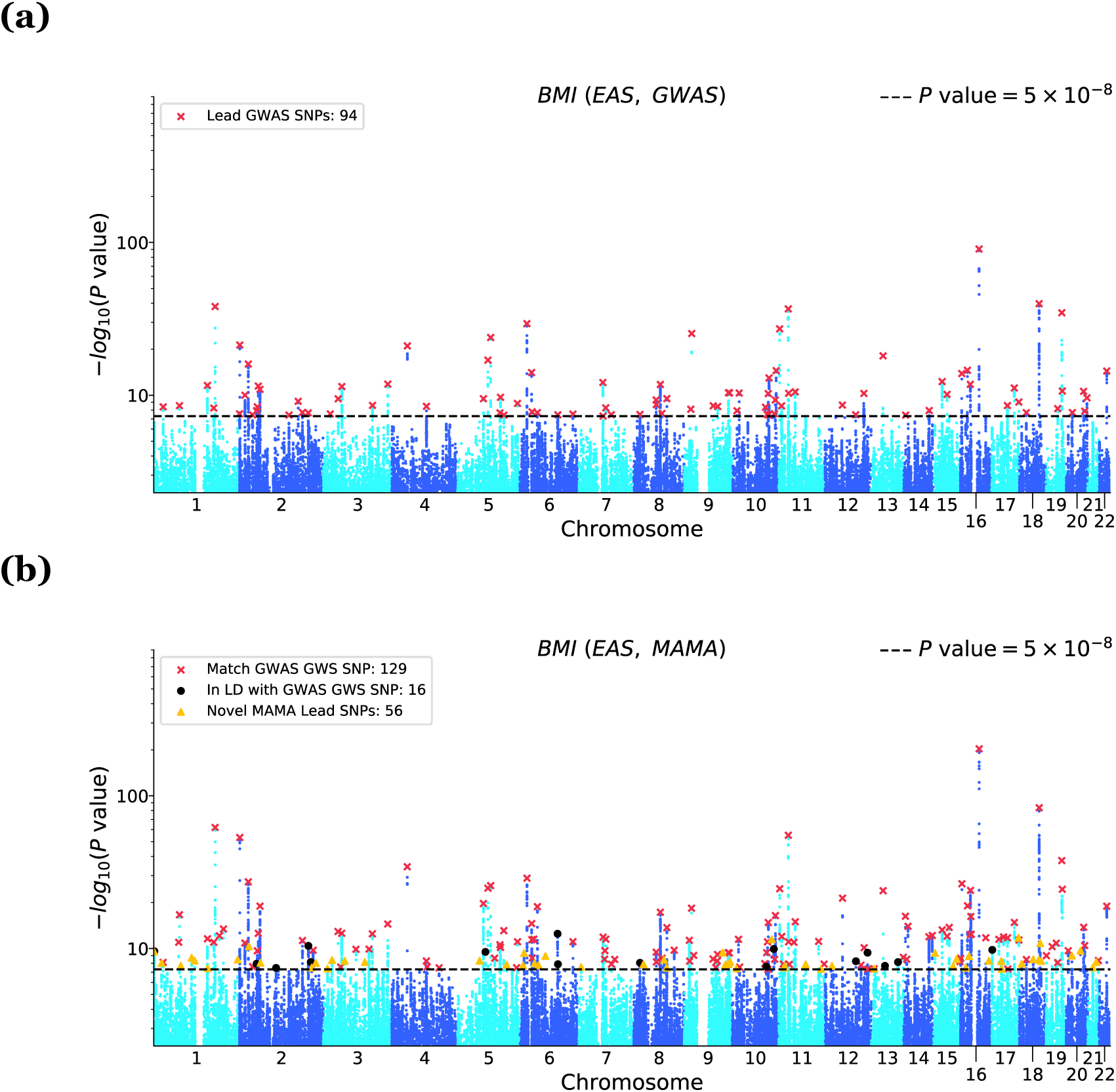
Manhattan plots for GWAS and MAMA results for BMI in an EAS population. Note: BMI for (a) EAS GWAS and (b) EAS MAMA. The x-axis is chromosomal position, and the y-axis is the P value on a -log10 scale (note the y-axes scale logarithmically). The dashed line marks the threshold for genome-wide significance (P = 5 × 10^−8^). For the GWAS Manhattan plots, lead SNPs (i.e., approximately independent SNPs surpassing the genome-wide-significance threshold) are marked with a red ×. For the MAMA Manhattan plots, lead SNPs are binned into one of three mutually exclusive categories: matching a GWAS lead SNP in either population (marked with a red ×), in LD with a GWAS lead SNP in either population (marked with a black •), or a novel SNP that is independent of any GWAS lead SNP in either population (marked with a yellow ▴). Details on lead SNP and novel SNP identification can be found in Online Methods.

**Figure 5.**
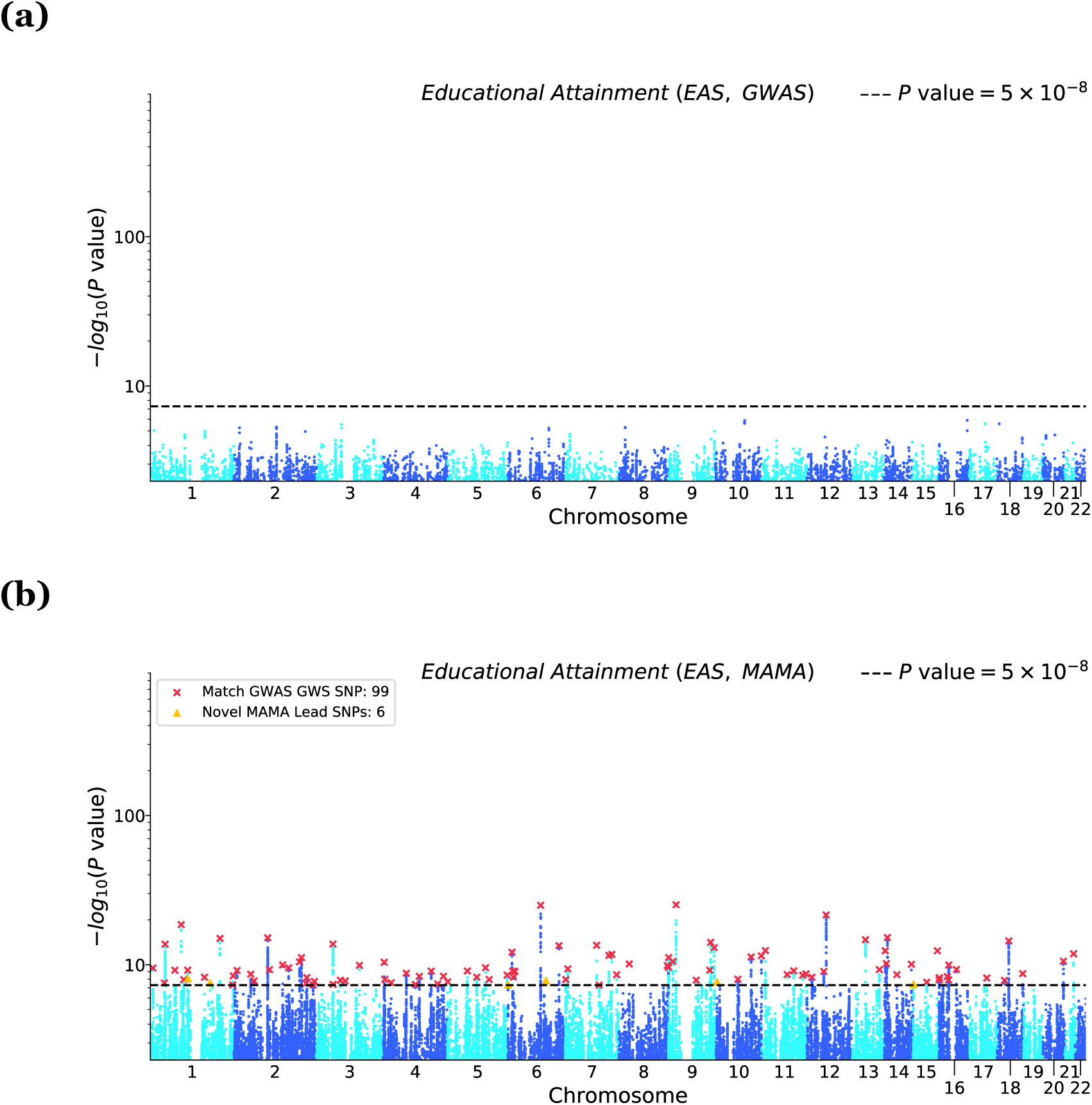
Manhattan plots for GWAS and MAMA results for Educational Attainment in an EAS population. Note: Educational Attainment for (a) EAS GWAS and (b) EAS MAMA. The x-axis is chromosomal position and the y-axis is the P value on a -log10 scale (note the y-axes scale logarithmically). The dashed line marks the threshold for genome-wide significance (P = 5 × 10^−8^). For the GWAS Manhattan plots, lead SNPs (i.e., approximately independent SNPs surpassing the genome-wide-significance threshold) are marked with a red ×. For the MAMA Manhattan plots, lead SNPs are binned into one of three mutually exclusive categories: matching a GWAS lead SNP in either population (marked with a red ×), in LD with a GWAS lead SNP in either population (marked with a black •), or a novel SNP that is independent of any GWAS lead SNP in either population (marked with a yellow ▴). Details on lead SNP and novel SNP identification can be found in Online Methods.

**Figure 7.**
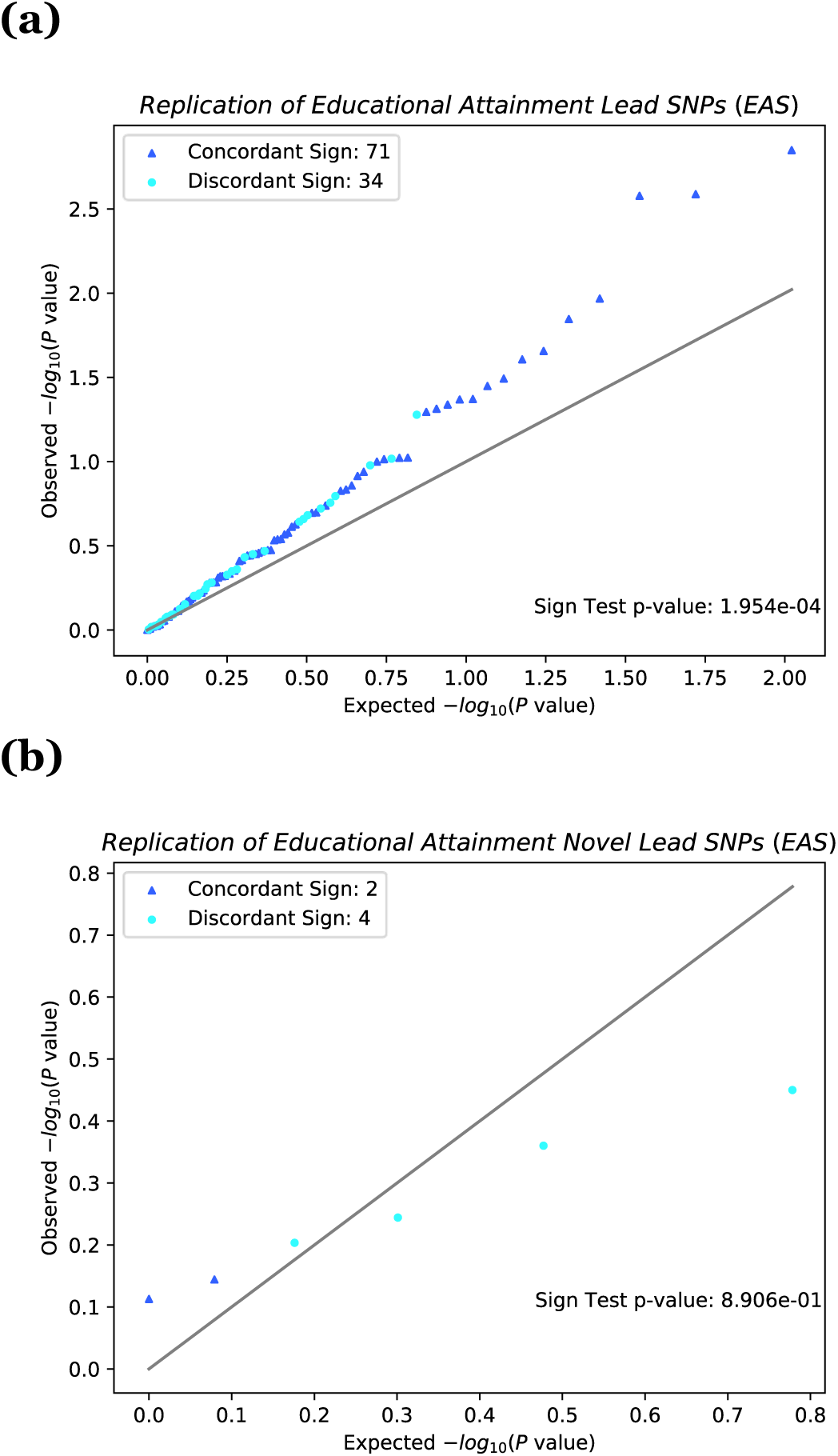
QQ plot of the lead SNPs and novel lead SNPs for Educational Attainment in an independent EAS population. Note: QQ plot and sign-test replication for Educational Attainment of (a) all lead SNPs in MAMA EAS and (b) all novel lead SNPs in MAMA EAS. The x-axis corresponds to the uniform distribution of P values expected under the null hypothesis, and the y-axis corresponds to the observed P values in the replication GWAS output. P values are reported on the -log10 scale. The 45-degree line is plotted for reference. The sign of each MAMA SNP is compared against an independent replication GWAS. SNPs whose sign matches the sign reported in the replication GWAS are marked with a dark blue ▴, and SNPs whose sign does not match the sign reported in the replication GWAS are marked with a light blue •. Sign test P values are drawn from the binomial distribution with success probability 1/2. Details on lead SNP and novel SNP identification can be found in Online Methods.

To address the concern that our assumption of perfect genetic correlation could lead to false discoveries, we validate the robustness of our results to alternate assumptions. When we assume a cross-ancestry genetic correlation of 0.9, we find that 179 out of 201 lead SNPs for BMI and 80 of 105 lead SNPs for educational attainment remain genome-wide significant. Out of the 56 novel lead SNPs for BMI and 6 for EA, 48 and 3 remain genome-wide significant, respectively.

We also address concerns about false discoveries with a replication analysis. In our independent EAS-ancestry holdout sample, MAMA-identified lead SNPs for BMI have inflated test statistics and have concordant signs with the MAMA associations more often than would be expected by chance if all MAMA lead SNPs were truly null. This is seen in Figures 5-6, which show QQ plots of the lead SNPs identified by MAMA but estimated in the EAS-ancestry replication sample. For BMI, 192 of the 201 lead SNPs have a concordant sign between the MAMA and replication samples (*P* ≤ 1.11 × 10^−16^). Among the novel lead SNPs (i.e., those not in LD with the lead SNPs identified in either the EAS or the EUR GWAS), 54 out of 56 SNPs have a concordant sign (*P* = 2.22 × 10^−14^; see Panel B). For educational attainment, 71 out of 105 lead SNPs have a concordant sign (*P* = 1.95 × 10^−4^). A test of concordant signs for the novel lead SNPs is not well powered because there are only 6. This low power is reflected in the sign-test *P* value of 0.891. Overall, however, these results, together with the robustness of lead SNPs at lower assumed levels of genetic correlation, lend credibility to the loci discovered by MAMA.

**Figure 6.**
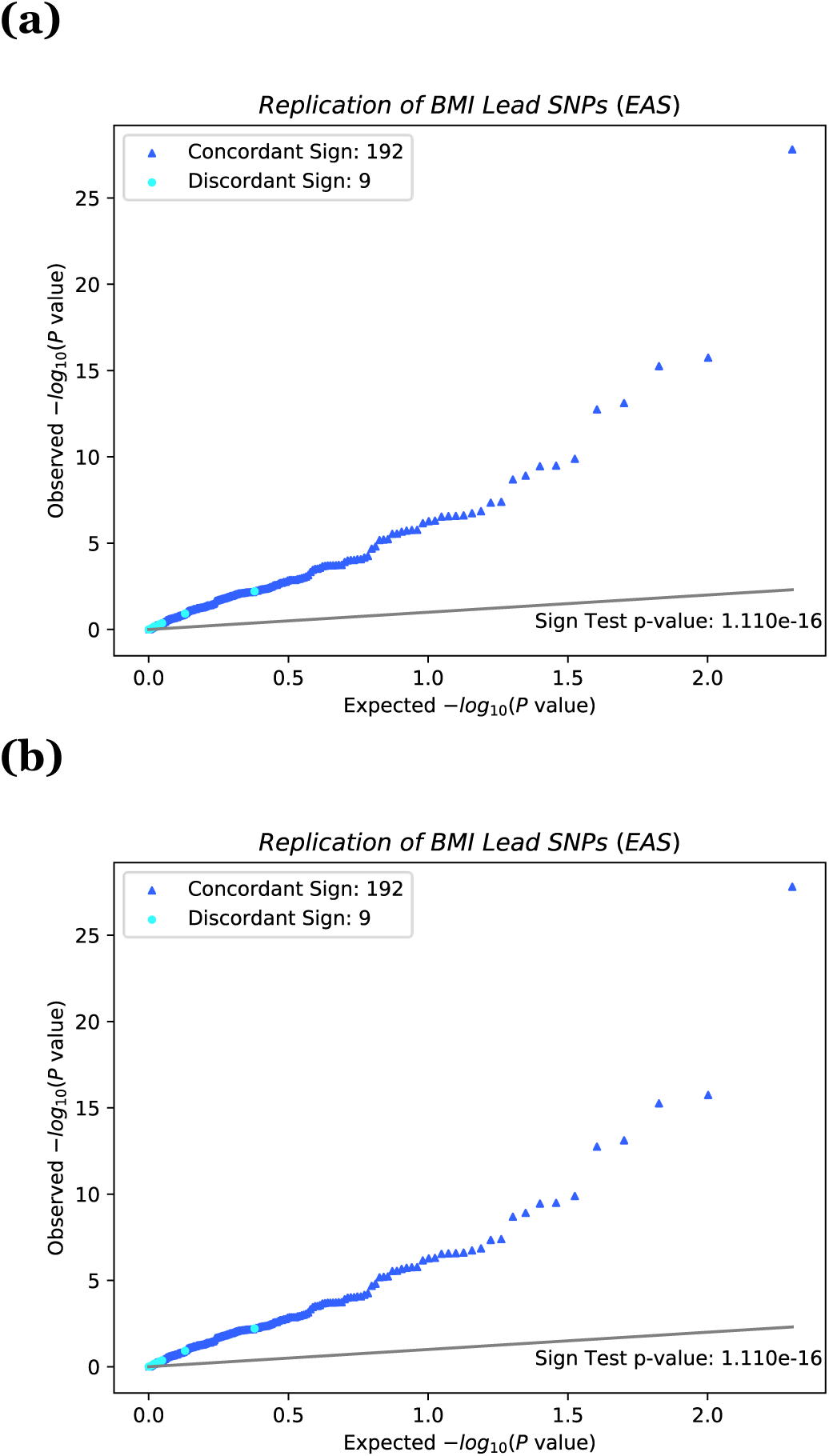
QQ plot of the lead SNPs and novel lead SNPs for BMI in an independent EAS population. Note: QQ plot and sign-test replication for BMI of (a) all lead SNPs in MAMA EAS and (b) all novel lead SNPs in MAMA EAS. The x-axis corresponds to the uniform distribution of P values expected under the null hypothesis, and the y-axis corresponds to the observed P values in the replication GWAS output. P values are reported on the - log10 scale. The 45-degree line is plotted for reference. The sign of each MAMA SNP is compared against an independent replication GWAS. SNPs whose sign matches the sign reported in the replication GWAS are marked with a dark blue ▴, and SNPs whose sign does not match the sign reported in the replication GWAS are marked with a light blue •. Sign test P values are drawn from the binomial distribution with success probability 1/2. Details on lead SNP and novel SNP identification can be found in Online Methods.

Table 1 contains a summary of results for all phenotypes tested. The patterns observed for the two example phenotypes are also present overall. The average increase in effective sample size of the MAMA summary statistics in the EAS population is 183,303 individuals. Across all phenotypes we find 3,257 lead SNPs for the EAS population, 359 of which are novel (i.e., are not identified in the GWAS results for either the EAS or EUR population). Of these 359 novel SNPs, 303 remain genome-wide-significant assuming a genetic correlation of 0.9 instead of 1 (See Supplementary Table 5). Finally, in a series of 42 sign tests (21 phenotypes each with 2 ancestries), the novel MAMA loci replicate with *P* values less than the Bonferroni-adjusted threshold of 0.05/42 in 38 cases. Overall, these results provide reassuring evidence that MAMA identifies robust associations.

## DISCUSSION

Substantial undersampling of non-European-ancestry individuals has resulted in disparities in the value of the genetics research conducted during the GWAS era. There is broad agreement that more genome-wide data should be gathered from underrepresented populations. To complements these data-gathering efforts, here we developed a method for cross-ancestry meta-analysis, MAMA, that efficiently leverages *existing* European-ancestry GWAS summary statistics to add information to GWAS summary statistics from other ancestry populations.

Like all tools, MAMA has strengths and limitations. We emphasize four limitations and interpretational caveats.

First, MAMA may produce spurious results if some SNPs affect the phenotype of interest in one population but not the other. That said, our simulation results suggest that MAMA is more robust (with respect to the Type-1 error rate) for such SNPs than other methods we considered.

Second, if the empirical estimates of ***Ω***_*j*_ are biased or imprecise, MAMA will generally have lower power to detect associations, and MAMA estimates may be biased.

There are two primary reasons estimates of ***Ω***_*j*_ may be poor: (i) the reference sample is too small or is not representative of the corresponding GWAS sample or (ii) the GWAS sample size is too small. Reason (i) will lead to biased estimates of ***Ω***_*j*_. Therefore, we recommend limiting MAMA analyses to summary statistics from populations with a large, representative reference sample available. Reason (ii) will lead to imprecise estimates of ***Ω***_*j*_. Therefore, if a large GWAS sample is not available, we recommend that users fix the genetic correlation between populations to 1 but check which novel loci remain significant when lower genetic correlation values are assumed.

Third, the gains from MAMA are not uniform genome-wide; increases in power will be larger for SNPs that have more similar LD structure between populations^26^. As a result, MAMA is more likely to miss genetic signal that is concentrated in regions of high LD variability across populations.

Finally, although a polygenic predictor based on MAMA-generated weights for population *j* should generally outperform a polygenic predictor derived from population *j*’s GWAS summary statistics alone, MAMA is not designed as a prediction tool. The unbiasedness of MAMA estimates is a strength for gene discovery, but it is a limitation for prediction. Methods that generate biased estimates but less sampling variance— including standard, inverse-variance-weighted meta-analysis of cross-ancestry GWAS summary statistics, not taking into account LD and other population differences—can outperform MAMA in prediction accuracy. Exploring optimal ways to use GWAS summary statistics from European-ancestry populations to improve polygenic prediction in non-European-ancestry populations is an active and important area of research^27,28^.

We have shown that MAMA can be a useful tool for multi-ancestry genetics research. In our simulations, MAMA did not always yield the greatest gains in reported statistical power, but we consistently found that MAMA estimates were less biased than other methods and more robust to violations of key assumptions. We anticipate that MAMA estimates will prove especially valuable when it is important to have approximately unbiased estimates of SNP associations. For example, many methods for fine-mapping^29^, partitioning heritability^30,31^, and biological annotation^32,33^ depend on reliable estimates of effect sizes.

## Supporting information

Supplementary Note and FAQ

## DATA AVAILABILITY

For each phenotype that we analyze, we report GWAS and MAMA summary statistics. With the exception of the summary statistics for educational attainment (which include data from 23andMe), complete summary statistics for both the EAS and EUR samples will be available for download at the SSGAC website (http://www.thessgac.org/data) upon publication. SNP-level summary statistics from analyses based entirely or in part on 23andMe data can only be reported for up to 10,000 SNPs. Therefore, for this phenotype, we provide summary statistics for only the genome-wide-significant SNPs from that analysis. In addition, we provide complete summary statistics for an analysis that omits 23andMe. The full GWAS summary statistics for the 23andMe discovery data set will be made available through 23andMe to qualified researchers under an agreement with 23andMe that protects the privacy of the 23andMe participants. Please visit https://research.23andme.com/collaborate/#dataset-access/ for more information and to apply to access the data.

## CODE AVAILABILITY

The MAMA software is available at https://github.com/JonJala/mama.

## ACKNOWLEDGEMENTS

This research was carried out with the support of the Social Science Genetic Association Consortium (SSGAC). This research was conducted using the UK Biobank Resource under application numbers 11425 and 31063. The study was supported by funding from the Ragnar Söderberg Foundation (E42/15, D.C.), the Swedish Research Council (2019-00244, D.C., PI: Sven Oskarsson), Open Philanthropy (010623-00001, D.J.B. and M.N.M.), Riksbankens Jubileumsfond P18-0782:1 (D.C., PI: S.O.), Pershing Square Fund for Research on the Foundations of Human Behavior (D.L.), and the NIA/NIH/NIMH through grants R24-AG065184 (D.J.B.) to the University of California Los Angeles; K99-AG062787-01 (P.T.), K99/R00MH117229 (A.R.M.) and K01-MH121659 (E.G.A.) to Massachusetts General Hospital; 1R01-MH101244-02 (B.M.N.), 5U01-MH109539-02 (B.M.N.), and 5 R37 MH107649 (B.M.N.) to the Broad Institute at Harvard and MIT. We thank the following biobanks and consortia for sharing GWAS summary statistics: BBJ, CKB, SSGAC (educational attainment), and the Psychiatric Genomics Consortium (PGC, schizophrenia). We thank the research participants and employees of 23andMe for making this work possible. A full list of acknowledgements is provided in the Supplementary Note.

China Kadoorie Biobank acknowledges the contribution of participants, project staff, and the China National Centre for Disease Control and Prevention (CDC) and its regional offices. China Kadoorie Biobank was supported as follows: Baseline survey and first re-survey: Hong Kong Kadoorie Charitable Foundation; long-term follow-up and second re-survey: UK Wellcome Trust (212946/Z/18/Z, 202922/Z/16/Z, 104085/Z/14/Z, 088158/Z/09/Z), National Natural Science Foundation of China (91846303), and National Key Research and Development Program of China (2016YFC 0900500, 0900501, 0900504, 1303904). DNA extraction and genotyping: GlaxoSmithKline, UK Medical Research Council (MC_PC_13049, MC-PC-14135). The UK Medical Research Council (MC_UU_00017/1, MC_UU_12026/2 MC_U137686851), Cancer Research UK (C16077/A29186; C500/A16896) and the British Heart Foundation (CH/1996001/9454), provide core funding to the Clinical Trial Service Unit and Epidemiological Studies Unit at Oxford University for the project.

The Biobank Japan Project was supported by the Tailor-Made Medical Treatment program (the BioBank Japan Project) of the Ministry of Education, Culture, Sports, Science, and Technology (MEXT), and the Japan Agency for Medical Research and Development (AMED).

## AUTHOR CONTRIBUTIONS

P.T., A.R.M., D.J.B. and B.M.N. oversaw the study. The theory underlying MAMA was conceived of and developed by P.T. and A.R.M., with contributions from B.M.N., D.J.B., D.C., and R.K.W. G.G., H.L. and J.B.J. developed the MAMA software. G.G. and H.L. were the primary analysts for the empirical applications in this paper and contributed equally to this work. M.K., R.K.W., C.C., D.P., M.Z., E.G.A., R.G.W, I.Y.M, K.L, Z.C, and L.L also made contributions to the empirical work. P.T., A.R.M., and D.J.B. coordinated the writing of the manuscript. S.C. and M.M. provided feedback on the ethical implications of the research. S.C. advised on the characterization of populations used in this work, emphasizing the importance of understanding genomic diversity within continental groups. P.T., A.R.M., S.C., D.L., M.M, and D.B. wrote the FAQs. G.G., M.K., and R.W. also contributed to the writing. All authors provided input and revisions for the final manuscript.

## TABLES

**Table 1. Summary of MAMA results of 23 phenotypes.**

## Online Methods

This article is accompanied by a Supplementary Note with further details.

## Data Generating Procedure for Simulations

In our simulations, we generate GWAS summary statistics for hypothetical AFR, EAS, and EUR populations. For the infinitesimal model used to measure bias and the mean *χ*^2^ statistic, we generate true marginal effect sizes with the following approach. First, for the corresponding subsample of the 1000 Genomes Project data^22^ (consisting of all individuals with a genotype missingness rate no greater than 2%), we estimate the LD between each pair of SNPs that have an allele frequency greater than 1% and a missingness rate no more than 2% within their subsample in every population. We fix the LD between a pair of SNPs to zero if they are farther than 1 centimorgan apart or if they are on different chromosomes. We then draw true conditional effect sizes for each standardized SNP in these samples from a standard normal distribution. We assume that the cross-ancestry genetic correlation is 1 for each pair of populations, which means that the conditional effect of each standardized SNP is the same in each population. True marginal effect sizes for each population are calculated using equation (3). For computational reasons, we retain only every 100^th^ SNP for the subsequent steps of the simulation, resulting in 60,107 SNPs.

For the spike-and-slab model used to measure type-1 error rate, we generate true marginal effect sizes using the same procedure as above with an additional step at the end. Specifically, we take the 60,107 SNPs for which we have marginal effects and we randomly set 10% of the marginal effects to zero in all three populations. We then randomly set the marginal effect to zero in just the EAS population for a random 10% set of SNPs that do not overlap with the 10% that have a zero marginal effect in all three populations. The EAS population was chosen arbitrarily in this simulation. By symmetry, we would anticipate comparable results no matter which population we chose to be null. Thus, 80% of SNPs have a non-zero marginal effect in all three populations, 10% of SNPs have a non-zero marginal effect in just the EUR and AFR populations, and 10% of SNPs have a zero marginal effect in all three populations.

Finally, in all simulations, we construct standardized GWAS effect-size estimates by adding mean-zero, normally distributed noise with variance 100 to each true marginal effect. Thus, the standard error for each SNP is 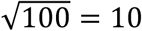. For the infinitesimal-model summary statistics, this results in a mean *χ*^2^ statistics of 2.271, 2.576, and 2.666 in the AFR, EAS, and EUR-based summary statistics, respectively. In the spike-and-slab model, the corresponding mean *χ*^2^ statistics are 2.144, 2.287, and 2.517, respectively. Next, we divide the true standardized marginal effect size, the standardized GWAS estimates, and standard errors by 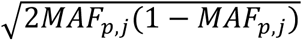, where *MAF*_*p,j*_ is the allele frequency of SNP *j* in population *p*, to transform these quantities into allele-count units rather than standardized units. This is done because published summary statistics are reported in allele-count units, and these are the units required for summary statistics provided to MAMA.

In simulations with just two populations, only the EAS and EUR summary statistics are retained, corresponding to the ancestries of the samples used in our empirical applications in this paper.

## Biobank Japan

The BioBank Japan Project (https://biobankjp.org/english/index.html) is a national hospital-based biobank started in 2003. The BBJ collected DNA, serum and clinical information of approximately 200,000 patients with any of 47 target diseases between fiscal years of 2003 and 2007 from 66 hospitals throughout Japan. Details of study design, sample collection, baseline clinical information, genotyping, and imputation are described elsewhere^34–36^. We retrieved individual phenotypes from medical records and applied standard quality control criteria followed by inverse-rank normalization as described previously^37^. We restricted our analysis to unrelated individuals and held out a random sample of 5,000 individuals as a replication sample. GWAS was conducted using Hail v0.2^38^ under a linear regression model with covariates including age, age^2^, sex, age × sex, age^2^ × sex, and the top 20 PCs.

## China Kadoorie Biobank

China Kadoorie Biobank (CKB, https://www.ckbiobank.org) is a population-based prospective cohort of approximately 513,000 adults aged 30-79 recruited in 2004-8 from 10 geographically defined regions of China^39^. Questionnaire and physical measurement data and biological samples were collected at baseline and at periodic re-surveys of 25,000 randomly selected surviving participants. Through linkages to death and disease registries and to health insurance databases, all participants are followed for cause-specific mortality and morbidity and for any hospital admission. Local, national and international ethics approval was obtained, and all participants provided written informed consent. Genotyping was conducted on custom Affymetrix Axiom^®^ arrays, with 100,706 unique samples and 511,885 variants passing QC. Genotypes were phased using SHAPEIT3 v4.12 and imputed into the 1000 Genomes Phase 3 reference with IMPUTE4 v4.r265. GWAS was conducted stratified by both sex and the 10 recruitment regions, using BOLT-LMM v2.3.2 with age and age^2^ as covariates. Analyses using CKB data were conducted under research approval 2018-0087.

## UK Biobank

The UK Biobank Project enrolled 500,000 people aged between 40-69 years in 2006-2010 from across the country, as described previously^40^. Details of the study design, data access, and other relevant information are described elsewhere (https://www.ukbiobank.ac.uk/). We conducted GWAS as described previously (http://www.nealelab.is/uk-biobank/ukbround2announcement) using a linear regression model with the same covariates as described in the BBJ analyses with one modification. We created a holdout validation set consisting of 10,000 individuals from the original GWAS analyses, then included the remaining individuals in GWAS, totaling a maximum of 351,194 individuals (in contrast, the original GWAS conducted by the Neale Lab had a maximum of *N*= 361,194 individuals).

## Construction of LD scores

LD scores used in these analyses were constructed using the 1000 Genomes Project Phase 3 samples. The LD scores were constructed using the –std-geno-ldsc units option, which assumes that the genotypes in the model are in allele-count units (see Supplementary Materials Section 1.2). We used the EUR samples (489 individuals) to construct LD scores for the European-ancestry-population summary statistics and used the EAS samples (481 individuals) to construct the LD scores for the East-Asian-ancestry-population summary statistics. Before making the score, we restricted the SNPs to just those with a minor allele frequency greater than 1% in either the EAS or EUR samples. This resulted in 10,294,523 SNPs in total (7,627,153 SNPs in the EAS population, 8,676,825 SNPs in the EUR population, and 6,009,455 SNPs in the intersection). The distributions of the LD scores and of the LD score correlations are in Supplementary Figure 50.

## MAMA settings in empirical applications

The MAMA analyses in this paper are conducted using the following settings. We assume that the phenotype has a genetic correlation of 1 between populations but that the heritability may differ between populations. We also assume that the model is in standardized units. This assumption implies that the conditional effect of a one-standard deviation change in genotype is independently and identically distributed across the genome. This assumption also implies that the expected magnitude of effect sizes (in allele count units) is larger for SNPs that are rarer. Finally, because there is little to no variation in sample size across SNPs within each population’s summary statistics in our data, there is also little variation in the standard errors. Thus, the intercept and squared standard error term are highly collinear in the LD score regression step of MAMA. To address this issue, we fix the intercept to zero and freely estimate the coefficient on the squared standard errors.

## GWAS-equivalent sample size

As one of the metrics of performance, MAMA reports the “GWAS-equivalent sample size.” It is parallel to the identically named metric used by MTAG^16^. This metric quantifies how large a GWAS sample you would need to observe the same gains in the mean *χ*^2^ statistic in MAMA relative to the original GWAS summary statistics. Because one minus the mean *χ*^2^ statistic scales linearly with sample size in expectation, the GWAS-equivalent sample size for MAMA is defined as

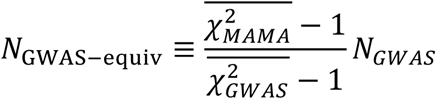

where 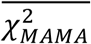 is the mean *χ*^2^ statistic of the MAMA summary statistics, 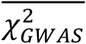 is the mean *χ*^2^ statistic of the GWAS summary statistics, and *N*_H*I*GJ_ is sample size of the GWAS. Because MAMA standard errors are implicitly corrected for population stratification using the LD score intercept, to make the mean *χ*^2^ statistic comparable between the GWAS and MAMA summary statistics, the GWAS standard errors were corrected using the LD score intercept before calculating 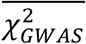. One difference between the GWAS-equivalent sample size in MAMA relative to MTAG is that the amount of weight put on the alternate set of summary statistics varies by SNP in MAMA, whereas it is constant in MTAG. As a result, the GWAS-equivalent sample size does not represent the sample size needed to obtain a certain level of power for any individual SNP, but instead it represents how large a sample would be needed to obtain an average amount of power genome-wide. For MAMA, the SNP-level effective sample size will be larger when the LD correlation is higher between the pairs of populations (see Supplementary Materials Section 1.5.1).

## Identifying lead SNPs

Using the discovery GWAS and MAMA results for each phenotype, we identify a set of genome-wide-significant lead SNPs. If there are no SNPs with a *P* value less than 5 ’ 10^−8^, then there are no genome-wide significant lead SNPs for that set of summary statistics. Otherwise, we use the following algorithm to identify lead SNPs. (1) The SNP with the smallest *P* value is set aside as a potential lead SNP. (2) All SNPs that are on the same chromosome as the SNP identified in (1) and that are correlated with the SNP at *R*^2^ > 0.1 are removed from the summary statistics. (3) Steps (1) and (2) are repeated until no SNP remains with a *P* value less than 5 × 10^−8^. (4) Among the list of potential lead SNPs, the SNP with the lowest *P* value is added to the final list of lead SNPs. (5) Any potential lead SNP within 500 kbp of the lead SNP identified in (4) is removed. (6) Steps (4) and (5) are repeated until no SNPs remain among the potential lead SNPs.

These steps are executed using two rounds of Plink’s clumping algorithm. *R*^2^ values were calculated using the 1000 Genomes Project reference panel sample corresponding to the population of the GWAS or MAMA summary statistics.

## Identifying Novel SNPs

To assess whether the lead SNPs identified by MAMA are novel relative to the input GWAS summary statistics, we first remove any of the SNPs from the MAMA lead-SNP list that have the same rsID as any genome-wide-significant SNPs from the GWAS summary statistics used to generate the MAMA summary statistics. We then combine the reduced set of MAMA lead SNPs with the lead SNPs from the input GWASs, adjusting the *P* values of the lead SNPs from the GWASs so they are smaller than the minimum *P* value in the MAMA lead-SNP list. We then conduct the two-stage clumping algorithm described in the Methods section “Identifying lead SNPs” to this combined set of SNPs. Any of the SNPs that are retained from the reduced set of MAMA lead SNPs after clumping are considered “novel lead SNPs.” Note that “novel” in this case does not imply that the SNPs are novel relative to the literature but rather that the SNPs are novel relative to the summary statistics used in the analysis.

## Replication

To validate the lead SNPs and the novel lead SNPs identified by MAMA, we look up these SNPs in GWAS in an independent sample from the same population. We then compare the sign of the novel lead SNPs from MAMA to the signs of the GWAS in the replication sample corresponding to the same population and phenotype. Under the null hypothesis that the novel lead SNPs from MAMA are null in the replication sample, the number of times the MAMA estimates have a concordant sign with the replication GWAS estimates will have a Binomial(*M*,0.5) distribution, where *M* is the number of novel lead SNPs. We perform a one-sided test measuring whether the sign concordance is higher than expected given the null hypothesis. We did not perform a replication analysis for Physical Activity because no lead SNPs were found, nor for Schizophrenia because we did not have access to a replication sample.

## REFERENCES

1. Visscher, P. M. et al. 10 Years of GWAS Discovery: Biology, Function, and Translation. Am. J. Hum. Genet. 101, 5–22 (2017).

2. Willer, C. J. et al. Six new loci associated with body mass index highlight a neuronal influence on body weight regulation. Nat. Genet. 41, 25 (2009).

3. Stein, J. L. et al. Identification of common variants associated with human hippocampal and intracranial volumes. Nat. Genet. 44, 552–561 (2012).

4. Yengo, L. et al. Meta-analysis of genome-wide association studies for height and body mass index in ∼700000 individuals of European ancestry. Hum. Mol. Genet. 27, 3641–3649 (2018).

5. King, E. A., Wade Davis, J. & Degner, J. F. Are drug targets with genetic support twice as likely to be approved? Revised estimates of the impact of genetic support for drug mechanisms on the probability of drug approval. PLoS Genet. 15, e1008489 (2019).

6. Nelson, M. R. et al. The support of human genetic evidence for approved drug indications. Nat. Genet. 47, 856–860 (2015).

7. Khera, A. V. et al. Genome-wide polygenic scores for common diseases identify individuals with risk equivalent to monogenic mutations. Nat. Genet. 50, 1219–1224 (2018).

8. Bonham, V. L., Green, E. D. & Pérez-Stable, E. J. Examining how race, ethnicity, and ancestry data are used in biomedical research. Jama 320, 1533–1534 (2018).

9. Popejoy, A. B. et al. The clinical imperative for inclusivity: race, ethnicity, and ancestry (REA) in genomics. Hum. Mutat. 39, 1713–1720 (2018).

10. Hindorff, L. A. et al. Prioritizing diversity in human genomics research. Nat. Rev. Genet. 19, 175 (2018).

11. Abul-Husn, N. S. & Kenny, E. E. Personalized medicine and the power of electronic health records. Cell 177, 58–69 (2019).

12. Investigators, A. of U. R. P. The “All of Us” Research Program. N. Engl. J. Med. 381, 668–676 (2019).

13. Bentley, A. R., Callier, S. L. & Rotimi, C. N. Evaluating the promise of inclusion of African ancestry populations in genomics. NPJ Genomic Med. 5, 1–9 (2020).

14. Bentley, A. R., Callier, S. & Rotimi, C. The emergence of genomic research in Africa and new frameworks for equity in biomedical research. Ethn. Dis. 29, 179 (2019).

15. Brown, B. C., Ye, C. J., Price, A. L., Zaitlen, N. & Consortium, A. G. E. N. T. 2 D. Transethnic genetic-correlation estimates from summary statistics. Am. J. Hum. Genet. 99, 76–88 (2016).

16. Turley, P. et al. Multi-trait analysis of genome-wide association summary statistics using MTAG. Nat. Genet. 50, 229–237 (2018).

17. Bigdeli, T. B. et al. Contributions of common genetic variants to risk of schizophrenia among individuals of African and Latino ancestry. Mol. Psychiatry 1–13 (2019).

18. Willer, C. J., Li, Y. & Abecasis, G. R. METAL: fast and efficient meta-analysis of genomewide association scans. Bioinformatics 26, 2190–2191 (2010).

19. Han, B. & Eskin, E. Random-effects model aimed at discovering associations in meta-analysis of genome-wide association studies. Am. J. Hum. Genet. 88, 586–598 (2011).

20. Morris, A. P. Transethnic meta-analysis of genomewide association studies. Genet. Epidemiol. 35, 809–822 (2011).

21. Mägi, R. et al. Trans-ethnic meta-regression of genome-wide association studies accounting for ancestry increases power for discovery and improves fine-mapping resolution. Hum. Mol. Genet. 26, 3639–3650 (2017).

22. Auton, A. et al. A global reference for human genetic variation. Nature 526, 68–74 (2015).

23. Hansen, L. P. Large sample properties of generalised method of moments estimators. Econometrica 50, 1029–1054 (1982).

24. Bulik-Sullivan, B. K. et al. LD Score regression distinguishes confounding from polygenicity in genome-wide association studies. Nat. Genet. 47, 291–295 (2015).

25. Lee, J. J. et al. Gene discovery and polygenic prediction from a genome-wide association study of educational attainment in 1.1 million individuals. Nat. Genet. 50, 1112–1121 (2018).

26. Marigorta, U. M. & Navarro, A. High trans-ethnic replicability of GWAS results implies common causal variants. PLoS Genet 9, e1003566 (2013).

27. Cai, M. et al. A unified framework for cross-population trait prediction by leveraging the genetic correlation of polygenic traits. Am. J. Hum. Genet. (2021).

28. Ruan, Y. et al. Improving Polygenic Prediction in Ancestrally Diverse Populations. medRxiv 2020.12.27.20248738 (2021). doi:10.1101/2020.12.27.20248738

29. Schaid, D. J., Chen, W. & Larson, N. B. From genome-wide associations to candidate causal variants by statistical fine-mapping. Nat. Rev. Genet. 19, 491–504 (2018).

30. Finucane, H. K. et al. Partitioning heritability by functional category using GWAS summary statistics. Nat. Genet. 47, 1228–1235 (2015).

31. Gazal, S. et al. Linkage disequilibrium–dependent architecture of human complex traits shows action of negative selection. Nat. Genet. 49, 1421–1427 (2017).

32. Watanabe, K., Taskesen, E., Bochoven, A. & Posthuma, D. Functional mapping and annotation of genetic associations with FUMA. Nat. Commun. 8, 1826 (2017).

33. de Leeuw, C. A. et al. MAGMA: Generalized Gene-Set Analysis of GWAS Data. PLoS Comput. Biol. 11, e1004219 (2015).

## References

34. Nagai, A. et al. Overview of the BioBank Japan Project: study design and profile. J. Epidemiol. 27, S2–S8 (2017).

35. Hirata, M. et al. Cross-sectional analysis of BioBank Japan clinical data: a large cohort of 200,000 patients with 47 common diseases. J. Epidemiol. 27, S9–S21 (2017).

36. Akiyama, M. et al. Characterizing rare and low-frequency height-associated variants in the Japanese population. Nat. Commun. 10, 1–11 (2019).

37. Kanai, M. et al. Genetic analysis of quantitative traits in the Japanese population links cell types to complex human diseases. Nat. Genet. 50, 390–400 (2018).

38. Team, H. Hail 0.2.

39. Chen, Z. et al. China Kadoorie Biobank of 0.5 million people: survey methods, baseline characteristics and long-term follow-up. Int. J. Epidemiol. 40, 1652–1666 (2011).

40. Bycroft, C. et al. The UK Biobank resource with deep phenotyping and genomic data. Nature 562, 203–209 (2018).

