## Supplementary Note and FAQ for "Multi-Ancestry Meta-Analysis yields novel genetic discoveries and ancestry-specific associations"

April 21, 2021

### 1 Theory

Here we propose a framework to combine GWAS summary statistics for corresponding to some trait in different ancestry groups. It is a generalization of the MTAG framework proposed in Turley et al.[1]

#### 1.1 Framework

The purpose of this method is to increase the precision of GWAS estimates for a particular phenotype in a particular ancestry group while reducing bias and controlling the type-1 error rate. This method uses sets of GWAS summary statistics for some phenotype (or for related phenotypes), possibly estimated in samples of different ancestries. It is closely related to the MTAG framework described in Turley et al.[1]. For simplicity throughout this document, we use "population" to refer to different sets of summary statistics that are used in the estimator even though the derivation holds when the method is applied to sets of summary statistics that are estimated from samples drawn from the same population or if the sets of summary statistics do not correspond to the same phenotype.

Assume we have a set of GWAS summary statistics for each of $P$ populations. For each SNP $j$, we denote the length-$P$ vector of GWAS estimates by $\hat{\beta}_j$. We denote the GWAS estimate for SNP $j$ in population $p$ by $\hat{\beta}_{p,j}$. This vector can be expressed as the sum of two components:

$$\hat{\beta}_j = \beta_j + \mathbf{e}_j,$$

where $\beta_j$ is the true genetic signal and $e_j$ is error, including sampling variation and biases. By "true genetic signal" we mean the GWAS estimate that would be produced if an infinitely-large sample size were available and we were able to perfectly control for population stratification and other biases. As in Turley et al. (2018), we treat each of these components as random variable and denote their variance-covariance matrices by $\mathbf{\Omega}_j \equiv \mathrm{Var}\left(\beta_j\right)$ and $\mathbf{\Sigma}_j \equiv \mathrm{Var}\left(\mathbf{e}_j\right)$.

If these matrices were known, an estimate of the marginal effect size of SNP $j$ on the trait in population $p$ could be estimated using generalized method of moments (GMM). As shown in Turley et al. (2018), projecting the GWAS estimate for SNP $j$ for each population onto the true marginal effect of SNP $j$ in population $p$, $\beta_{p,j}$, yields the following set of moment conditions

$$\mathbb{E}\left(\hat{\beta}_j - \frac{\mathbf{\Omega}'_{\cdot p,j}}{\Omega_{pp,j}}\beta_{p,j}\right) = 0,$$

where $\mathbf{\Omega}_{\cdot p,j}$ is the $p$-th column of $\mathbf{\Omega}_j$ and $\Omega_{pp,j}$ is the $p$-th entry of $\mathbf{\Omega}_{\cdot p,j}$ (or equivalently, the $p$-th diagonal entry of $\mathbf{\Omega}_j$). Using the efficient GMM weighting matrix, these moment conditions yield the Multi-Ancestry Meta-Analysis (MAMA) estimator

$$\hat{\beta}_{\mathrm{MAMA},j,p} = \frac{\frac{\mathbf{\Omega}'_{\cdot p,j}}{\Omega_{pp,j}}\left(\mathbf{\Omega}_j - \frac{\mathbf{\Omega}_{\cdot p,j}\mathbf{\Omega}'_{\cdot p,j}}{\Omega_{pp,j}} + \mathbf{\Sigma}_j\right)^{-1}\hat{\beta}_j}{\frac{\mathbf{\Omega}'_{\cdot p,j}}{\Omega_{pp,j}}\left(\mathbf{\Omega}_j - \frac{\mathbf{\Omega}_{\cdot p,j}\mathbf{\Omega}'_{\cdot p,j}}{\Omega_{pp,j}} + \mathbf{\Sigma}_j\right)^{-1}\frac{\mathbf{\Omega}_{\cdot p,j}}{\Omega_{pp,j}}}. \tag{1}$$

Also similar to Turley et al., the sampling variance of the MAMA estimator is

$$\text{Var}\left(\hat{\beta}_{\text{MAMA},j,p} - \beta_{j,p}|\beta_{j,p}\right) = \frac{1}{\frac{\boldsymbol{\Omega}'_{\cdot p,j}}{\Omega_{pp,j}}\left(\boldsymbol{\Omega}_j - \frac{\boldsymbol{\Omega}_{\cdot p,j}\boldsymbol{\Omega}'_{\cdot p,j}}{\Omega_{pp,j}} + \boldsymbol{\Sigma}_j\right)^{-1}\frac{\boldsymbol{\Omega}_{\cdot p,j}}{\Omega_{pp,j}}}. \tag{2}$$

The standard errors of a MAMA estimate can be obtained by taking the square root of (2) for each SNP $j$.

Recall that an important difference with MTAG is that the matrix $\boldsymbol{\Omega}_j$ can vary by SNP, unlike in Turley et al., where $\boldsymbol{\Omega}_j$ is assumed to be constant across SNPs. This variation across $j$ matters for both the MAMA estimates and their standard errors, and it is crucial for taking into account how the LD correlation across populations depends on the SNP.

The properties of the MAMA estimator are discussed in Section 1.5 of this document.

#### 1.2 Derivation of moment conditions for $\boldsymbol{\Omega}_j$ and $\boldsymbol{\Sigma}_j$

The MAMA estimator in equation (1) assumes that $\boldsymbol{\Omega}_j$ and $\boldsymbol{\Sigma}_j$ are known, but in practice they are not known and must be estimated. Here we describe how moment conditions for these matrices can be derived using a variation on the derivation of LD score regression[2].

In this derivation, we begin with an additive model for each population $p$:

$$y_{p,i} = \sum_k x_{p,k,i}b_{p,k} + F_{p,i} + \varepsilon_{p,i}$$

where $y_{p,i}$ is the phenotype value for individual $i$ in population $p$, $x_{p,k,i}$ is the genotype of that individual at SNP $k$, $b_{k,p}$ is the conditional effect of that SNP on $y_{p,i}$, $F_{p,i}$ is a set of unobserved confounding factors that may be correlated with $x_{p,k,i}$, and $\varepsilon_{p,i}$ is the residual, which is uncorrelated with $x_{p,k,i}$ by construction. The set of SNPs $\{x_{p,k,i}\}$ in the model are those from a reference panel and are assumed to be a dense set of SNPs (which may include SNPs that are monomorphic in population $p$). It is important to model the effects of a dense set of SNPs in order to capture the possibility that measured SNPs in the GWAS sample tag the (unmeasured) causal SNPs differently in different populations. Note that in this model, $b_{p,k}$ are effect sizes conditional on the set of SNPs in the reference panel as opposed to conditional on all SNPs (or all genetic variants) in the genome. Because we use a reference panel with a dense set of SNPs, we anticpate that this distinction will not be meaningful in practice.

In this derivation, our results are generalizable to any assumption about the units of the genotype. This allows for any transformation of the genotype (e.g., allele counts, standard deviations, dominance, etc.). However, whichever genotype-unit assumption is used will have implications for the relationship between allele-frequency and effect size within and across populations. We assume that $b_{p,k}$ are independent and identically distributed for all SNPs $k$ within a population $p$, and we denote the common variance of their effect sizes by $\omega_{pp} \equiv \text{Var}(b_{p,k})$. We denote the common covariance of SNPs' effect sizes between populations $p$ and $q$ by $\omega_{pq} \equiv \text{Cov}(b_{p,k}, b_{q,k})$. Note that the between-population covariance of the effect sizes is assumed to be constant across SNPs, but the between-population covariance may vary across pairs of populations. We refer to the assumption that $\omega_{pp}$ and $\omega_{pq}$ do not depend on the SNP $k$ for any populations $p$ and $q$ as the "homogeneous-$\omega$ assumption." Because of the homogeneous-$\omega$ assumption, the choice of the units of the genotype will lead to different implicit assumptions for the genetic architecture of the trait and for the relationship of the genetic achitectures across populations.

For example, if $x_{p,k,i}$ is measured in standard-deviation units, as is done in LD score regression[2], the homogeneous-$\omega$ assumption would imply that the effect of each allele for some SNP $k$ would tend to be larger in a population with a lower allele frequency, since each allele corresponds to a larger change in standard deviation units in that population. In cases where differences in allele frequency are driven by selection, such a model may be reasonable: if an allele is associated with decreased fitness in one population, then we would expect that allele to become more rare. If differences in allele frequency are instead driven by drift, then modeling SNP effects using allele-count units may be more reasonable.

GWAS summary statistics are estimates of the marginal effect of each SNP on the outcome. Therefore, the GWAS estimate of SNP $j$ on $y_{p,i}$ can be written as

$$\hat{\beta}_{p,j} = \sum_k \frac{r_{p,jk}}{r_{p,jj}} b_{p,k} + f_{p,j} + \xi_{p,j}, \tag{3}$$

where $r_{p,jk} \equiv \text{Cov}(x_{p,j,i}, x_{p,k,i})$ is the covariance between the genotypes for SNPs $j$ and $k$ in population $p$, $f_{p,j} \equiv \frac{\text{Cov}(x_{p,j,i},F_{p,i})}{r_{p,jj}}$ is the bias due to confounding, and $\xi_{p,j}$ is mean-zero estimation error. We also treat $f_{p,j}$ and $\xi_{p,j}$ as random, with $\text{Var}(f_{p,j}) \equiv \sigma_{f,pp}$, $\text{Cov}(f_{p,j}, f_{q,j}) \equiv \sigma_{f,pq}$, $\text{Var}(\xi_{p,j}) \equiv \sigma_{\xi,j,pp}$, and $\text{Cov}(\xi_{p,j}, \xi_{q,j}) \equiv \sigma_{\xi,pq,j}$. Note that we have implicitly assumed that $\sigma_{f,pp}$ and $\sigma_{f,pq}$ are constant across SNPs. This is consistent with a model of genetic drift. The variance and covariance of the estimation error, $\sigma_{\xi,pp,j}$ and $\sigma_{\xi,pq,j}$ respectively, will vary predictably from SNP to SNP as a function of allele frequency and sample size for that SNP. (If the genotypes are standardized and the sample size is constant across SNPs, then $\sigma_{\xi,pp,j} = \sigma_{\xi,pp,k}$.) From equation (3), we see that the first term is the part of the GWAS estimate due to true genetic factors, so $\beta_{p,j} = \sum_k \frac{r_{p,jk}}{r_{p,jj}} b_{p,k}$. The remaining terms are the error: $e_{p,j} = f_{p,j} + \xi_{p,j}$.

We obtain the matrices $\mathbf{\Omega}_j$ and $\mathbf{\Sigma}_j$ following a similar strategy to that of LD score regression. To obtain the $p$-th diagonal elements of these matrices, we assume that (i) $\text{Cov}(b_{p,j}, b_{p,k}) = 0$ for all $j \neq k$, (ii) $\text{Cov}(b_{p,k}, f_{p,j}) = 0$ for all $j, k$, (iii) $\text{Cov}(b_{p,k}, \xi_{p,j}) = 0$ for all $j, k$, and (iv) $\text{Cov}(f_{p,j}, \xi_{p,j}) = 0$ for all $j$. It then follows that

$$\begin{aligned}
\text{Var}\left(\hat{\beta}_{p,j}\right) &= \text{Var}\left(\sum_k \frac{r_{p,jk}}{r_{p,jj}} b_{p,k} + f_{p,j} + \xi_{p,j}\right) \\
&= \sum_k \frac{r_{p,jk}^2}{r_{p,jj}^2} \text{Var}(b_{p,k}) + \text{Var}(f_{p,j}) + \text{Var}(\xi_{p,j}) \\
&= \omega_{pp} \sum_k \frac{r_{p,jk}^2}{r_{p,jj}^2} + \sigma_{f,pp} + \sigma_{\xi,pp,j} \\
&= \omega_{pp} \ell_{pp,j} + \sigma_{f,pp} + \sigma_{\xi,pp,j} \\
&= \Omega_{pp,j} + \Sigma_{pp,j},
\end{aligned}$$

where $\Omega_{pp,j} \equiv \omega_{pp} \ell_{pp,j}$ is the $p$-th diagonal element of $\mathbf{\Omega}_j$, $\mathbf{\Sigma}_{pp,j} \equiv \sigma_{f,pp} + \sigma_{\xi,pp,j}$ is the $p$-th diagonal element of $\mathbf{\Sigma}_j$, and $\ell_{pp,j}$ is a generalization of the LD score for SNP $j$ in population $p$:

$$\ell_{pp,j} \equiv \sum_k \frac{r_{p,jk}^2}{r_{p,jj}^2}. \tag{4}$$

For the off-diagonal terms corresponding to populations $p$ and $q$, we have

$$\begin{aligned}
\text{Cov}\left(\hat{\beta}_{p,j}, \hat{\beta}_{q,j}\right) &= \text{Cov}\left(\sum_k \frac{r_{p,jk}}{r_{p,jj}} b_{p,k} + f_{p,j} + \xi_{p,j}, \sum_k \frac{r_{q,jk}}{r_{q,jj}} b_{q,k} + f_{q,j} + \xi_{q,j}\right) \\
&= \sum_k \frac{r_{p,jk}}{r_{p,jj}} \frac{r_{q,jk}}{r_{q,jj}} \text{Cov}(b_{p,k}, b_{q,k}) + \text{Cov}(f_{p,j}, f_{q,j}) + \text{Cov}(\xi_{p,j}, \xi_{q,j}) \\
&= \omega_{pq} \sum_k \frac{r_{p,jk}}{r_{p,jj}} \frac{r_{q,jk}}{r_{q,jj}} + \sigma_{f,pq} + \sigma_{\xi,pq,j} \\
&= \omega_{pq} \ell_{pq,j} + \sigma_{f,pq} + \sigma_{\xi,pq,j} \\
&= \Omega_{pq,j} + \Sigma_{pq,j},
\end{aligned}$$

where $\Omega_{pq,j} \equiv \omega_{pq} \ell_{pq,j}$ is the $(p, q)$-th element of $\mathbf{\Omega}_j$, $\Sigma_{pq,j} \equiv \sigma_{f,pq} + \sigma_{\xi,pq,j}$ is the $(p, q)$-th element of $\mathbf{\Sigma}_j$, and $\ell_{pq,j}$ is a cross-ancestry generalization of an LD score for SNP $j$:

$$\ell_{pq,j} \equiv \sum_k \frac{r_{p,jk} r_{q,jk}}{r_{p,jj} r_{q,jj}}. \tag{5}$$

This cross-ancestry LD score is a generalization of the LD score used in the cross-ancestry genetic correlation method POPCORN[3].

#### 1.3 Estimation of $\mathbf{\Omega}_j$ and $\mathbf{\Sigma}_j$

For MAMA, we estimate $\mathbf{\Omega}_j$ and $\mathbf{\Sigma}_j$ using a variation on LD score regression. Recall that

$$\text{Var}\left(\hat{\beta}_{p,j}\right) = \omega_{pp} \ell_{pp,j} + \sigma_{f,pp} + \sigma_{\xi,pp,j}.$$

We will consider each of these terms in turn. First, under the homogeneous-$\omega$ assumption, $\omega_{pp}$ does not vary by SNP. Therefore, the first term is proportional to the LD score for population $p$, $\ell_{pp,j}$. The value of the second term, $\sigma_{f,pp}$, which corresponds to the variation in the GWAS summary statistics due to bias, is constant across SNPs under a model of drift. Finally, the third term, $\sigma_{\xi,pp,j}$, corresponds to the estimation error of the GWAS estimate. It therefore should be equal to the squared standard error, which we denote by $S_{p,j} \equiv \left[\text{SE}\left(\hat{\beta}_{p,j}\right)\right]^2$. However, if there is cryptic relatedness or sample overlap within the GWAS summary statistics for some population, the standard errors will be too small (i.e., $S_{p,j} < \sigma_{\xi,pp,j}$). Even then, assuming this source of bias is constant across SNPs, $S_{p,j}$ will be proportional to $\sigma_{\xi,pp,j}$.

Given these relationships, for each population $p$, we can regress the squared GWAS estimates on the population LD score, the squared GWAS standard error and constant:

$$\hat{\beta}_{p,j}^2 = \gamma_{\ell,pp}\ell_{pp,j} + \gamma_{f,pp} + \gamma_{\xi,p}S_{p,j}. \tag{6}$$

We can then use the estimated coeffients from this regression to obtain estimates of the diagonals for $\mathbf{\Omega}_j$ and $\mathbf{\Sigma}_j$. Specifically, we will use

$$\begin{aligned}\hat{\Omega}_{pp,j} &= \hat{\gamma}_{\ell,pp}\ell_{pp,j} \\ \hat{\Sigma}_{pp,j} &= \hat{\gamma}_{f,pp} + \hat{\gamma}_{\xi,p}S_{p,j}.\end{aligned}$$

When there is little variation in the sample size across SNPs within each population, $S_{p,j}$ will approximately constant across SNPs, making it nearly collinear with the constant term. For this reason, except in cases where there is substantial variation in $S_{p,j}$, MAMA will be more stable by simply regressing $\hat{\beta}_{p,j}^2$ on the population LD score and a constant. In this simplified specification, the constant will capture both $\sigma_{f,pp}$ and $\sigma_{\xi,pp,j}$, and $\hat{\Sigma}_{pp,j}$ is set equal to the constant.

The off-diagonal elements of $\mathbf{\Omega}_j$ and $\mathbf{\Sigma}_j$ are constructed in a similar way. Recall that

$$\text{Cov}\left(\hat{\beta}_{p,j}, \hat{\beta}_{q,j}\right) = \omega_{pq}\ell_{pq,j} + \sigma_{f,pq} + \sigma_{\xi,pq,j}.$$

As before, $\omega_{pq}$ is constant across SNPs, so the first term is proportional to the cross-population LD score, $\ell_{pq,j}$. Also, the second term, $\sigma_{pq,f}$, does not vary across SNPs under a model of genetic drift. The third term corresponds to the sampling covariance in the two sets of summary statistics. Since the samples for populations $p$ and $q$ necessarily don't overlap (since they are made up of individuals from different ancestral populations), $\sigma_{\xi,pq,j} = 0$.

Therefore, in the cross-population case, we regress the product of GWAS summary statistics from the pair of populations on the cross-population LD score and a constant:

$$\hat{\beta}_{p,j}\hat{\beta}_{q,j} = \gamma_{\ell,pq}\ell_{pq,j} + \gamma_{f,pq}. \tag{7}$$

We then set the off-diagonal elements of the estimated $\mathbf{\Omega}_j$ and $\mathbf{\Sigma}_j$ matrices as:

$$\begin{aligned}\hat{\Omega}_{pq,j} &= \hat{\gamma}_{\ell,pq}\ell_{pq,j} \\ \hat{\Sigma}_{j,pq} &= \hat{\gamma}_{f,pq}.\end{aligned}$$

Having constructed all elements of $\hat{\mathbf{\Omega}}$ and $\hat{\mathbf{\Sigma}}$, these parameters and the GWAS estimates can be substituted into equation (1) to produce MAMA estimates.

#### 1.4 Restrictions on $\mathbf{\Omega}_j$ and $\mathbf{\Sigma}_j$

In some circumstances, we may wish to impose restriction on the coefficients of the LD score regression described above (and therefore restrictions on $\mathbf{\Omega}_j$ and $\mathbf{\Sigma}_j$). Below we describe a few of these potential restrictions

##### 1.4.1 Uncorrelated stratification bias across populations

In the model described above, $\sigma_{f,pq} = \text{Cov}(f_{p,j}, f_{q,j})$ is the covariance of the bias term in GWAS estimates between populations. If the two populations are closely related (e.g., two populations within Europe), we may expect that this covariance is non-zero. However, in most applications of MAMA between continental populations, we do not expect the bias for any particular phenotype to be correlated across populations. For this reason, in the applications in this paper and in the software defaults, we assume $\sigma_{f,pq} = 0$. In practice, this is done by assuming that $\gamma_{f,pq} = 0$ when we estimate equation (7). Then $\boldsymbol{\Omega}_j$ and $\boldsymbol{\Sigma}_j$ are constructed for each SNP using the coefficient estimates from this restricted regression.

##### 1.4.2 Perfect genetic correlation across populations (or some other assumed, known genetic correlation)

If the heritability of a phenotype and the GWAS sample size is small for one or more populations, then the estimate of $\boldsymbol{\Omega}_j$ may be very imprecise, which can cause MAMA estimates to be imprecise. In addition, the point estimates of the elements of $\boldsymbol{\Omega}_j$ are likely to be inaccurate, which can cause MAMA estimates to be biased. For these reasons, it may be preferable to assume that the conditional genetic effects are perfectly correlated across populations.

Using the notation above, conditional effect sizes between populations $p$ and $q$ will be perfectly correlated if $\omega_{pq}/\sqrt{\omega_{pp}\omega_{qq}} = 1$. In practice, we impose this restriction by first freely estimating $\hat{\gamma}_{\ell,pp}$ and $\hat{\gamma}_{\ell,qq}$ from equation (6) for each population. We then estimate equation (7), with the restriction that $\hat{\gamma}_{\ell,pq} = \sqrt{\hat{\gamma}_{\ell,pp}\hat{\gamma}_{\ell,qq}}$. Then $\boldsymbol{\Omega}_j$ and $\boldsymbol{\Sigma}_j$ are constructed for each SNP using the coefficient estimates from this restricted regression. Notice that by estimating $\hat{\gamma}_{\ell,pp}$ and $\hat{\gamma}_{\ell,qq}$ freely, we allow the heritability of the phenotype to differ across populations.

While it is plausible that the perfect genetic correlation assumption will hold for many phenotypes (see Supplementary Table 1), it is still important to verify that the SNPs identified by MAMA under this assumption are robust. We recommend that researchers replicate their findings in an independent GWAS sample if available. We also recommend that researchers assess what fraction of SNPs remain genome-wide significant if a lower genetic correlation is assumed. This is done similarly to above, instead restricting equation (7) such that $\hat{\gamma}_{\ell,pq} = r_{g,pq}\sqrt{\hat{\gamma}_{\ell,pp}\hat{\gamma}_{\ell,qq}}$, where $r_{g,pq}$ is the assumed genetic correlation between the populations.

In our applications in this paper, we make the perfect-genetic-correlation assumption. We therefore replicate the lead SNPs identified for all phenotypes except for schizophrenia (since no replication sample was available), and we verify the robustness of our results assuming $r_{g,pq} = 0.9$.

#### 1.5 Properties of MAMA

Because the MAMA estimator has the same functional form as MTAG at the SNP level, MAMA will have all of the same statistical properties as MTAG. For example, MAMA is a consistent estimator of $\beta_{p,j}$ as either the number of populations increases or as the sample size of the target population increases. Also, the expected mean squared error of of the MAMA estimate for some population will be strictly smaller than the expected mean squared error of the corresponding GWAS estimate. Proofs of these results are found in Turley et al.[1] Due to the treatment of differences in LD structure, MAMA also has additional properties that are described in this section.

##### 1.5.1 Mean Squared Error and LD Score Correlation

The gains in power due to MAMA are not uniformly spread across the genome. MAMA will increase statistical power most in regions where LD patterns are most similar across populations. We illustrate this property below with a simple two-population example.

We will show that the mean squared error of a MAMA coefficient is decreasing the LD correlation of the SNP. We denote the *LD score correlation* between populations $p$ and $q$ at SNP $j$ by

$$r_{LD,pq,j} \equiv \frac{\ell_{pq,j}}{\sqrt{\ell_{pp,j}\ell_{qq,j}}}. \tag{8}$$

While not formally a correlation, $r_{LD,pq,j}$ has the intuitive properties of a correlation. For example, from equations (4) and (5), if $r_{p,jk} = r_{q,jk}$ for every $k$, then $\ell_{pq,j} = \ell_{pp,j} = \ell_{qq,j}$ and therefore $r_{LD,pq,j} = 1$. Similarly, if $r_{p,jk}$ is uncorrelated with $r_{q,jk}$, then $r_{LD,pq,j} = \ell_{pq,j} = 0$. So $r_{LD,pq,j}$ may be considered a metric of the similarity of LD patterns local to SNP $j$.

We will assume that $r_{LD,pq,j} \geq 0$ through this proof. This is a weak assumption because we expect genetically related populations to have similar LD structure. Over time, due to genetic drift, $r_{LD,pq,j}$ is expected to fall in magnitude until $\mathbb{E}\left(r_{p,jk}r_{q,jk}\right) = 0$ for $j \neq k$. However, $\mathbb{E}\left(r_{p,jj}r_{q,jj}\right) \geq 0$ since $r_{p,jj} \geq 0$ and $r_{q,jj} \geq 0$. This mean that $\ell_{pq,j} \equiv \sum_k \left(r_{p,jk}r_{q,jk}\right) / \left(r_{p,jj}r_{q,jj}\right)$ will converge over time to some non-negative number.

Using the notation from equation (8) in a two-population setting, we can express $\mathbf{\Omega}_j$ as

$$\mathbf{\Omega}_j = \begin{bmatrix} \ell_{pp,j}\omega_{pp} & r_{LD,pq,j}\sqrt{\ell_{pp,j}\ell_{qq,j}}\omega_{pq} \\ r_{LD,pq,j}\sqrt{\ell_{pp,j}\ell_{qq,j}}\omega_{pq} & \ell_{qq,j}\omega_{qq} \end{bmatrix}.$$

Also, since the data for the two populations are non-overlapping, $\mathbf{\Sigma}_j$ is the diagonal matrix

$$\mathbf{\Sigma}_j = \begin{bmatrix} \Sigma_{pp,j} & 0 \\ 0 & \Sigma_{qq,j} \end{bmatrix}.$$

The mean squared error of the estimate for SNP $j$ in population $p$ is

$$\begin{aligned}
\mathrm{MSE}_{p,j} &= \frac{1}{\frac{\mathbf{\Omega}'_{\cdot p,j}}{\Omega_{pp,j}}\left(\mathbf{\Omega}_j - \frac{\mathbf{\Omega}_{\cdot p,j}\mathbf{\Omega}'_{\cdot p,j}}{\Omega_{pp,j}} + \mathbf{\Sigma}_j\right)^{-1}\frac{\mathbf{\Omega}_{\cdot p,j}}{\Omega_{pp,j}}} \\
&= \frac{1}{\begin{bmatrix} 1 \\ r_{LD,pq,j}\sqrt{\frac{\ell_{qq,j}}{\ell_{pp,j}}}\frac{\omega_{pq}}{\omega_{pp}} \end{bmatrix}' \begin{bmatrix} \Sigma_{pp,j} & 0 \\ 0 & \ell_{qq,j}\omega_{qq} - r^2_{LD,pq,j}\ell_{qq,j}\frac{\omega^2_{pq}}{\omega_{pp}} + \Sigma_{qq,j} \end{bmatrix}^{-1} \begin{bmatrix} 1 \\ r_{LD,pq,j}\sqrt{\frac{\ell_{qq,j}}{\ell_{pp,j}}}\frac{\omega_{pq}}{\omega_{pp}} \end{bmatrix}} \\
&= \frac{1}{\begin{bmatrix} 1 \\ r_{LD,pq,j}\sqrt{\frac{\ell_{qq,j}}{\ell_{pp,j}}}\frac{\omega_{pq}}{\omega_{pp}} \end{bmatrix}' \begin{bmatrix} \frac{1}{\Sigma_{pp,j}} & 0 \\ 0 & \frac{1}{\ell_{qq,j}\omega_{qq} - r^2_{LD,pq,j}\ell_{qq,j}\frac{\omega^2_{pq}}{\omega_{pp}} + \Sigma_{qq,j}} \end{bmatrix} \begin{bmatrix} 1 \\ r_{LD,pq,j}\sqrt{\frac{\ell_{qq,j}}{\ell_{pp,j}}}\frac{\omega_{pq}}{\omega_{pp}} \end{bmatrix}} \\
&= \frac{1}{\frac{1}{\Sigma_{pp,j}} + r^2_{LD,pq,j}\frac{\ell_{qq,j}}{\ell_{pp,j}}\frac{\omega^2_{pq}}{\omega^2_{pp}}\frac{1}{\ell_{qq,j}\omega_{qq} - r^2_{LD,pq,j}\ell_{qq,j}\frac{\omega^2_{pq}}{\omega_{pp}} + \Sigma_{qq,j}}} \\
&= \frac{1}{\frac{1}{\Sigma_{pp,j}} + \frac{r^2_{LD,pq,j}\ell_{qq,j}\omega^2_{pq}}{\ell_{pp,j}\ell_{qq,j}\omega^2_{pp}\omega_{qq} - r^2_{LD,pq,j}\ell_{pp,j}\ell_{qq,j}\omega_{pp}\omega^2_{pq} + \ell_{pp,j}\omega^2_{pp}\Sigma_{qq,j}}}.
\end{aligned}$$

Note that if $r_{LD,pq,j} \geq 0$, then

$$\frac{\partial}{\partial r_{LD,pq,j}} r^2_{LD,pq,j}\ell_{qq,j}\omega^2_{pq} = 2r_{LD,pq,j}\ell_{qq,j}\omega^2_{pq} \geq 0,$$

and

$$\frac{\partial}{\partial r_{LD,pq,j}}\left(\ell_{pp,j}\ell_{qq,j}\omega^2_{pp}\omega_{qq} - r^2_{LD,pq,j}\ell_{pp,j}\ell_{qq,j}\omega_{pp}\omega^2_{pq} + \ell_{pp,j}\omega^2_{pp}\Sigma_{qq,j}\right) = -2r_{LD,pq,j}\ell_{pp,j}\ell_{qq,j}\omega_{pp}\omega^2_{pq} \leq 0.$$

Therefore,

$$\frac{\partial}{\partial r_{LD,pq,j}}\left(\frac{1}{\Sigma_{pp,j}} + \frac{r^2_{LD,pq,j}\ell_{qq,j}\omega^2_{pq}}{\ell_{pp,j}\ell_{qq,j}\omega^2_{pp}\omega_{qq} - r^2_{LD,pq,j}\ell_{pp,j}\ell_{qq,j}\omega_{pp}\omega^2_{pq} + \ell_{pp,j}\omega^2_{pp}\Sigma_{qq,j}}\right) \geq 0,$$

which in turn implies that

$$\frac{\partial}{\partial r_{LD,pq,j}}\mathrm{MSE}_j = \frac{\partial}{\partial \rho_{\ell,j}}\frac{1}{\frac{1}{\Sigma_{pp,j}} + \frac{r^2_{LD,pq,j}\ell_{qq,j}\omega^2_{pq}}{\ell_{pp,j}\ell_{qq,j}\omega^2_{pp}\omega_{qq} - r^2_{LD,pq,j}\ell_{pp,j}\ell_{qq,j}\omega_{pp}\omega^2_{pq} + \ell_{pp,j}\omega^2_{pp}\Sigma_{qq,j}}} \leq 0.$$

Furthermore, these inequalities will be strict when $r_{LD,pq,j} > 0$. Thus, as long as $r_{LD,pq,j} \geq 0$, the mean squared error of MAMA estimates decreases most in regions where LD patterns are most similar.

##### 1.5.2 MAMA is the Best Linear Unbiased Estimator

Here we will show that MAMA is the best linear unbiased estimator. The proof will proceed as follows. First, we will show that MAMA is an unbiased estimator. Second, we will establish a necessary condition for the set of all linear, unbiased estimators. Finally, we will show that MAMA has the lowest variance of all such estimators.

*Unbiasedness of MAMA.* As shown in the derivation of the MAMA estimator, we know that

$$\mathbb{E}\left(\hat{\beta}_j\right) = \frac{\mathbf{\Omega}_{\cdot p,j}}{\Omega_{pp,j}}\beta_{p,j}.$$

Therefore, the the expected value of the MAMA estimator is

$$\begin{aligned}
\mathbb{E}\left(\hat{\beta}_{\text{MAMA},p,j}\right) &= \mathbb{E}\left(\frac{\frac{\mathbf{\Omega}'_{\cdot p,j}}{\Omega_{pp,j}}\left(\mathbf{\Omega}_j - \frac{\mathbf{\Omega}_{\cdot p,j}\mathbf{\Omega}'_{\cdot p,j}}{\Omega_{pp,j}} + \mathbf{\Sigma}_j\right)^{-1}\hat{\beta}_j}{\frac{\mathbf{\Omega}'_{\cdot p,j}}{\Omega_{pp,j}}\left(\mathbf{\Omega}_j - \frac{\mathbf{\Omega}_{\cdot p,j}\mathbf{\Omega}'_{\cdot p,j}}{\Omega_{pp,j}} + \mathbf{\Sigma}_j\right)^{-1}\frac{\mathbf{\Omega}_{\cdot p,j}}{\Omega_{pp,j}}}\right) \\
&= \frac{\frac{\mathbf{\Omega}'_{\cdot p,j}}{\Omega_{pp,j}}\left(\mathbf{\Omega}_j - \frac{\mathbf{\Omega}_{\cdot p,j}\mathbf{\Omega}'_{\cdot p,j}}{\Omega_{pp,j}} + \mathbf{\Sigma}_j\right)^{-1}\mathbb{E}\left(\hat{\beta}_j\right)}{\frac{\mathbf{\Omega}'_{\cdot p,j}}{\Omega_{pp,j}}\left(\mathbf{\Omega}_j - \frac{\mathbf{\Omega}_{\cdot p,j}\mathbf{\Omega}'_{\cdot p,j}}{\Omega_{pp,j}} + \mathbf{\Sigma}_j\right)^{-1}\frac{\mathbf{\Omega}_{\cdot p,j}}{\Omega_{pp,j}}} \\
&= \frac{\frac{\mathbf{\Omega}'_{\cdot p,j}}{\Omega_{pp,j}}\left(\mathbf{\Omega}_j - \frac{\mathbf{\Omega}_{\cdot p,j}\mathbf{\Omega}'_{\cdot p,j}}{\Omega_{pp,j}} + \mathbf{\Sigma}_j\right)^{-1}\frac{\mathbf{\Omega}_{\cdot p,j}}{\Omega_{pp,j}}\beta_{p,j}}{\frac{\mathbf{\Omega}'_{\cdot p,j}}{\Omega_{pp,j}}\left(\mathbf{\Omega}_j - \frac{\mathbf{\Omega}_{\cdot p,j}\mathbf{\Omega}'_{\cdot p,j}}{\Omega_{pp,j}} + \mathbf{\Sigma}_j\right)^{-1}\frac{\mathbf{\Omega}_{\cdot p,j}}{\Omega_{pp,j}}} \\
&= \beta_{p,j}.
\end{aligned}$$

So MAMA is an unbiased estimator of $\beta_{p,j}$.

*Necessary condition for linear, unbiased estimators.* For ease of notation, we define the MAMA weight vector as

$$\mathbf{W} \equiv \frac{\frac{\mathbf{\Omega}'_{\cdot p,j}}{\Omega_{pp,j}}\left(\mathbf{\Omega}_j - \frac{\mathbf{\Omega}_{\cdot p,j}\mathbf{\Omega}'_{\cdot p,j}}{\Omega_{pp,j}} + \mathbf{\Sigma}_j\right)^{-1}}{\frac{\mathbf{\Omega}'_{\cdot p,j}}{\Omega_{pp,j}}\left(\mathbf{\Omega}_j - \frac{\mathbf{\Omega}_{\cdot p,j}\mathbf{\Omega}'_{\cdot p,j}}{\Omega_{pp,j}} + \mathbf{\Sigma}_j\right)^{-1}\frac{\mathbf{\Omega}_{\cdot p,j}}{\Omega_{pp,j}}}$$

such that $\beta_{MAMA,p,j} = \mathbf{W}\hat{\beta}_j$. Notice that the set of all linear estimators can be characterized as $\mathbf{W} + \mathbf{C}$ for some vector $\mathbf{C}$. By the definition of unbiasedness,

$$\mathbb{E}\left[\left(\mathbf{W} + \mathbf{C}\right)\hat{\beta}_j\right] = \beta_{p,j}.$$

It follows that

$$\begin{aligned}
0 &= \mathbb{E}\left(\mathbf{W}\hat{\beta}_j + \mathbf{C}\hat{\beta}_j\right) - \beta_{p,j} \\
&= \mathbb{E}\left(\mathbf{W}\hat{\beta}_j\right) + \mathbf{C}\mathbb{E}\left(\hat{\beta}_j\right) - \beta_{p,j} \\
&= \mathbb{E}\left(\hat{\beta}_{MAMA,p,j}\right) + \mathbf{C}\mathbb{E}\left(\hat{\beta}_j\right) - \beta_{p,j} \\
&= \beta_{p,j} + \mathbf{C}\mathbb{E}\left(\hat{\beta}_j\right) - \beta_{p,j} \text{ (by the unbiasednes of MAMA)} \\
&= \beta_{p,j} + \mathbf{C}\frac{\mathbf{\Omega}_{\cdot p,j}}{\Omega_{pp,j}}\beta_{p,j} - \beta_{p,j} \text{ (by MAMA's moment conditions)} \\
&= \mathbf{C}\frac{\mathbf{\Omega}_{\cdot p,j}}{\Omega_{pp,j}}\beta_{p,j} \\
0 &= \mathbf{C}\mathbf{\Omega}_{\cdot p,j}. \text{ (multiplying the right and left by } \Omega_{pp,j}/\beta_{p,j}\text{)}
\end{aligned}$$

So $\left(\mathbf{W} + \mathbf{C}\right)\hat{\beta}_j$ will only be an unbiased estimator if $\mathbf{C}\mathbf{\Omega}_{\cdot p,j} = 0$.

*MAMA has lowest variance of all linear, unbiased estimators.* Assume that $(\mathbf{W}+\mathbf{C})\hat{\beta}_j$ is an unbiased estimator. We will show that the variance of $(\mathbf{W}+\mathbf{C})\hat{\beta}_j$ is greater than the variance of $\mathbf{W}\hat{\beta}_j$. We calculate

$$\begin{aligned}\text{Var}\left[(\mathbf{W}+\mathbf{C})\hat{\beta}_j\right]-\text{Var}\left(\mathbf{W}\hat{\beta}_j\right) &= (\mathbf{W}+\mathbf{C})\,\text{Var}\left(\hat{\beta}_j\right)(\mathbf{W}+\mathbf{C})'-\mathbf{W}\text{Var}\left(\hat{\beta}_j\right)\mathbf{W}' \\ &= \mathbf{W}\text{Var}\left(\hat{\beta}_j\right)\mathbf{W}'+2\mathbf{C}\text{Var}\left(\hat{\beta}_j\right)\mathbf{W}' \\ &\quad+\mathbf{C}\text{Var}\left(\hat{\beta}_j\right)\mathbf{C}'-\mathbf{W}\text{Var}\left(\hat{\beta}_j\right)\mathbf{W}' \\ &= 2\mathbf{C}\text{Var}\left(\hat{\beta}_j\right)\mathbf{W}'+\mathbf{C}\text{Var}\left(\hat{\beta}_j\right)\mathbf{C}'. \qquad (9)\end{aligned}$$

Consider the first term of this expression:

$$\begin{aligned}2\mathbf{C}\text{Var}\left(\hat{\beta}_j\right)\mathbf{W}' &= 2\mathbf{C}\text{Var}\left(\hat{\beta}_j\right)\frac{\left(\mathbf{\Omega}_j-\frac{\mathbf{\Omega}_{\cdot p,j}\mathbf{\Omega}'_{\cdot p,j}}{\Omega_{pp,j}}+\mathbf{\Sigma}_j\right)^{-1}\frac{\mathbf{\Omega}_{\cdot p,j}}{\Omega_{pp,j}}}{\frac{\mathbf{\Omega}'_{\cdot p,j}}{\Omega_{pp,j}}\left(\mathbf{\Omega}_j-\frac{\mathbf{\Omega}_{\cdot p,j}\mathbf{\Omega}'_{\cdot p,j}}{\Omega_{pp,j}}+\mathbf{\Sigma}_j\right)^{-1}\frac{\mathbf{\Omega}_{\cdot p,j}}{\Omega_{pp,j}}} \\ &= 2\mathbf{C}\left(\mathbf{\Omega}_j-\frac{\mathbf{\Omega}_{\cdot p,j}\mathbf{\Omega}'_{\cdot p,j}}{\Omega_{pp,j}}+\mathbf{\Sigma}_j\right)\frac{\left(\mathbf{\Omega}_j-\frac{\mathbf{\Omega}_{\cdot p,j}\mathbf{\Omega}'_{\cdot p,j}}{\Omega_{pp,j}}+\mathbf{\Sigma}_j\right)^{-1}\frac{\mathbf{\Omega}_{\cdot p,j}}{\Omega_{pp,j}}}{\frac{\mathbf{\Omega}'_{\cdot p,j}}{\Omega_{pp,j}}\left(\mathbf{\Omega}_j-\frac{\mathbf{\Omega}_{\cdot p,j}\mathbf{\Omega}'_{\cdot p,j}}{\Omega_{pp,j}}+\mathbf{\Sigma}_j\right)^{-1}\frac{\mathbf{\Omega}_{\cdot p,j}}{\Omega_{pp,j}}} \\ &= 2\frac{\mathbf{C}\frac{\mathbf{\Omega}_{\cdot p,j}}{\Omega_{pp,j}}}{\frac{\mathbf{\Omega}'_{\cdot p,j}}{\Omega_{pp,j}}\left(\mathbf{\Omega}_j-\frac{\mathbf{\Omega}_{\cdot p,j}\mathbf{\Omega}'_{\cdot p,j}}{\Omega_{pp,j}}+\mathbf{\Sigma}_j\right)^{-1}\frac{\mathbf{\Omega}_{\cdot p,j}}{\Omega_{pp,j}}} \\ &= 0.\end{aligned}$$

The last step follows from the necessary condition for unbiasedness derived above. Substituting this into (9), we have

$$\text{Var}\left[(\mathbf{W}+\mathbf{C})\hat{\beta}_j\right]-\text{Var}\left(\mathbf{W}\hat{\beta}_j\right)=\mathbf{C}\text{Var}\left(\hat{\beta}_j\right)\mathbf{C}'\geq 0.$$

This last step follows because variance-covariance matrices are positive semidefinite. Therefore

$$\text{Var}\left[(\mathbf{W}+\mathbf{C})\hat{\beta}_j\right]\geq\text{Var}\left(\mathbf{W}\hat{\beta}_j\right).$$

So MAMA has the (weakly) minimum variance of all linear, unbiased estimators.

##### 1.5.3 The Homogeneous-Omega Assumption

A key assumption of MTAG is

$$\text{Var}\left(\beta_j\right)=\mathbf{\Omega},$$

where $\Omega$ is not a function of $j$. This effectively assumes that SNPs throughout the genome are exchangeable and that the relationship between SNP associations across traits does not vary as a function of SNP characteristics. A primary example of when this assumption would be violated is if there is a mass of SNPs that are associated with one trait but not another and a different mass of SNPs that are associated with both traits. In such a case, there would be zero covariance between SNP associations for the former and some non-zero covariance for the latter.

In MAMA, the assumption is slightly different. Namely, we assume that

$$\text{Var}\left(\mathbf{b}_j\right)=\omega,$$

where the $(p,q)^{\text{th}}$ element of $\omega$ is $\omega_{pq}$. Similar to MTAG, we assume that $\omega$ is not a function of $j$ or the attributes of SNP $j$. In contrast to MTAG's assumption, however, which is based on the marginal associations, MAMA's key assumptions is based on the associations *conditional on all other SNPs.* In cases when MAMA is used to meta-analyze summary statisitics corresponding to the same phenotype, we anticipate that the homogeneous-omega assumption is likely to be an excellent approximation for most phenotypes.

Substantial violations of the assumption would be expected to occur only if there are different biological pathways across different populations.

We caution, however, that if MAMA is used to meta-analyze different phenotypes across different populations, the same risks of an increased false-discovery rate and difficulties in interpretation that are discussed in Turley et al. (2018) apply to MAMA coefficient estimates.

##### 1.5.4 Polygenic Scores

As with MTAG, MAMA coefficient estimates will have smaller mean squared error than their corresponding GWAS estimates. As a result, we can generally anticipate that polygenic scores based on MAMA results will be more predictive than polygenic scores based on GWAS results. It is not the case, however, that MAMA-based summary statistics will have greater predictive power than polygenic scores based on other meta-analysis methods, including polygenic scores based on simple, variance-weighted, meta-analysis. The reason for this is that MAMA estimates are unbiased. In producing polygenic scores, there is a trade-off between reducing the bias and reducing the variance of coefficient estimates. In virtually all cases, predictive power can be improved by allowing for some bias in exchange for reduced variance.

##### 1.5.5 Computational Performance of the MAMA software

MAMA is computationally efficient because of all its steps have closed-form solutions. To benchmark its performance, we timed five identical runs of the BMI EAS-EUR meta-analysis (9.3M SNPs in raw EAS discovery GWAS, 13.4M SNPs in raw EUR discovery GWAS) on a single core of a 2.60 GHz Intel(R) Xeon(R) CPU E5-4627 v3 processor. The median runtime was 18 minutes and 38 seconds, inclusive of file I/O, GWAS QC, and the MAMA algorithm. Of course, run time may vary as a function of the computing environment.

### 2 Quality Control

Quality control (QC) of the genotypic data used in each of the GWAS differed by cohort. The QC criteria for SNPs included thresholds on minor allele frequency, call rate, Hardy-Weinberg Equilibrium $P$ value and imputation quality. Each cohort also had subject-level QC criteria, including a call-rate threshold. The thresholds used by each cohort can be found in Supplementary Table 3.

The complete set of GWAS summary statistics for 28 different phenotypes received for this project, the estimated heritability from LD Score regression[2], discovery and replication sample sizes, and phenotypic mean and standard deviation are found in Supplementary Table 2. Five of the phenotypes from the China Kadoorie Biobank (Life satisfaction, Mother's age of death, Number of children ever born, Self-rated health, and Smoking cessation) had heritability estimates that were not statistically distinguishable from zero at a $P$ value threshold of 0.05, so these phenotypes were omitted from further analysis.

Upon receiving the GWAS summary statistics, some SNPs were filtered out using an additional QC protocol. The QC criteria used are based on the criteria set by the "munge" step of the original LD Score Regression software. An additional filter that is used in our QC protocol is to only include SNPs that are in a master SNPlist panel. These are SNPs that are included in the HapMap3 set of SNPs and which have a minor allele frequency of at least 1% in both the EAS or EUR sample of the 1000 Genome Project data[4]. By these criteria, the SNPlist includes 980,861 SNPs. This is done to ensure that each SNP passed into the MAMA software is reliably measured and is available in both sets of summary statistics. These filters and the number of SNP dropped in each step of QC is found in Supplementary Table 4.

### 3 Acknowledgements

#### 3.1 China Kadoorie Biobank

**International Steering Committee:** Junshi Chen, Zhengming Chen (PI), Robert Clarke, Rory Collins, Yu Guo, Liming Li (PI), Chen Wang, Jun Lv, Richard Peto, Robin Walters.

**International Co-ordinating Centre, Oxford:** Daniel Avery, Ruth Boxall, Derrick Bennett, Ka Hung Chan, Yumei Chang, Yiping Chen, Zhengming Chen, Robert Clarke, Huaidong Du, Zammy Fairhurst-Hunter, Wei Gan, Simon Gilbert, Alex Hacker, Parisa Hariri, Mike Hill, Michael Holmes, Pek Kei Im, Andri Iona, Maria Kakkoura, Christiana Kartsonaki, Rene Kerosi, Kuang Lin, Iona Millwood, Qunhua Nie, Alfred Pozarickij, Paul Ryder, Sam Sansome, Dan Schmidt, Paul Sherliker, Rajani Sohoni, Becky Stevens, Iain Turnbull, Robin Walters, Lin Wang, Neil Wright, Ling Yang, Xiaoming Yang, Pang Yao.

**National Co-ordinating Centre, Beijing:** Yu Guo, Xiao Han, Can Hou, Chun Li, Chao Liu, Jun Lv, Pei Pei, Canqing Yu.

**10 Regional Co-ordinating Centres:** **Guangxi** Provincial CDC: Naying Chen, Duo Liu, Zhenzhu Tang. Liuzhou CDC: Ningyu Chen, Qilian Jiang, Jian Lan, Mingqiang Li, Yun Liu, Fanwen Meng, Jinhuai Meng, Rong Pan, Yulu Qin, Ping Wang, Sisi Wang, Liuping Wei, Liyuan Zhou. **Gansu** Provincial CDC: Caixia Dong, Pengfei Ge, Xiaolan Ren. Maiji CDC: Zhongxiao Li, Enke Mao, Tao Wang, Hui Zhang, Xi Zhang. **Hainan** Provincial CDC: Jinyan Chen, Ximin Hu, Xiaohuan Wang. Meilan CDC: Zhendong Guo, Huimei Li, Yilei Li, Min Weng, Shukuan Wu. **Heilongjiang** Provincial CDC: Shichun Yan, Mingyuan Zou, Xue Zhou. Nangang CDC: Ziyan Guo, Quan Kang, Yanjie Li, Bo Yu, Qinai Xu. **Henan** Provincial CDC: Liang Chang, Lei Fan, Shixian Feng, Ding Zhang, Gang Zhou. Huixian CDC: Yulian Gao, Tianyou He, Pan He, Chen Hu, Huarong Sun, Xukui Zhang. **Hunan** Provincial CDC: Biyun Chen, Zhongxi Fu, Yuelong Huang, Huilin Liu, Qiaohua Xu, Li Yin. Liuyang CDC: Huajun Long, Xin Xu, Hao Zhang, Libo Zhang. **Jiangsu** Provincial CDC: Jian Su, Ran Tao, Ming Wu, Jie Yang, Jinyi Zhou, Yonglin Zhou. Suzhou CDC: Yihe Hu, Yujie Hua, Jianrong Jin Fang Liu, Jingchao Liu, Yan Lu, Liangcai Ma, Aiyu Tang, Jun Zhang. **Qingdao** Qingdao CDC: Liang Cheng, Ranran Du, Ruqin Gao, Feifei Li, Shanpeng Li, Yongmei Liu, Feng Ning, Zengchang Pang, Xiaohui Sun, Xiaocao Tian, Shaojie Wang, Yaoming Zhai, Hua Zhang, Licang CDC: Wei Hou, Silu Lv, Junzheng Wang. **Sichuan** Provincial CDC: Xiaofang Chen, Xianping Wu, Ningmei Zhang, Weiwei Zhou. Pengzhou CDC: Xiaofang Chen, Jianguo Li, Jiaqiu Liu, Guojin Luo, Qiang Sun, Xunfu Zhong. **Zhejiang** Provincial CDC: Weiwei Gong, Ruying Hu, Hao Wang, Meng Wan, Min Yu. Tongxiang CDC: Lingli Chen, Qijun Gu, Dongxia Pan, Chunmei Wang, Kaixu Xie, Xiaoyi Zhang.

**Acknowledgements:** The chief acknowledgment is to the participants, the project staff, and the China National Centre for Disease Control and Prevention (CDC) and its regional offices for assisting with the fieldwork. We thank Judith Mackay in Hong Kong; Yu Wang, Gonghuan Yang, Zhengfu Qiang, Lin Feng, Maigeng Zhou, Wenhua Zhao, Yan Zhang and Zheng Bian in China CDC; Lingzhi Kong, Xiucheng Yu, and Kun Li in the Chinese Ministry of Health; and Garry Lancaster, Sarah Clark, Martin Radley, Mike Hill, Hongchao Pan, and Jill Boreham at the CTSU, Oxford, for assisting with the design, planning, organization, and conduct of the study.

#### 3.2 The Social Science Genetics Association Consortium

Here we list alphabetically the co-authors of Okbay et al. (2016) and Lee et al. (2018)[5, 6]. The GWAS summary statistics for Educational Attainment are based on the contributions of these researchers.

Abdel Abdellaoui, Tarunveer S. Ahluwalia, Behrooz Z. Alizadeh, Maris Alver, Najaf Amin, John R. Attia, Jonas Bacelis, Andrew Bakshi, Yanchun Bao, Clemens Baumbach, Sebastian E. Baumeister, Jonathan P. Beauchamp, Daniel J. Benjamin, David A. Bennett, Klaus Berger, Lars Bertram, Ginevra Biino, Hans Bisgaard, Gyda Bjornsdottir, Jason D. Boardman, Dorret I. Boomsma, Ingrid B. Borecki, Peter Bowers, Patricia A. Boyle, Johannes H. Brsma, Ute Bultmann, Klaus Bønnelykke, Harry Campbell, Francesco P. Cappuccio, David Cesarini, Christopher F. Chabris, Guo-Bo Chen, David W. Clark, Maria Pina Concas, Dalton C. Conley, Francesco Cucca, Daniele Cusi, Gail Davies, Felix R. Day, Ian J. Deary, George V. Dedoussis, Panos Deloukas, Ilja Demuth, Jaime Derringer, Jun Ding, Cornelia M. van Duijn, Peter Eibich, Lewin Eisele, Niina Eklund, Valur Emilsson, Johan G. Eriksson, Tõnu Esko, David M. Evans, Jessica D. Faul, Mary F. Feitosa, Mark A. Fontana, Andreas J. Forstner, Barbara Franke, Lude Franke, Jeremy Freese, Nicholas A. Furlotte, Tessel E. Galesloot, Paolo Gasparini, Pablo V. Gejman, Christian Gieger, Ilaria Gin, Giorgia Girotto, Hans-Jörgen Grabe, Jacob Gratten, Patrick J.F. Groenen, Vilmundur Gudnason, Bjarni Gunnarsson, Richa Gupta, Leanne M. Hall, Bjarni V. Halldórsson, Kathleen Mullan Harris, Sarah E. Harris, Tamara B. Harris, Pim van der Harst, Caroline Hayward, Andrew C. Heath, Pamela Herd, David A. Hinds, Lynne J. Hocking, Edith Hofer, Wolfgang Hoffmann, Albert Hofman, Elizabeth G. Holliday, Georg Homuth, Michael A. Horan, Momoko Horikoshi, Jouke-Jan Hottenga, Jennifer E. Huffman, Elina Hyppönen,

William G. Iacono, Bo Jacobsson, Philip L. de Jager, Magnus Johannesson, Peter K. Joshi, Astan Jugessur, Marjo-Riitta Järvelin, Karl-Heinz Jöckel, Marika A. Kaakinen, Kadri Kaasik, Ioanna P. Kalafati, Stavroula Kanoni, Jaakko Kaprio, Sharon L.R. Kardia, Robert Karlsson, Liisa Keltigangas-Järvinen, Kathryn E. Kemper, Lambertus A.L.M. Kiemeney, Aaron Kleinman, Philipp D. Koellinger, Ivana Kolcic, Augustine Kong, Edward Kong, Seppo Koskinen, Aldi T. Kraja, Martin Kroh, Robert F. Krueger, Meena Kumari, Tushar Kundu, Zoltan Kutalik, Mika Kähönen, Jari Lahti, David I. Laibson, Max Lam, Claudia Langenberg, Antti Latvala, Lenore J. Launer, Maël P. Lebreton, Chanwook Lee, James J. Lee, Sven J. van der Lee, Christiaan de Leeuw, Steven F. Lehrer, Terho Lehtimäki, Todd Lencz, Douglas F. Levinson, Hui Li, Ruoxi Li, Paul Lichtenstein, Peter Lichtner, David C.M. Liewald, Penelope A. Lind, Karl-Oskar Lindgren, Richard Karlsson Linnér, Tian Liu, Anu Loukola, Jian'an Luan, Pamela A. Madden, Omeed Maghzian, Patrik K. E. Magnusson, Anil K. Malhotra, Massimo Mangino, Riccardo E. Marioni, Pedro Marques-Vidal, Jonathan Marten, Nicholas G. Martin, Matt McGue, George McMahon, Gerardus A. Meddens, S. Fleur W. Meddens, Sarah E. Medl, Christa Meisinger, Thomas Meitinger, Andres Metspalu, Michelle N. Meyer, Evelin Mihailov, Yusplitri Milaneschi, Lili Milani, Michael B. Miller, Grant W. Montgomery, Peter J. van der Most, Ronny Myhre, Reedik Mägi, Tomi Mäki-Opas, Christopher P. Nelson, Jan-Emmanuel De Neve, Tuan Anh Nguyen-Viet, Dale R. Nyholt, Aysu Okbay, Christopher Oldmeadow, William E.R. Ollier, Ken K. Ong, Sven Oskarsson, Aarno Palotie, Lavinia Paternoster, Antony Payton, Nancy L. Pedersen, Neil Pendleton, Brenda W.J.H. Penninx, Markus Perola, John R. B. Perry, Tune H. Pers, Natalia Pervjakova, Katja E. Petrovic, Wouter J. Peyrot, Joseph K. Pickrell, Mario Pirastu, Nicola Pirastu, Ozren Polasek, Raymond A. Poot, David J. Porteous, Danielle Posthuma, Beate Pourcain, Christine Power, Michael A. Province, Yong Qian, Lydia Quaye, Olli Raitakari, Cornelius A. Rietveld, Susan M. Ring, Marylyn D. Ritchie, Antonietta Robino, Matthew R. Robinson, Frank J.A. van Rooij, Olga Rostapshova, Rebecca Royer, Igor Rudan, Rico Rueedi, Aldo Rustichini, Katri Räikkönen, Veikko Salomaa, Erika Salvi, Nilesh J. Samani, Antti-Pekka Sarin, David Schlessinger, Börge Schmidt, Helena Schmidt, Reinhold Schmidt, Katharina E. Schraut, Rodney J. Scott, Alan R. Sers, Jianxin Shi, Julia Sidorenko, Melissa C. Smart, Albert V. Smith, Blair H. Smith, George Davey Smith, Jennifer A. Smith, Tim D. Spector, Jan A. Staessen, Kari Stefansson, Elisabeth Steinhagen-Thiessen, Konstantin Strauch, Thorkild I.A. Sørensen, Antonio Terracciano, Alexer Teumer, Kevin Thom, Gudmar Thorleifsson, Unnur Thorsteinsdottir, A. Roy Thurik, Henning Tiemeier, Nicholas J. Timpson, Pascal N. Timshel, Martin D. Tobin, Joey W. Trampush, Joyce Y. Tung, Patrick Turley, André G. Uitterlinden, Sheila Ulivi, Simona Vaccargiu, Cristina Venturini, Shefali Setia Verma, Niek Verweij, Anna A.E. Vinkhuyzen, Peter M. Visscher, Veronique Vitart, Ronald de Vlaming, Peter Vollenweider, Judith M. Vonk, Diego Vozzi, Dragana Vuckovic, Uwe Völker, Henry Völzke, Johannes Waage, Raymond K. Walters, Erin B. Ware, Nicholas J. Wareham, Chelsea Watson, Robbee Wedow, David R. Weir, Juergen Wellmann, Harm-Jan Westra, Gonneke Willemsen, Emily A. Willoughby, James F. Wilson, Alan F. Wright, Yang Wu, Jian Yang, Jingyun Yang, Loïc Yengo, Meghan Zacher, Jing Hua Zhao, Wei Zhao, Zhili Zheng, and Zhihong Zhu.

### References

[1] Turley, P. *et al.* Multi-trait analysis of genome-wide association summary statistics using MTAG. *Nature Genetics* **50**, 229–237 (2018).

[2] Bulik-Sullivan, B. K. *et al.* LD Score regression distinguishes confounding from polygenicity in genome-wide association studies. *Nature Genetics* **47**, 291–295 (2015). URL `http://dx.doi.org/10.1038/ng.3211`.

[3] Brown, B. C., Ye, C. J., Price, A. L., Zaitlen, N. & Consortium, A. G. E. N. T. . D. Transethnic genetic-correlation estimates from summary statistics. *The American Journal of Human Genetics* **99**, 76–88 (2016).

[4] Auton, A. *et al.* A global reference for human genetic variation (2015).

[5] Okbay, A. *et al.* Genome-wide association study identifies 74 loci associated with educational attainment. *Nature* (2016).

[6] Lee, J. J. *et al.* Gene discovery and polygenic prediction from a genome-wide association study of educational attainment in 1.1 million individuals. *Nature Genetics* **50**, 1112–1121 (2018). URL `https://doi.org/10.1038/s41588-018-0147-3`.

#### Frequently Asked Questions (FAQs)

This document provides information about the study:

Turley *et al.* (2021) "Multi-Ancestry Meta-Analysis yields novel genetic discoveries and ancestry-specific associations" *bioRxiv*.

The document was prepared by Daniel J. Benjamin, Shawneequa Callier, David Laibson, Alicia Martin, and Michelle N. Meyer. It has the following sections:

1. **Background**
2. **Terminology**
3. **The MAMA Method**
4. **Results of the Application of MAMA**
5. **Social and Ethical Implications**

These FAQs accompany our paper, "Multi-Ancestry Meta-Analysis yields novel genetic discoveries and ancestry-specific associations." Most existing genetic data are from, and therefore most genetics studies are done with, people of European ancestries. Our paper reports a new method, Multi-Ancestry Meta-Analysis (MAMA), for leveraging results of existing studies in people of European ancestries to enable more research on underrepresented groups. We hope MAMA can be used in concert with more diverse data-collection efforts to help eliminate disparities in genetics research.

The paper describes our new method and the results of applying MAMA to data for several outcomes. These FAQs provide context for and describe some of the limitations of our analyses in a less technical way than the paper. They do not go into comprehensive detail about MAMA or its applications, but instead describe the overarching goals of the project, how analyses were conducted, and many limitations of the work. The practice of producing these public-facing FAQ documents was first adopted by the Social Science Genetic Association Consortium (https://www.thessgac.org/faqs) and is becoming a common practice among genetics researchers. This FAQ draws from the structure and content of others that have been previously released. For a more detailed description of the paper's analyses and technical details, we invite people to read the paper itself, posted on *bioRxiv*.

If you have questions or comments about these FAQs, please contact Patrick Turley.

### 1. Background

#### 1.1. Who conducted this study?

A large, international, interdisciplinary team of researchers. The key set of researchers were drawn from the Analytic & Translational Genetics Unit (ATGU) at Massachusetts General Hospital, the Broad Institute of MIT and Harvard, and the Social Science Genetic Association Consortium (SSGAC). Much of the preliminary analysis was done by our collaborators at the China Kadoorie Biobank and BioBank Japan.

#### 1.2. Why was this study done?

Our goal in this research was to develop a statistical method that would benefit groups of people who are underrepresented in genetics research by leveraging the massive amount of genetic research that has already been done with populations of European ancestries. We hope that this work will accelerate genetic discoveries that benefit everyone.

#### 1.3. What data were used in this study? Where did your participants come from?

The data used in these analyses are from three sources: the UK Biobank, Biobank Japan, and the China Kadoorie Biobank. The UK Biobank is a database containing biomedical information, genetic data, and survey responses for roughly 500,000 individuals living in the UK. Biobank Japan is a hospital-based biobank containing clinical and genetic data on approximately 200,000 individuals who were treated in hospitals throughout Japan. The China Kadoorie Biobank is a dataset of around 500,000 individuals (100,000 of whom have contributed genetic data) who have been surveyed and whose data has been linked to administrative health records. Ethics approval was obtained for all three data sets, and all participants gave their informed consent to have their data used in studies like this.

### 2. Terminology

#### 2.1. What is a GWAS?

A Genome-Wide Association Study (GWAS; pronounced JEE-wahs) is a systematic set of tests looking for genetic variants that are associated with a specific outcome. The study is “genome-wide” because millions of genetic variants are tested from across the whole genome.

#### 2.2. What is a MAMA?

A Multi-Ancestry Meta-Analysis (MAMA; pronounced MA-ma), as described in more detail later (see the section titled "The MAMA Method"), is a statistical method that combines GWAS results from different ancestry groups. Relative to the GWAS results, in each ancestry group, MAMA gives a more precise estimate of the association between the outcome and each genetic variant.

#### 2.3. What genetic variants are studied in GWAS (and MAMA)?

GWAS and MAMA are used to study a specific type of genetic variant called a single-nucleotide polymorphism (SNP; pronounced SNIP). Our DNA is made up of long strings (billions of pairs) of molecules called "nucleotides." If you compared the DNA sequences for any two people in the world, you'd find that 99% of their DNA consisted of the same sequence of nucleotides. However, there are many millions of positions in the sequence where some people have one type of nucleotide and others have a different type of nucleotide. These positions where people can vary by single nucleotides are called SNPs. The different nucleotides that can possibly be found at a particular SNP are called "alleles" for that SNP. Below, when we refer to a person "with a certain genetic variant," we mean a person who has a certain allele at a particular SNP.

#### 2.4. What is a meta-analysis?

Meta-analysis is a general term for combining the results of multiple studies that ask the same question (in this case, the relationship between a set of genetic variants and an outcome) in different groups of people. When many sets of GWAS results are combined in a meta-analysis, it often produces a single, more powerful study that can yield more genetic discoveries.

#### 2.5. What is meant by "association"? Why aren't GWAS associations causal?

There is an "association" between an outcome and a genetic variant if people with the genetic variant tend to have a different outcome than people without the genetic variant. For example, if people with a certain genetic variant are on average taller than those without the variant, then we would say the variant is associated with height. Or if a larger fraction of people with a genetic variant are diagnosed with schizophrenia than those without a genetic variant, we would say that the variant is associated with a schizophrenia diagnosis.

When we find that a genetic variant is associated with an outcome, for several reasons, that does not necessarily mean that the variant causes the outcome. First, it could be that certain genetic variants are by chance more common in some geographic or ancestral groups. If those groups also face different environments that influence the outcome (e.g., pollution, racism, or social programs), then those variants would be associated with the outcome. This is not because the allele causes the differences in the outcome; rather, it reflects the *environmental* channels that are correlated with the variant. For example, in places where those with darker skin are discriminated against, a genetic variant associated with skin pigmentation may be associated with a variety of worse health and/or other outcomes—but the colorism (i.e., discrimination against those with darker skin tone), not the genetic variant, is likely the direct cause of the worse outcomes.

Second, GWAS cannot distinguish between correlated genetic variants. If a certain genetic variant is correlated with another genetic variant (see the section titled "What is linkage disequilibrium?"), and only one variant influences an outcome, this will cause both variants to be associated with the outcome.

Finally, in some cases, genetic variants may only be associated with the outcome in certain settings. For example, a variant associated with better memory may therefore also be associated with more years of education in a society where school emphasizes memorization. If a different society (or the same society at a different time) instead chose to emphasize experiential learning, that variant might still be associated with better memory but be much less strongly associated with years of education (or no longer associated with that outcome at all).

In sum, generally, GWAS can only tell us *if* a genetic variant is associated with an outcome in a given population and at a given time and place; it cannot tell us *why* the variant is associated, and it therefore does not demonstrate that the relationship is causal or that it is likely to hold up in other populations, times, or places.

So if GWAS can't tell us if a SNP causes a change in an outcome and it can't even tell us if it will be associated with the outcome in the future, why do researchers conduct GWAS? The main reason is that results from GWAS help guide researchers in their future work. For example, testing drugs that may treat disease is a very expensive process, and only one out of ten drugs that get tested in this process is effective enough to get approved. However, drugs that are based on genetic evidence (including evidence from GWAS) are twice as likely to be effective and get approved compared to drugs that are not developed with genetic evidence (King, Davis, and Degner, 2019). This means that, using GWAS results, we are able to find treatments for disease more quickly and for less money. In addition to helping develop treatments for disease, GWAS results can be a useful input for identifying the causal variants. (see the section titled "What can be done with the results of research using MAMA? What are the potential benefits of this research?"). GWAS results can also help us make genetic predictors that are useful in a wide variety of research applications and may eventually be clinically useful, e.g., as a way to identify patients who should get additional screening for certain diseases.

#### 2.6. What is ancestry?

When genetics researchers refer to "ancestry" they almost always mean "genetic ancestry." As explained below, genetic ancestry is not the same as race or ethnicity.

In theory, when genetics researchers talk about individuals with genetic ancestries from a particular region (e.g., East Asian ancestries or European ancestries), they mean that these individuals inherited a substantial fraction of their genetic variants from a hypothetical group of closely-related individuals who lived in that region hundreds or thousands of years ago. However, because populations have always been mobile, and many individuals have offspring with partners outside the region where they grew up, everyone has inherited genetic variants from people who lived in multiple, different places hundreds or thousands of years ago. Thus, ancestry is actually a continuum, with each individual having some mixture of ancestries from several generations ago (and each of these ancestries themselves being a mixture of earlier ancestries). Any grouping of ancestries into discrete categories is an approximation of the much more complicated reality. Researchers make these approximations—using concepts of ancestral groups such as "East Asian"—when they want to study individuals who are likely to have a certain amount of shared genetic ancestry (see the section titled "Why are different ancestry groups usually studied in separate GWAS? Why are certain individuals normally excluded altogether from GWAS?").

In practice, when researchers assign a person to a genetic ancestry group for a particular region, they mean that the person is genetically related to (i.e., the person shares many genetic variants with) a *modern* reference group of individuals who are *thought* to trace their family tree to that region for a few generations. This means that, when assigning ancestry, researchers must make three decisions: (1) whom they will include in their reference group, (2) how they will measure genetic relatedness, and (3) how closely related a person must be to the reference group to be assigned to the same ancestry group. These decisions are not straightforward because human migratory history is extraordinarily complicated. As a result, there are loose rules of thumb that researchers and data providers use in making these decisions, and there are still many differences in how these decisions are made across studies. Because of this, the same person could be assigned into different ancestry groups, depending on the decisions of a particular researcher or data provider.

Again, and to elaborate, ancestry should not be confused with "race" or "ethnicity." Race and ethnicity group people based on perceived shared physical characteristics or on cultural, national, linguistic, religious, or other social affiliations. Racial and ethnic categories, and the definitions that are used to categorize people, are different in different times and places. In many cases, racial and ethnic categories are not closely related to genetic ancestry. For example, consider two people whose respective family trees can be traced for several generations to South Africa and to Kenya. Both would likely be perceived as Black in the US today. However, the current, non-immigrant populations of Kenya and South Africa are at least as distantly genetically related to each other as the current, non-immigrant populations of England and Japan. Therefore, in most research studies, researchers would classify these two people as having distinct genetic ancestries.

Treating genetic ancestry, ethnicity, and race as equivalent concepts leads to scientific confusion and can harm disadvantaged populations by inviting people to believe—*incorrectly*—that current disparities among people of different races or ethnicities are rooted in biology. See the section titled "Do genes determine the choices we make and who we are?".

#### 2.7. Why are different ancestry groups usually studied in separate GWAS? Why are certain individuals normally excluded altogether from GWAS?

Different ancestry groups are studied separately and certain individuals are excluded in GWAS for statistical reasons. Using current GWAS methods, studying all available individuals in a single analysis could produce results that overwhelmingly represent environmental factors instead of identifying promising genetic variants for the future study of biological mechanisms.

A commonly used example is a hypothetical GWAS of chopstick use (Hamer and Sirota, 2000). Genetic variants are, *by chance*, more and less common in different populations. If a GWAS of chopstick use were conducted in a sample that includes some people with East Asian ancestries and some people with non-East Asian ancestries, we would almost surely find that many genetic variants that are more or less common among those of East Asian ancestries would be associated with chopstick use. But these associations would not likely reflect biological mechanisms that affect manual dexterity or a personal preference for wooden cutlery. Rather, these associations would be purely driven by environmental/cultural differences in where and how a person is raised.

When GWAS is conducted in a group of individuals with similar ancestries (that is, a group of people who are all genetically related to each other), associations are less likely to be driven by these types of non-causal factors, which makes the results easier to interpret and makes follow-up work more productive. Because of this, researchers generally restrict the people they study to a group whose members are sufficiently genetically related to each other, and omit the others. Because a large group is required to conduct a statistically sound GWAS, small groups of individuals cannot be studied until many additional individuals who are genetically related to them are also recruited to contribute genetic data. As more data are gathered and more methods are developed, it will eventually no longer be necessary to omit individuals due to their genetic ancestry.

#### 2.8. What about people with ancestries from multiple groups?

As described above (see the section titled "What is ancestry?"), genetic ancestry is a continuum. Many people may be genetically related to multiple groups from several regions.

For example, suppose the ancestry groups we are considering are French and German. Then our reference groups that define these ancestries might be individuals currently living in, and whose families have for generations lived in, Paris and Berlin, respectively. In that case, being categorized as having "French ancestry" really means being more similar genetically to the Parisians in our reference group than to the Berliners in our reference group. Historically, people have tended to live and find mates near where they grew up. Consequently, individuals who live farther from Paris and closer to Berlin (regardless of which side of the national border they live on) are likely on average to look, genetically, "in between" French and German because some of their ancestors have descendents who now live in Paris and some of their ancestors have descendents who now live in Berlin.

As is clear from the above example, if we define our groups narrowly enough, almost everyone could be considered to have "mixed" ancestry. However, in GWAS, if the different ancestry groups are made up of people who are closely genetically related (like they would be in our French/German example), it is possible to measure and account for the genetic differences between the groups and for how much ancestry a person has from each group. As discussed above (see the section titled "Why are different ancestry groups usually studied in separate GWAS? Why are certain individuals normally excluded altogether from GWAS?"), GWAS runs into problems when some of the people studied are distantly genetically related to other people studied (e.g., those with European vs East Asian ancestries or with West African vs South African ancestries).

Because of this, there is one case of "mixed" ancestry that presents a problem for GWAS: when people have *recent* ancestors (within the last 3 or 4 generations) from very distantly related genetic ancestries. In this case (often called "recent admixture"), large segments of the person's DNA will be from one of the ancestry groups but not the other, while it will be vice-versa for other large segments of the person's DNA. Standard statistical tools for genetic studies (including MAMA) cannot account for this situation. As a result, such individuals are generally not included in GWAS. However, research on how to incorporate such individuals into GWAS is an important and quickly advancing area. We are hopeful that as more data for diverse-ancestry GWAS are collected and as more robust statistical tools are developed, genomics research can also become more representative and beneficial for all people.

##### 2.9. Why can't you learn anything about biological differences between ancestry groups from GWAS associations?

As discussed above (see the section titled "What is meant by "association"? Why aren't GWAS associations causal?"), GWAS associations are not necessarily causal and depend on the environment of the individuals included in the GWAS. Different ancestry groups arise in the world because they became partially separated from each other many generations ago, for example, due to geographic barriers or social forces. When two groups are geographically or socially separated, genetic variants can by chance become more or less common in each group. The groups also face different environments. Those environments not only may have direct effects on certain outcomes (such as disease risk) but may also change the effects of specific genetic variants (see the section titled "What is meant by "association"? Why aren't GWAS associations causal?"). Therefore, when individuals from two ancestry groups have different average outcomes, it is impossible to conclude from GWAS whether the difference is due to average genetic differences between the groups or to the different environments faced by the groups. For this reason, **it is scientifically invalid to make general statements about ancestry group differences based on GWAS associations.**

##### 2.10. What is linkage disequilibrium?

Linkage disequilibrium refers to the correlation across genetic variants: when people have one genetic variant, they are more likely to also have other genetic variants that are physically nearby on the same string of DNA. The strength of the correlations between nearby variants changes over time due to the random ways that genes are inherited from generation to generation. As a result, the pattern of linkage disequilibrium differs between ancestry groups, especially between ancestry groups that are distantly related.

#### 3. The MAMA Method

##### 3.1. What is the goal of the MAMA method?

The MAMA method was developed in order to improve scientists' ability to make genetic discoveries in groups that have historically been underrepresented in genetics research. Today, even though individuals who would be classified as having European ancestries are a small fraction of the global population, they make up the vast majority of the individuals studied in GWAS. As a result, most genetic knowledge we have acquired from these studies can only be reliably applied to European-ancestry individuals. Because genetic knowledge can help improve medical care, the disparity in knowledge could potentially exacerbate health disparities that already exist in society.

MAMA improves our ability to gain genetic knowledge about non-European ancestry groups. More precisely, it is a statistical method that leverages information from large GWAS in overrepresented populations to boost the power for making genetic discoveries in GWAS of underrepresented ones. MAMA is not a replacement for the important efforts of diverse data collection, but we hope it can be used in concert with such efforts to help eliminate disparities in genetics research.

#### 3.2. What do the results of MAMA tell you?

Like the results of a GWAS study, the results of a MAMA study are estimates of the associations between an outcome of interest and a large set of genetic variants throughout the genome. As such, MAMA associations have all the same strengths and limitations of GWAS associations. For example, MAMA associations may not imply a genetic variant has a causal impact on the outcome (see the section titled "What is meant by "association"? Why aren't GWAS associations causal?"), and MAMA associations cannot be compared across ancestry groups (see the section titled "Why can't you learn anything about biological differences between ancestry groups from GWAS associations?"). However, like GWAS associations, MAMA associations provide clues to genetics researchers about where they should prioritize future studies of human health and behavior.

#### 3.3. What does MAMA do, and how?

MAMA takes as its inputs results from GWAS conducted in two or more ancestry groups. In our applications in the paper, these inputs are results from GWAS in European-ancestry samples and from GWAS in East-Asian-ancestry samples. The purpose of MAMA is to increase power to identify genetic associations in one ancestry group (in our case, East Asian) by leveraging the information in another ancestry group or groups (in our case, European).

There are two reasons why GWAS results from one ancestry group are not directly comparable to, and therefore cannot simply be combined with, GWAS results from another (distantly related) ancestry group. First, recall that linkage disequilibrium patterns differ across distantly related ancestry groups. Ignoring these differences would lead to incorrect conclusions for either ancestry group if we directly combined the results using standard meta-analysis methods (see the section titled "What is a meta-analysis?"). Second, for some outcomes—especially behavioral and social outcomes such as educational attainment—it is likely that some genetic variants have stronger or weaker associations with the outcome in different ancestry groups due to the different environments experienced by individuals in those groups (see the section titled "What is meant by "association"? Why aren't GWAS associations causal?"). For this reason, too, standard methods of combining the results would lead to biased conclusions for either ancestry group.

MAMA overcomes both of these obstacles. It combines the results across ancestry groups in a way that addresses the differences across the groups, and it delivers estimates of each genetic variant's association with the outcome in *each* of the ancestry groups included in the analysis. In contrast to association estimates from a standard meta-analysis, MAMA estimates will be correct on average. MAMA overcomes the first obstacle by accounting for the fact that genetic associations are likely to be less similar across ancestry groups in places in the genome where the linkage disequilibrium patterns are less similar across the ancestry groups. To overcome the second obstacle, MAMA conducts an analysis of the GWAS results from the different ancestry groups and estimates the overall similarity of the genetic associations. It then uses this estimate of overall similarity to help determine how similar the associations are likely to be at any specific genetic variant. For every specific genetic variant, using this information on the overall similarity of the genetic associations across ancestry groups and the linkage disequilibrium patterns near the variant, MAMA appropriately incorporates the GWAS results from each ancestry group into the GWAS results of the other ancestry group(s).

##### 3.4. Why can't you learn anything about biological differences between ancestry groups from MAMA associations?

MAMA associations have the same interpretation as GWAS associations. As a result, the same reasons why it is scientifically invalid to draw conclusions about biological differences between ancestry groups with GWAS associations apply to MAMA associations as well (see the section titled "Why can't you learn anything about biological differences between ancestry groups from GWAS associations?").

#### 4. Results of the Application of MAMA

##### 4.1. What was done?

To illustrate the MAMA method, we applied MAMA to GWAS associations based on European-ancestry and East-Asian-ancestry populations for 23 outcomes. After identifying associations in both populations, we verified that these associations could also be found using independent replication samples drawn from the same populations.

##### 4.2. What outcomes did you study?

The outcomes studied in our paper were chosen because they are of ongoing interest in genetics research and because we could study them in the datasets that were available to us. They include anthropometric outcomes (e.g., height and BMI), biomarker outcomes (e.g., white blood cell count and hemoglobin), and behavioral outcomes (e.g., educational attainment and physical activity). We hope that other researchers will find MAMA useful in studying these and many other outcomes in the future.

#### 4.3. How did you decide which ancestry groups to include in your application of MAMA?

In the applications of MAMA in our paper, we only use GWAS associations based on samples with European ancestries and East-Asian ancestries. We study samples of European ancestries because these are the largest existing GWAS samples, and we want to investigate whether information from GWAS in European-ancestry samples can be leveraged to learn about genetic associations in other ancestry groups. We study samples of East-Asian ancestries because, although much smaller than the European-ancestry samples, they are large enough that for many of the outcomes, GWAS can reliably identify some genetic variants even when conducted only in the East-Asian-ancestry samples. While that is not important for MAMA to work—and, indeed, MAMA may prove most useful when applied to other ancestry groups—it is valuable for our purpose in this paper of validating the MAMA method. By studying samples of East-Asian ancestry, we can show that MAMA identifies genetic variants that could not be identified in the European-ancestry or East-Asian-ancestry samples analyzed separately, as well as that these associations replicate in an independent sample of East-Asian-ancestry individuals.

#### 4.4. What did you learn by applying MAMA to several outcomes?

Applying MAMA, we are able to identify hundreds of independent genetic variants that are associated with various outcomes. Many of the associated variants we identified would not have been identified in either of the set of European or East-Asian ancestry GWAS associations alone. When we compare our associated variants from MAMA to association estimates from an independent sample, the associations tend to be in the same direction more often than would be expected by chance. This implies that at least some of the MAMA associations represent real associations in the corresponding population. Also, mathematical theory and computer simulations give us further confidence that MAMA is generating valid estimates for each ancestry group.

In the figure below, we give an example of how MAMA helps make new genetic discoveries when applied to GWAS results for BMI. We begin with two sets of GWAS results: one based on 350,011 European-ancestry people from the UK and one based on 188,613 East-Asian-ancestry people from Japan and China. The heights of the green bars (on the left) report how many unique genetic variants are identified in each set of GWAS results. Unsurprisingly, we find more associated genetic variants in the European-based population since we have many more European-ancestry people in our data. However, when we use MAMA—as shown by the total height of the blue bar (on the right)—we find even more associated genetic variants for our East-Asian-ancestry group than we found in the GWAS results from the (larger) European-ancestry group. Some of these variants are the same as those already found by the original GWAS with European or East Asian samples, but roughly a quarter of the variants found by MAMA are novel, meaning they are not included in either of the two green bars.

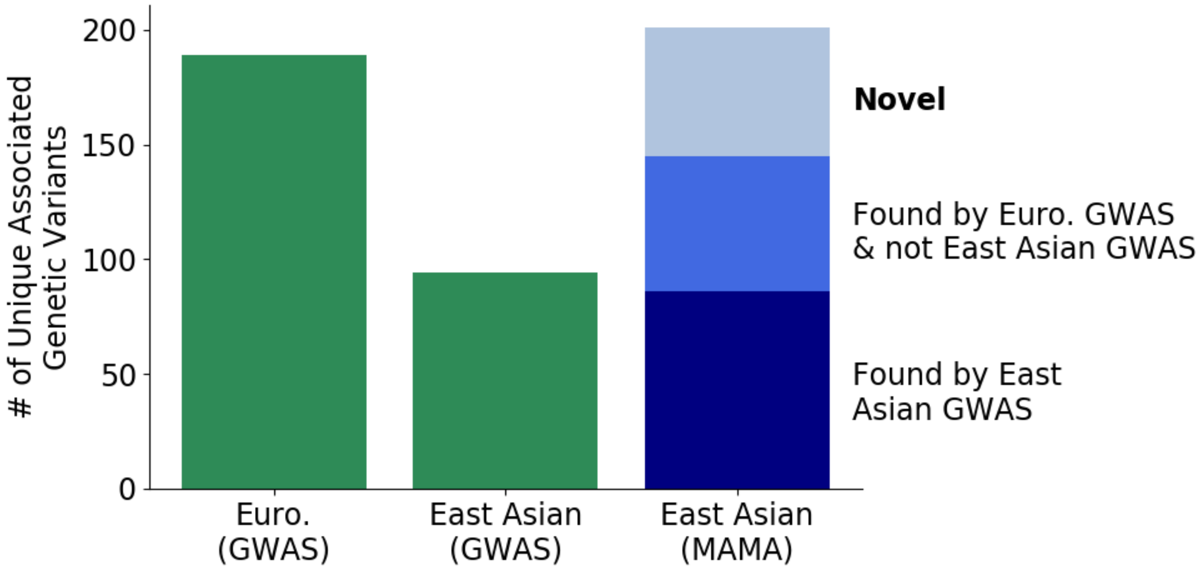

#### 4.5. Can you use MAMA results to make genetic predictors?

Yes, but there are other tools that are designed to make genetic predictors that are more powerful than predictors based on MAMA results.

MAMA was designed as a method to improve how precisely we can estimate the associations between genetic variants and outcomes in specific ancestry groups. This is a useful feature for making genetic predictors, but it is not the only important feature. As a result, genetic predictors based on MAMA results for an ancestry group will be more powerful than predictors based on just the GWAS results for the ancestry group, but other approaches that are designed specifically to make genetic predictors when you have GWAS results from multiple ancestry groups will do better.

### 5. Social and Ethical Implications

#### 5.1. What can be done with the results of research using MAMA? What are the potential benefits of this research?

Incorporating data on diverse populations in genetics research benefits everyone, and may have the greatest benefits for currently underrepresented populations.

Perhaps the most notable example of how greater diversity in genetics research can benefit everyone is 'fine-mapping,' which uses statistical methods to attempt to identify which variant is responsible for an observed association in a GWAS analysis. It can similarly be applied to results from a MAMA analysis. When a genetic variant is identified by (GWAS or) MAMA, it could be because of that specific genetic variant, but it is more often due to a nearby genetic variant that is correlated with it (i.e., in linkage disequilibrium with it; see the section titled "What is linkage disequilibrium?"). In different ancestry groups, the patterns of correlation among genetic variants differ. Fine-mapping exploits these differences in the correlation patterns to pinpoint the genetic variant responsible for an association.

For example, suppose MAMA finds an association with genetic variant A in one ancestry group, and genetic variant A is correlated with genetic variant B in that ancestry group. And suppose MAMA finds an association with genetic variant C in another ancestry group, and in this ancestry group, genetic variant C is correlated with genetic variant B. Putting these two findings together provides evidence that genetic variant B is truly responsible for both findings.

By pinpointing more precisely which variants to examine in follow-up laboratory studies, fine-mapping accelerates the process of linking genetic associations to potential biological pathways and the development of potential pharmacological therapies. One example of leveraging multi-ancestral populations to better determine causal gene sets is in a publication on inflammatory bowel disease (Huang et al., 2017). By using a combination of European and East Asian cohorts, researchers were able to refine their GWAS signals to specific variants with high certainty. This allowed them to link inflammatory bowel disease to specific immune cells and gut mucosa. Including diverse populations thus allows improved determination of strong genetic factors and pathways for disease that can be targeted therapeutically, both for the underrepresented groups themselves, but more generally for people of all ancestries.

In order to carry out fine-mapping as described above, researchers need precise estimates of the associations between the outcome they are interested in and many genetic variants. MAMA makes association estimates from a GWAS more precise, which makes fine-mapping more likely to succeed.

GWAS and MAMA results are also useful for identifying tissues and biological systems that are important for certain outcomes. For example, a recent analysis of GWAS results for schizophrenia found that genetic variants associated with schizophrenia are concentrated in regions of DNA related to neurons but not in regions related to other types of brain cells (Skene et al., 2018). This finding helps focus future schizophrenia research on the types of cells that are most important.

#### 5.2. Are there additional benefits and risks to studying behavioral outcomes?

This study includes results for a large number of outcomes for which we could conduct a well-powered GWAS in the UK Biobank and either the China Kadoorie Biobank or Biobank Japan. This included several behavioral outcomes that were collected by the UK Biobank and the China Kadoorie Biobank (including age at first birth, educational attainment, physical activity, and smoking initiation).

Several researchers, including some involved in this project, have also conducted GWAS of behavioral outcomes in populations with only European ancestries. Behavioral research on these topics is particularly sensitive. We recommend that interested readers should also read the FAQs for those papers. Those FAQs go into greater detail on the interpretation and social and ethical implications of studying the genetics of behavior: https://www.thessgac.org/faqs.

#### 5.3. Do genes determine the choices we make and who we are?

No. Genes and genetic variation do not determine our choices or who we become.

If they did, identical twins would make all of the same decisions, have the same interests, etc. Years of twin studies have shown that, while identical twins tend to be more similar than fraternal twins, they are nevertheless different. This implies that environmental factors also play a large role in our outcomes. Even for outcomes that are strongly associated with some genetic variants, the associations may not represent causal mechanisms. And even when associations represent causal relationships, these causal pathways are complicated and interact with the environment.

The fact the genes and environments interact has two important implications. First, genetic variants that influence an outcome in one setting might not in another setting. For example, suppose a major pandemic caused countries to switch to remote schooling. In such a scenario, the genetic influences that are related to academic achievement in the pandemic regime may be different than the influences that are related to academic achievement during in-person schooling. Second, the environment can alter or even wipe out what would have been the effect of genetic variants, which means that genes aren't destiny. For example, someone genetically predisposed towards being taller than average might end up being shorter than average if they lacked adequate nutrition during childhood. Likewise, someone genetically predisposed to have poor eyesight can have perfect eyesight through the use of glasses, contact lenses, or surgery.

#### 5.4. Should clinicians or other practitioners use MAMA or its results to make decisions?

No. GWAS and MAMA associations alone are not actionable for decisions being made by clinicians and other practitioners. They are only an important first step in basic science research that might someday be useful in helping clinicians and practitioners make decisions. Like GWAS, MAMA can help identify genetic variants associated with an outcome of interest. Subsequent basic research studies of those variants would then be needed to confirm their relationship to the outcome. If successful, "translational" research might then proceed to developing interventions to improve outcomes. For example, if the intervention is a drug, the translational research might involve preclinical animal studies and a series of phased human trials before the intervention would possibly be ready to be used in practice.

#### 5.5. How has genetics research been used to harm different groups?

Genetics research has a historical legacy of racist and classist inferences that have harmed underrepresented groups.

Indeed, the term "eugenics" was coined in the late 1800s by one of most prominent early researchers of heredity, Francis Galton. In the first half of the 20th century, Ronald Fisher, who was one of the most influential geneticists and statisticians in the history of science, was an active proponent of the belief that socioeconomic disparities in society were primarily caused by genetic factors. The racist and classist attitudes endorsed by Galton, Fisher, and many others in the scientific community laid the groundwork for 20th century forced sterilizations, anti-miscegenation laws, immigration restrictions, and even genocide. Genetics research also has the potential to stigmatize groups. Today, racist individuals and groups continue to misinterpret the results of genetics research to justify their claims.

Acknowledging the harm done to certain groups in the name of science reminds us of the importance of careful communication of the implications of scientific research and the need for intense vigilance to ensure that disadvantaged groups are not further harmed by this and related work.

#### 5.6. Could MAMA and its applications be used to harm certain groups (e.g., through discrimination or stigmatization)?

Possibly, but excluding these groups from genetics research is also harmful.

As described above, genetics has a long history of being used to stigmatize certain groups. Although our results do not imply that phenotypic differences between groups are due to genetic or biological differences, we worry that some individuals could mistakenly or willfully misinterpret our study to advance racist ideologies. However, the exclusion of diverse groups from genetics research can lead to inequity in scientific knowledge and to underrepresented populations not benefitting from that knowledge. In our view, the best way to proceed is to conduct and promote more diverse genetics research, while mitigating potential harms. We have carried out a number of activities in an effort to maximize benefits and minimize risks from this research. See our response in the next section, “What has been done to reduce the potential harms of this research?”, for more information.

#### 5.7. What has been done to reduce the potential harms of this research?

We have adopted several strategies to reduce the potential harms of this research.

First, we have written this FAQ to help the public and journalists understand the value and the many limitations of this work. We welcome feedback from the community if any aspect of our analyses and interpretations remains unclear.

Second, we also discussed this project and this FAQ with groups and individuals with diverse expertise and perspectives. These included two co-authors on this paper: Shawneequa Callier, a bioethics professor who specializes in the ethical, legal, and social implications of genetics research and racial categories, and Michelle Meyer, an assistant professor at Geisinger Health System who specializes in research ethics and communication. Professors Callier and Meyer both provided feedback on this FAQ and the considerations described in it. We also met during early stages of the MAMA project with members of Shades@Broad, an identity-based affinity group whose mission is to advocate for and support the recruitment, development, and success of ethnic minorities at the Broad. In these meetings, group members provided input on how the work was carried out and communicated. Several of their questions and concerns are directly addressed in this FAQ.

Third, we are developing a Terms of Use for researchers who would like to use the associations we have estimated with MAMA in their own research. Researchers will agree to “conduct research that strictly adheres to the principles articulated by the American Society of Human Genetics (ASHG) position statement: ASHG Denounces Attempts to Link Genetics and Racial Supremacy. (See also International Genetic Epidemiological Society Statement on Racism and Genetic Epidemiology.).” Data-users also will agree to “not use these data to make comparisons across ancestral groups.”

Finally, even with these efforts, it is still likely that some will mistakenly or purposefully misinterpret this work. As such occasions arise, we will attempt to correct the public record.

### 6. References

Hamer, D. H., & Sirota, R. (2000). Beware the chopsticks gene. *Molecular psychiatry, 5*(1), 11-13.

Huang, H., Fang, M., Jostins, L., Mirkov, M. U., Boucher, G., Anderson, C. A., . . . Crins, F. (2017). Fine-mapping inflammatory bowel disease loci to single-variant resolution. *Nature, 547*(7662), 173-178.

King, E. A., Davis, J. W., & Degner, J. F. (2019). Are drug targets with genetic support twice as likely to be approved? Revised estimates of the impact of genetic support for drug mechanisms on the probability of drug approval. *PLoS genetics, 15*(12), e1008489.

Skene, N. G., Bryois, J., Bakken, T. E., Breen, G., Crowley, J. J., Gaspar, H. A., . . . Muñoz-Manchado, A. B. (2018). Genetic identification of brain cell types underlying schizophrenia. *Nature genetics, 50*(6), 825-833.
