## Supplementary Figures for "Multi-Ancestry Meta-Analysis yields novel genetic discoveries and ancestry-specific associations"

Tables and Figures

Figure S1. Bias Simulation in Three Populations.

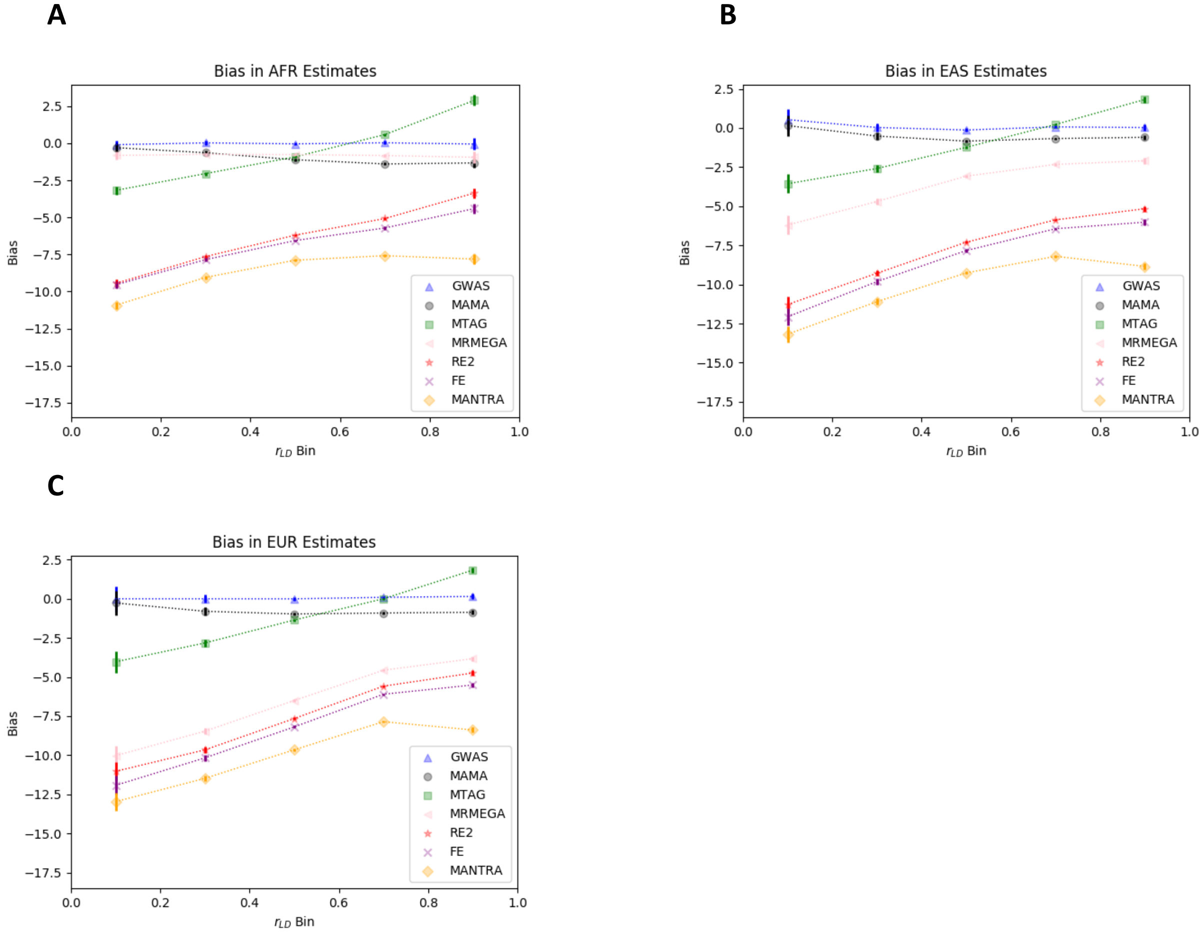


Note: Bias estimates in (a) AFR, (b) EAS, and (c) EUR, defined as the difference between marginal SNP effects and meta-analyzed SNP effects averaged within LD correlation bins. All SNPs are oriented such that their marginal effect is positive. 95% confidence intervals are represented by vertical bars. SNPs are binned into 5 groups of equal width between zero and one, and the bias is reported for SNPs within a bin.

Figure S2. Mean χ$^2$ Simulation in Three Populations.

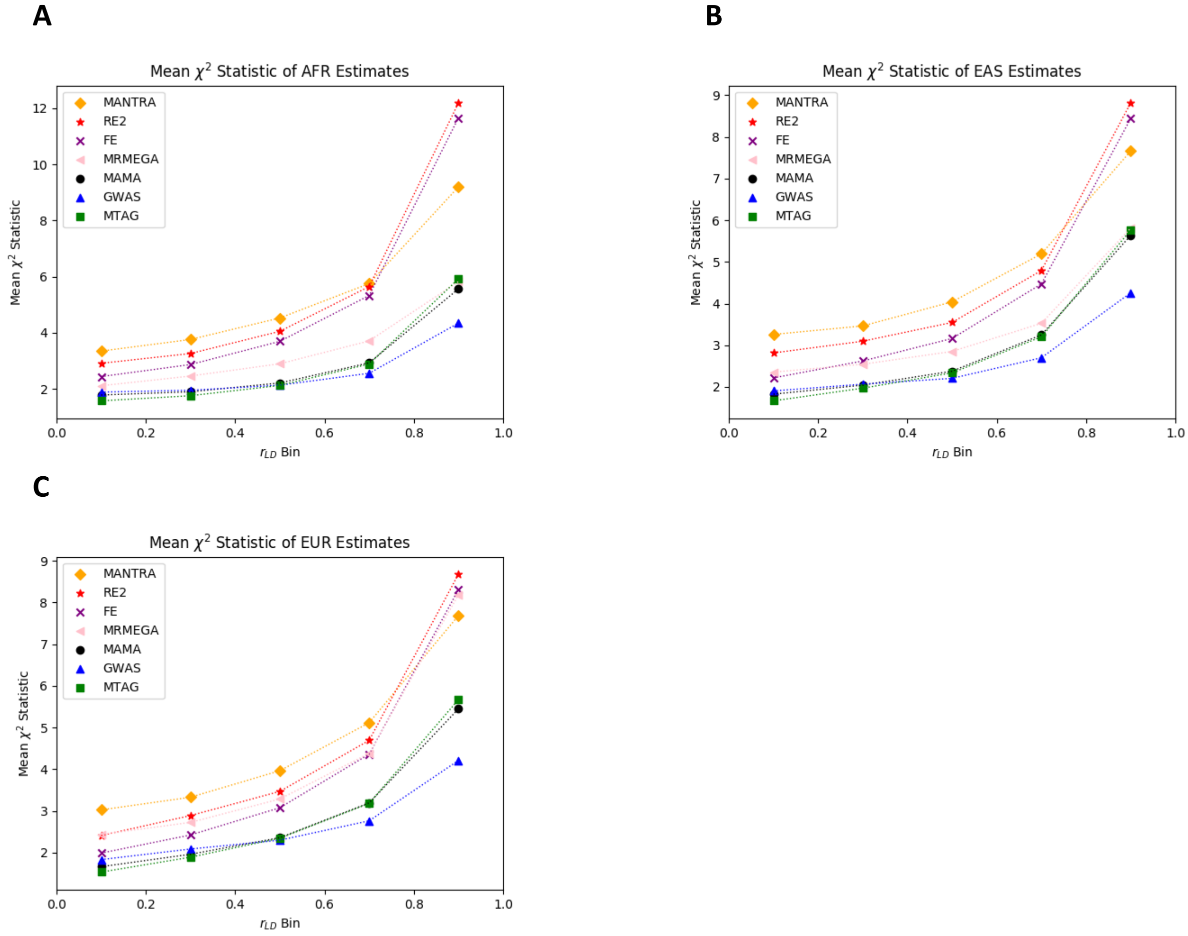


Note: Mean $\chi^2$ estimates in (a) AFR, (b) EAS, and (c) EUR, defined as the difference between marginal SNP effects and meta-analyzed SNP effects averaged within LD correlation bins RE2 mean $\chi^2$ uses the reported RE2 p-value and evaluates it on the inverse $\chi^2$ distribution with one degree of freedom. SNPs are binned into 5 groups of equal width between zero and one, and the mean $\chi^2$ estimate is reported for SNPs within a bin.

Figure S3. Type 1 Error Simulation in Three Populations.

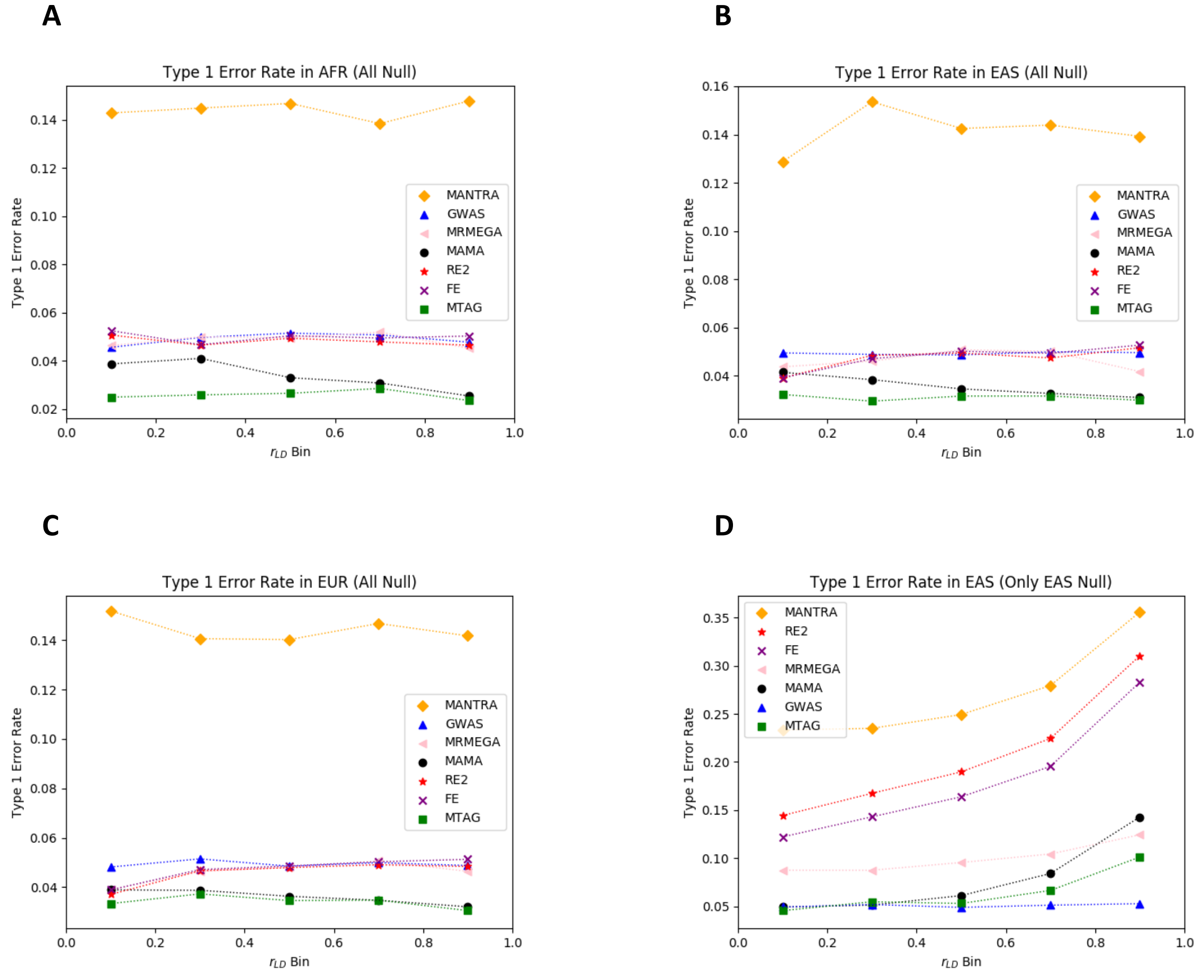


Note: Type 1 error rate in (a) AFR, (b) EAS, and (c) EUR for SNPs that are null in all three populations. Panel (d) reports the Type 1 Error Rate for the EAS population for SNPs that are null in the EAS population but nonnull in the AFR and EUR population. Type 1 error rate is defined as the fraction of null SNPs whose *P* value is less than 0.05. SNPs are binned into 5 groups of equal width between zero and one, and the Type 1 Error Rate is reported for SNPs within a bin.

Figure S4. Bias Simulation in Two Populations.

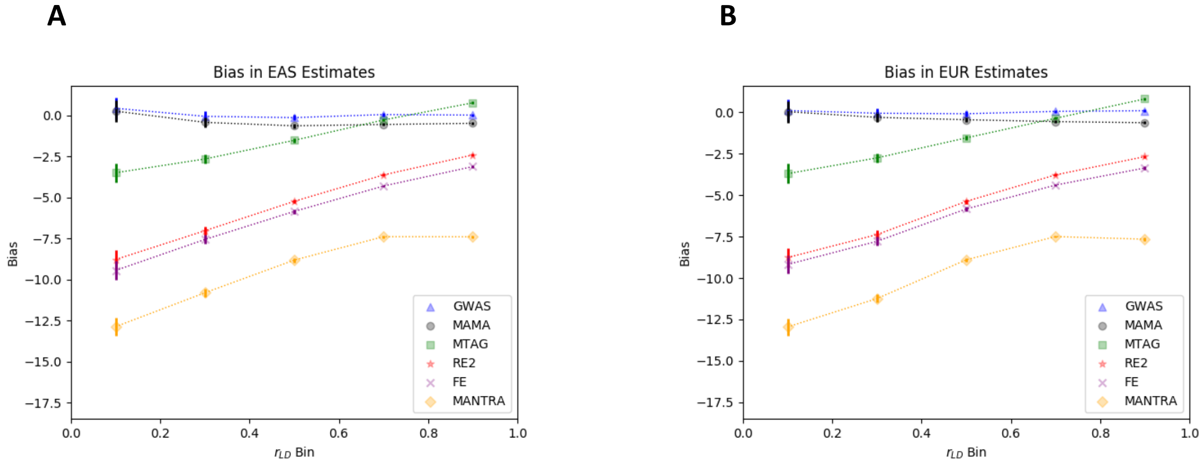


Note: Bias estimates in (a) EAS and (b) EUR, defined as the difference between marginal SNP effects and meta-analyzed SNP effects averaged within LD correlation bins. All SNPs are oriented such that their marginal effect is positive. 95% confidence intervals are represented by vertical bars. SNPs are binned into 5 groups of equal width between zero and one, and the bias is reported for SNPs within a bin.

Figure S5. Mean $\chi^2$ Simulation in Two Populations.

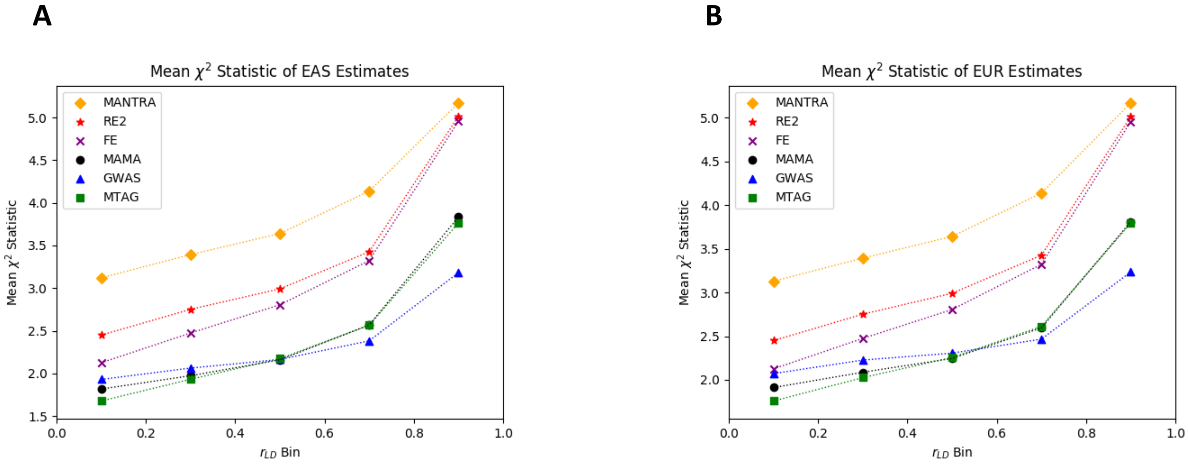


Note: Mean $\chi^2$ estimates in (a) EAS and (b) EUR, defined as the squared ratio of the meta-analyzed SNP effect over the SNP's standard error averaged within LD correlation bins. RE2 mean $\chi^2$ uses the reported RE2 p-value and evaluates it on the inverse $\chi^2$ distribution with one degree of freedom. SNPs are binned into 5 groups of equal width between zero and one, and the mean $\chi^2$ estimate is reported for SNPs within a bin.

Figure S6: Type 1 Error Simulation in Two Populations.

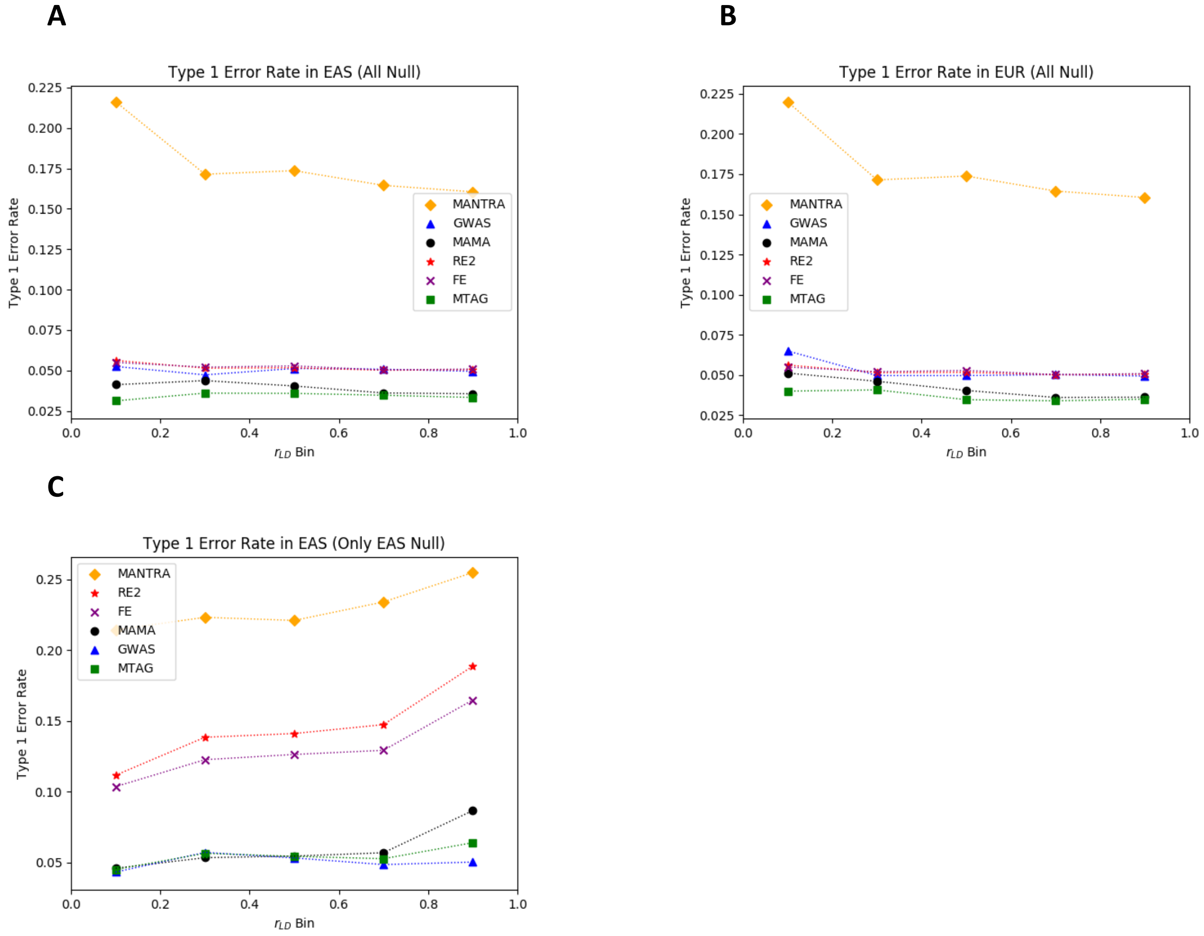


Note: Type 1 error rate in (a) EAS and (b) EUR for SNPs that are null in both populations. Panel (c) reports the Type 1 Error Rate for the EAS population for SNPs that are null in the EAS population but nonnull in the EUR population. Type 1 error rate is defined as the fraction of null SNPs whose *P* value is less than 0.05. SNPs are binned into 5 groups of equal width between zero and one, and the Type 1 Error Rate is reported for SNPs within a bin.

Figure S7. GWAS and MAMA Manhattan Plots for Age at First Birth

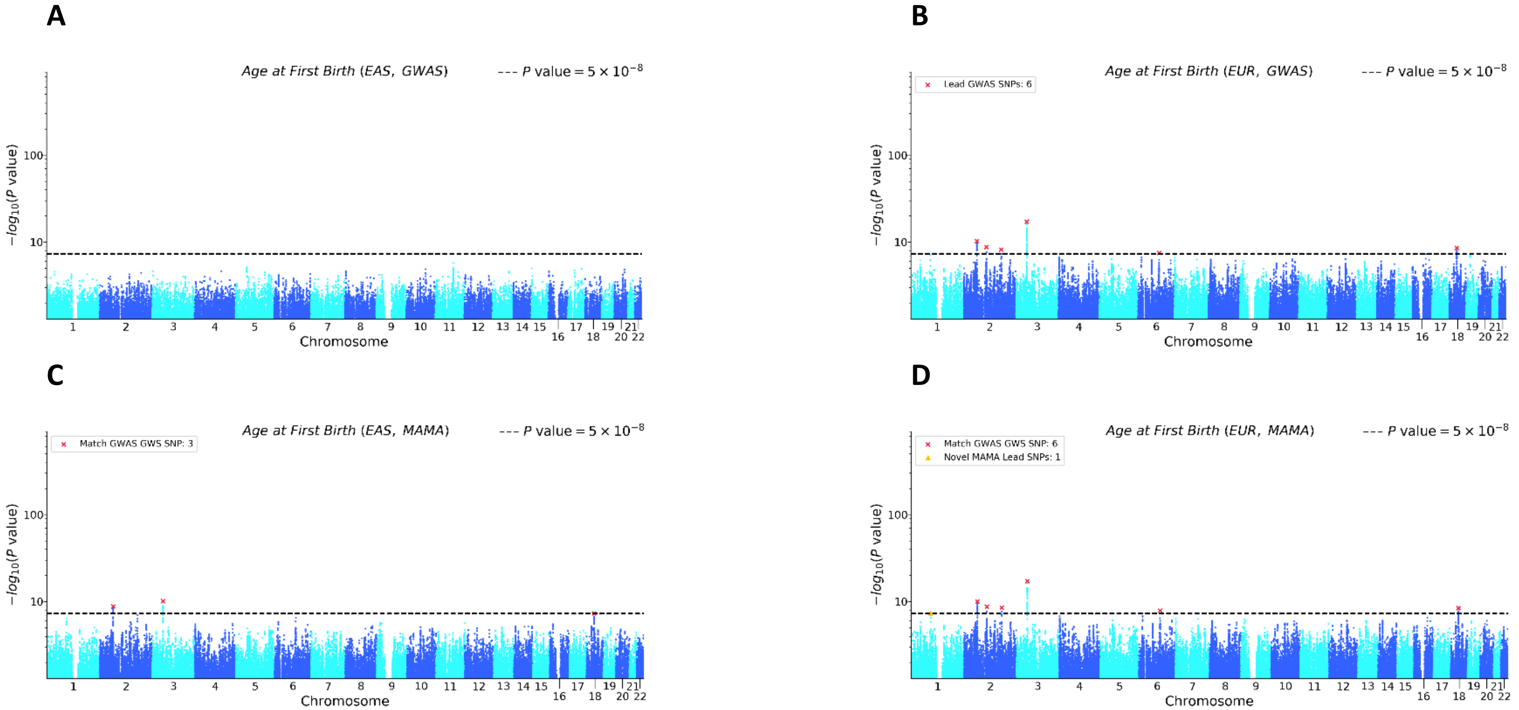


Note: (a) EAS GWAS, (b) EUR GWAS, (c) EAS MAMA, and (d) EUR MAMA. The x-axis is chromosomal position and the y-axis is the *P* value on a -log$_{10}$ scale (note the y-axes scale logarithmically). The dashed line marks the threshold for genome-wide significance ($P = 5 \times 10^{-8}$). For the GWAS Manhattan plots, lead SNPs (i.e., approximately independent SNPs surpassing the significance threshold) are marked with a red ×. For the MAMA Manhattan plots, lead SNPs are binned into one of three mutually exclusive categories: matching a GWAS genome-wide significant (GWS) SNP in either population (marked with a red ×), in LD with a GWAS gnome-wide significant (GWS) SNP in either population (marked with a black •), or a novel SNP that is independent of any GWAS lead SNP in either population (marked with a yellow ▲). Details on lead SNP and novel SNP identification can be found in Online Methods.

Figure S8. GWAS and MAMA Manhattan Plots for Basophil Count

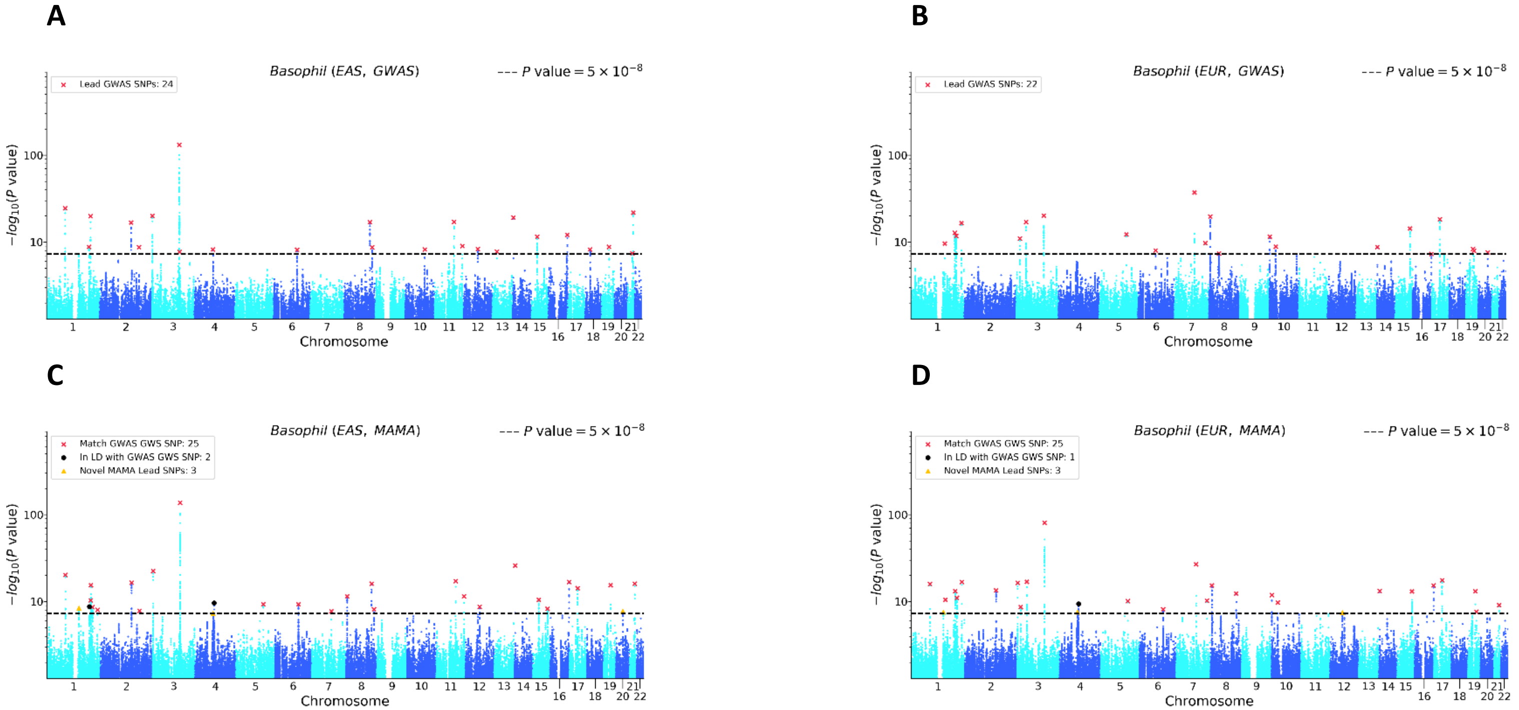


Note: (a) EAS GWAS, (b) EUR GWAS, (c) EAS MAMA, and (d) EUR MAMA. The x-axis is chromosomal position and the y-axis is the *P* value on a -$\log_{10}$ scale (note the y-axes scale logarithmically). The dashed line marks the threshold for genome-wide significance ($P = 5 \times 10^{-8}$). For the GWAS Manhattan plots, lead SNPs (i.e., approximately independent SNPs surpassing the significance threshold) are marked with a red ×. For the MAMA Manhattan plots, lead SNPs are binned into one of three mutually exclusive categories: matching a GWAS genome-wide significant (GWS) SNP in either population (marked with a red ×), in LD with a GWAS gnome-wide significant (GWS) SNP in either population (marked with a black •), or a novel SNP that is independent of any GWAS lead SNP in either population (marked with a yellow ▲). Details on lead SNP and novel SNP identification can be found in Online Methods.

Figure S9. GWAS and MAMA Manhattan Plots for Blood Pressure (Diastolic)

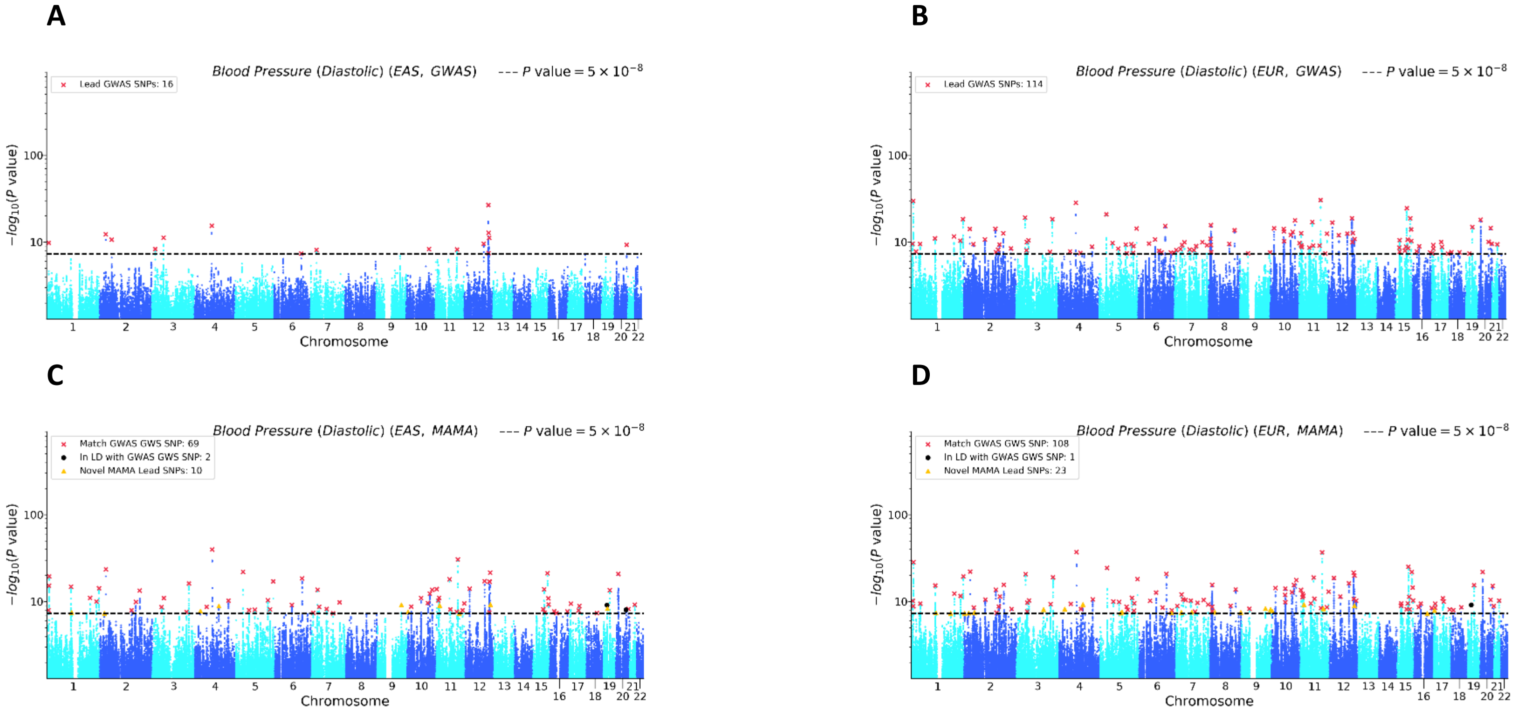


Note: (a) EAS GWAS, (b) EUR GWAS, (c) EAS MAMA, and (d) EUR MAMA. The x-axis is chromosomal position and the y-axis is the *P* value on a -$\log_{10}$ scale (note the y-axes scale logarithmically). The dashed line marks the threshold for genome-wide significance ($P = 5 \times 10^{-8}$). For the GWAS Manhattan plots, lead SNPs (i.e., approximately independent SNPs surpassing the significance threshold) are marked with a red ×. For the MAMA Manhattan plots, lead SNPs are binned into one of three mutually exclusive categories: matching a GWAS genome-wide significant (GWS) SNP in either population (marked with a red ×), in LD with a GWAS gnome-wide significant (GWS) SNP in either population (marked with a black •), or a novel SNP that is independent of any GWAS lead SNP in either population (marked with a yellow ▲). Details on lead SNP and novel SNP identification can be found in Online Methods.

Figure S10. GWAS and MAMA Manhattan Plots for Blood Pressure (Systolic)

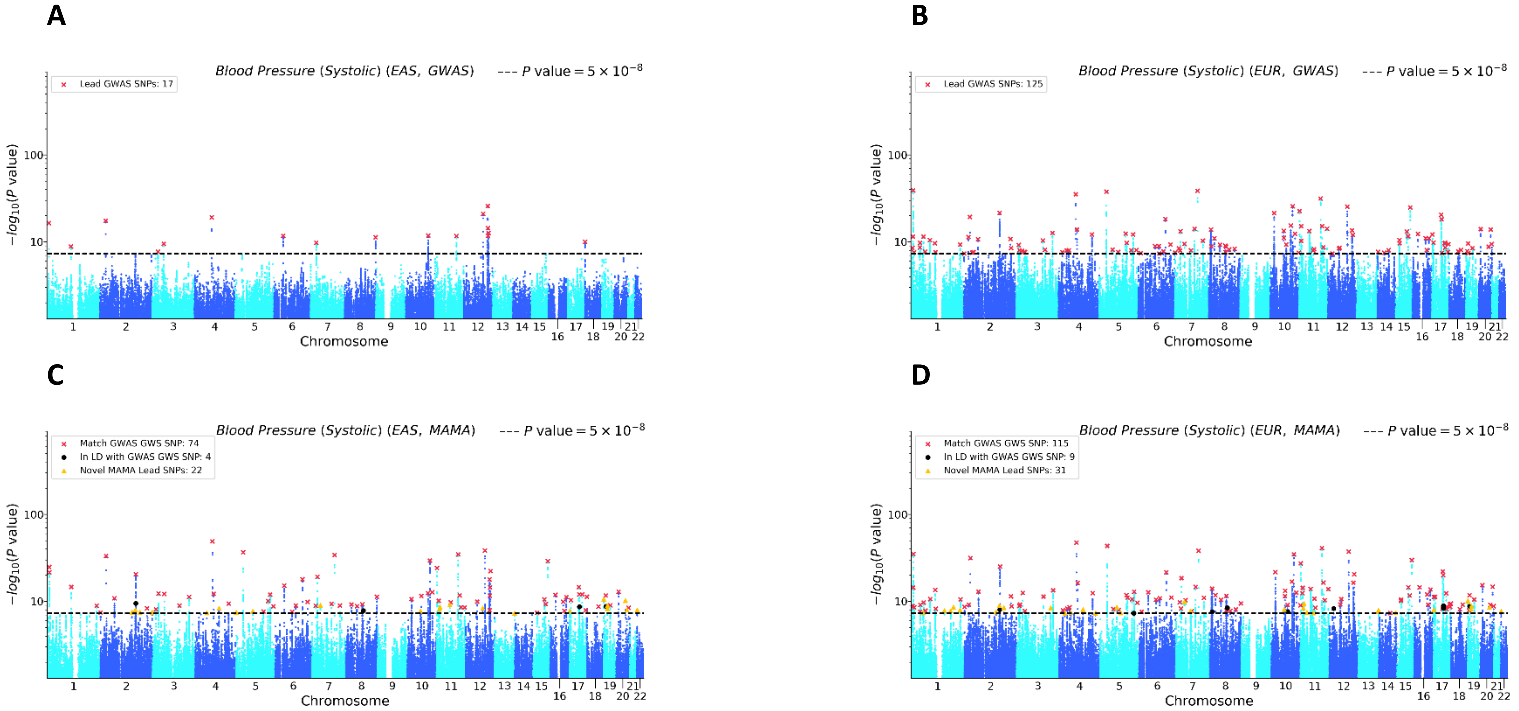


Note: (a) EAS GWAS, (b) EUR GWAS, (c) EAS MAMA, and (d) EUR MAMA. The x-axis is chromosomal position and the y-axis is the *P* value on a -$\log_{10}$ scale (note the y-axes scale logarithmically). The dashed line marks the threshold for genome-wide significance ($P = 5 \times 10^{-8}$). For the GWAS Manhattan plots, lead SNPs (i.e., approximately independent SNPs surpassing the significance threshold) are marked with a red ×. For the MAMA Manhattan plots, lead SNPs are binned into one of three mutually exclusive categories: matching a GWAS genome-wide significant (GWS) SNP in either population (marked with a red ×), in LD with a GWAS gnome-wide significant (GWS) SNP in either population (marked with a black •), or a novel SNP that is independent of any GWAS lead SNP in either population (marked with a yellow ▲). Details on lead SNP and novel SNP identification can be found in Online Methods.

Figure S11. GWAS and MAMA Manhattan Plots for BMI

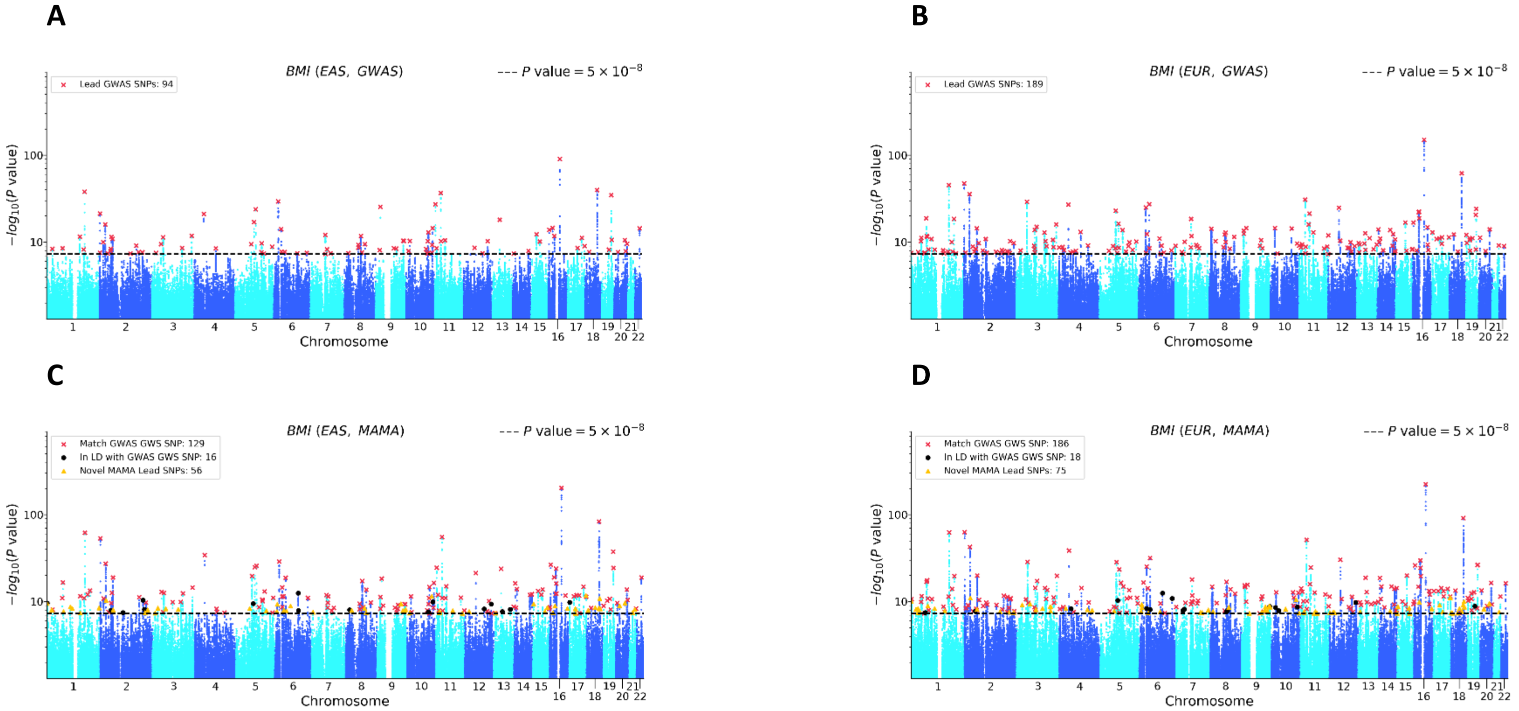


Note: (a) EAS GWAS, (b) EUR GWAS, (c) EAS MAMA, and (d) EUR MAMA. The x-axis is chromosomal position and the y-axis is the *P* value on a $-\log_{10}$ scale (note the y-axes scale logarithmically). The dashed line marks the threshold for genome-wide significance ($P = 5 \times 10^{-8}$). For the GWAS Manhattan plots, lead SNPs (i.e., approximately independent SNPs surpassing the significance threshold) are marked with a red ×. For the MAMA Manhattan plots, lead SNPs are binned into one of three mutually exclusive categories: matching a GWAS genome-wide significant (GWS) SNP in either population (marked with a red ×), in LD with a GWAS gnome-wide significant (GWS) SNP in either population (marked with a black •), or a novel SNP that is independent of any GWAS lead SNP in either population (marked with a yellow ▲). Details on lead SNP and novel SNP identification can be found in Online Methods.

Figure S12. GWAS and MAMA Manhattan Plots for Educational Attainment

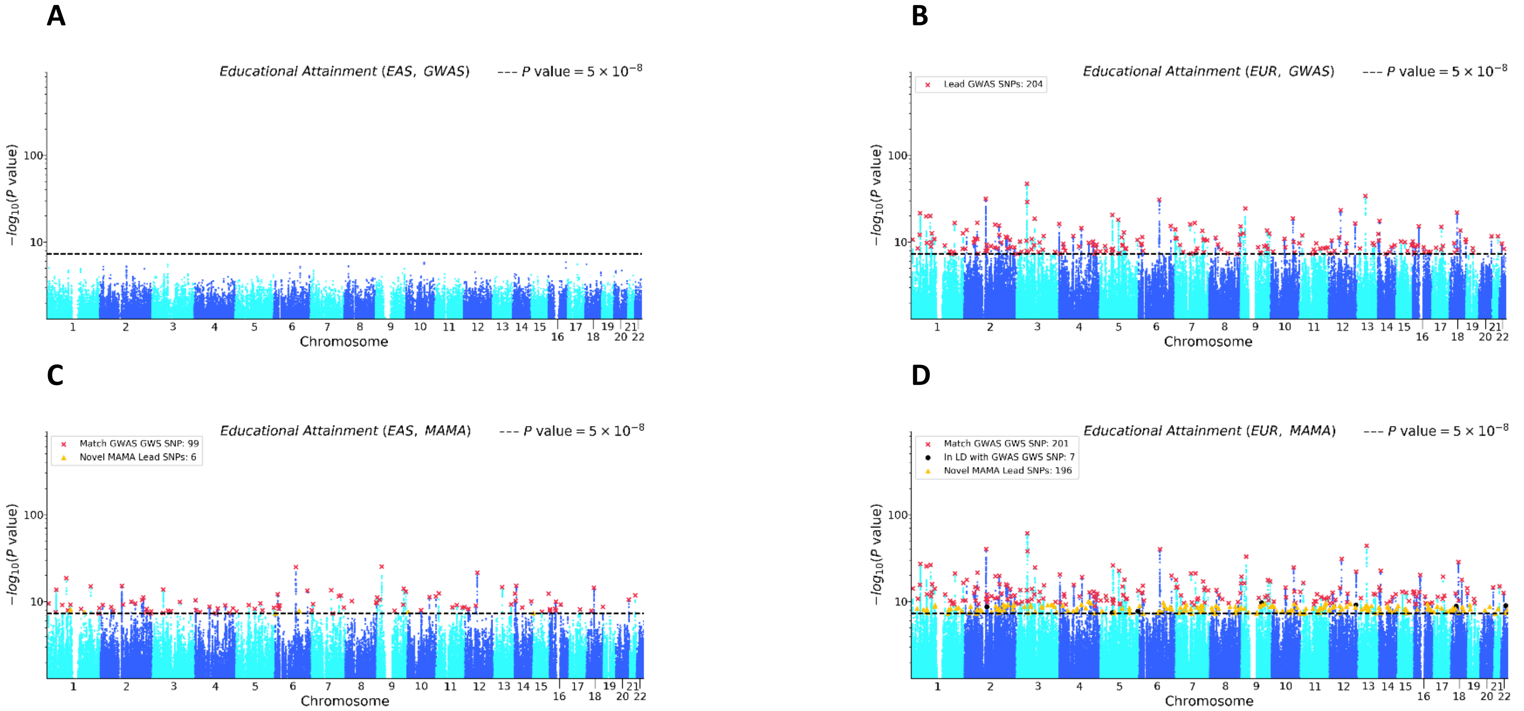


Note: (a) EAS GWAS, (b) EUR GWAS, (c) EAS MAMA, and (d) EUR MAMA. The x-axis is chromosomal position and the y-axis is the *P* value on a $-\log_{10}$ scale (note the y-axes scale logarithmically). The dashed line marks the threshold for genome-wide significance ($P = 5 \times 10^{-8}$). For the GWAS Manhattan plots, lead SNPs (i.e., approximately independent SNPs surpassing the significance threshold) are marked with a red ×. For the MAMA Manhattan plots, lead SNPs are binned into one of three mutually exclusive categories: matching a GWAS genome-wide significant (GWS) SNP in either population (marked with a red ×), in LD with a GWAS gnome-wide significant (GWS) SNP in either population (marked with a black •), or a novel SNP that is independent of any GWAS lead SNP in either population (marked with a yellow ▲). Details on lead SNP and novel SNP identification can be found in Online Methods.

Figure S13. GWAS and MAMA Manhattan Plots for Eosinophil Count

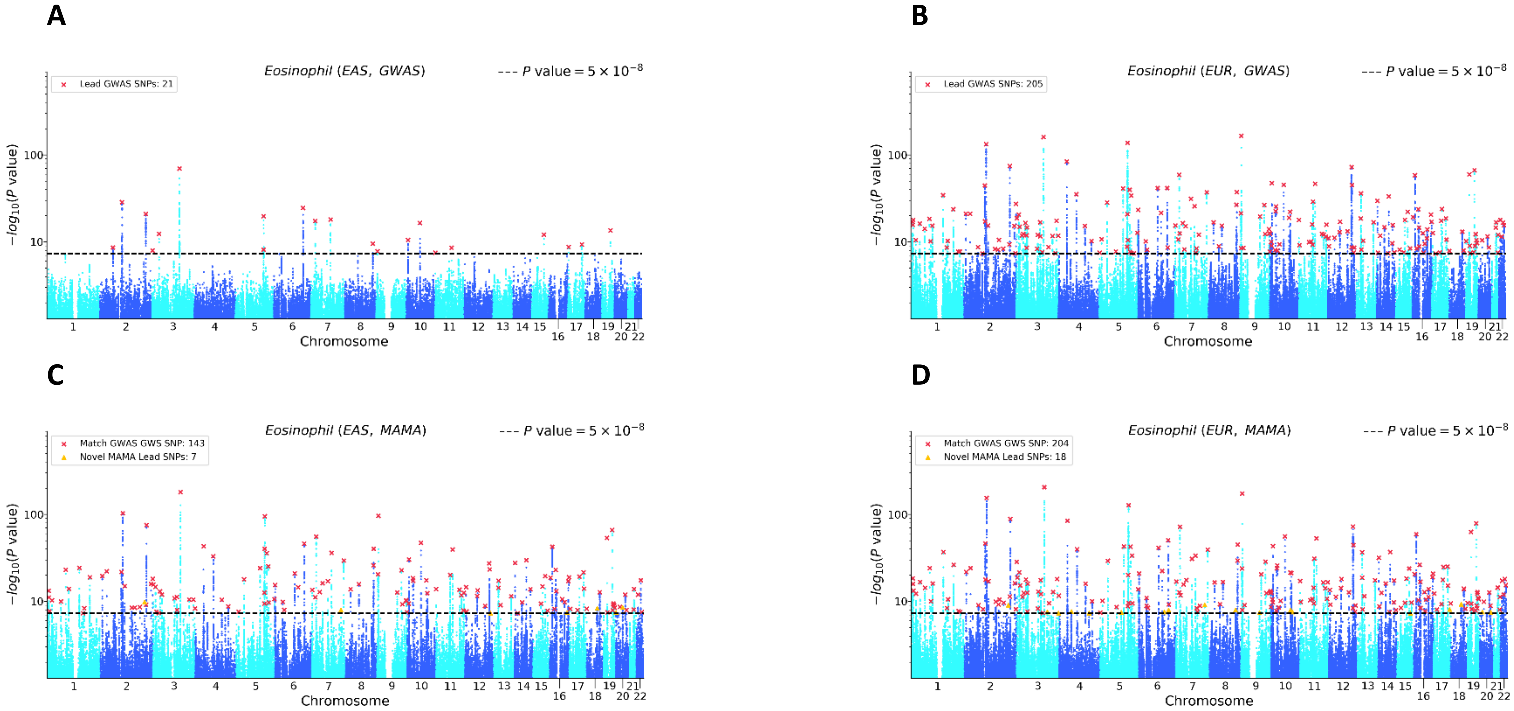


Note: (a) EAS GWAS, (b) EUR GWAS, (c) EAS MAMA, and (d) EUR MAMA. The x-axis is chromosomal position and the y-axis is the *P* value on a -$\log_{10}$ scale (note the y-axes scale logarithmically). The dashed line marks the threshold for genome-wide significance ($P = 5 \times 10^{-8}$). For the GWAS Manhattan plots, lead SNPs (i.e., approximately independent SNPs surpassing the significance threshold) are marked with a red ×. For the MAMA Manhattan plots, lead SNPs are binned into one of three mutually exclusive categories: matching a GWAS genome-wide significant (GWS) SNP in either population (marked with a red ×), in LD with a GWAS gnome-wide significant (GWS) SNP in either population (marked with a black •), or a novel SNP that is independent of any GWAS lead SNP in either population (marked with a yellow ▲). Details on lead SNP and novel SNP identification can be found in Online Methods.

Figure S14. GWAS and MAMA Manhattan Plots for Height

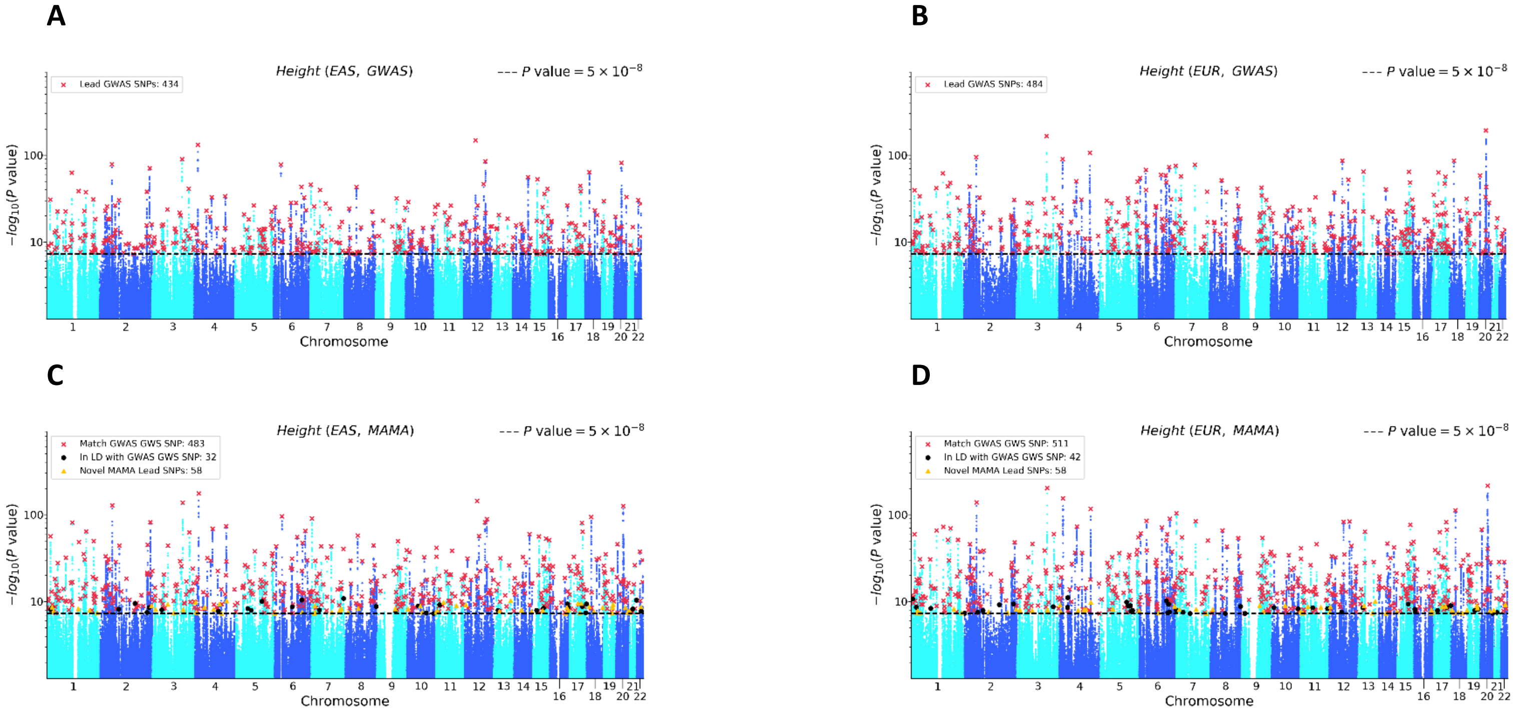


Note: (a) EAS GWAS, (b) EUR GWAS, (c) EAS MAMA, and (d) EUR MAMA. The x-axis is chromosomal position and the y-axis is the *P* value on a -$\log_{10}$ scale (note the y-axes scale logarithmically). The dashed line marks the threshold for genome-wide significance ($P = 5 \times 10^{-8}$). For the GWAS Manhattan plots, lead SNPs (i.e., approximately independent SNPs surpassing the significance threshold) are marked with a red ×. For the MAMA Manhattan plots, lead SNPs are binned into one of three mutually exclusive categories: matching a GWAS genome-wide significant (GWS) SNP in either population (marked with a red ×), in LD with a GWAS gnome-wide significant (GWS) SNP in either population (marked with a black •), or a novel SNP that is independent of any GWAS lead SNP in either population (marked with a yellow ▲). Details on lead SNP and novel SNP identification can be found in Online Methods.

Figure S15. GWAS and MAMA Manhattan Plots for Hematocrit

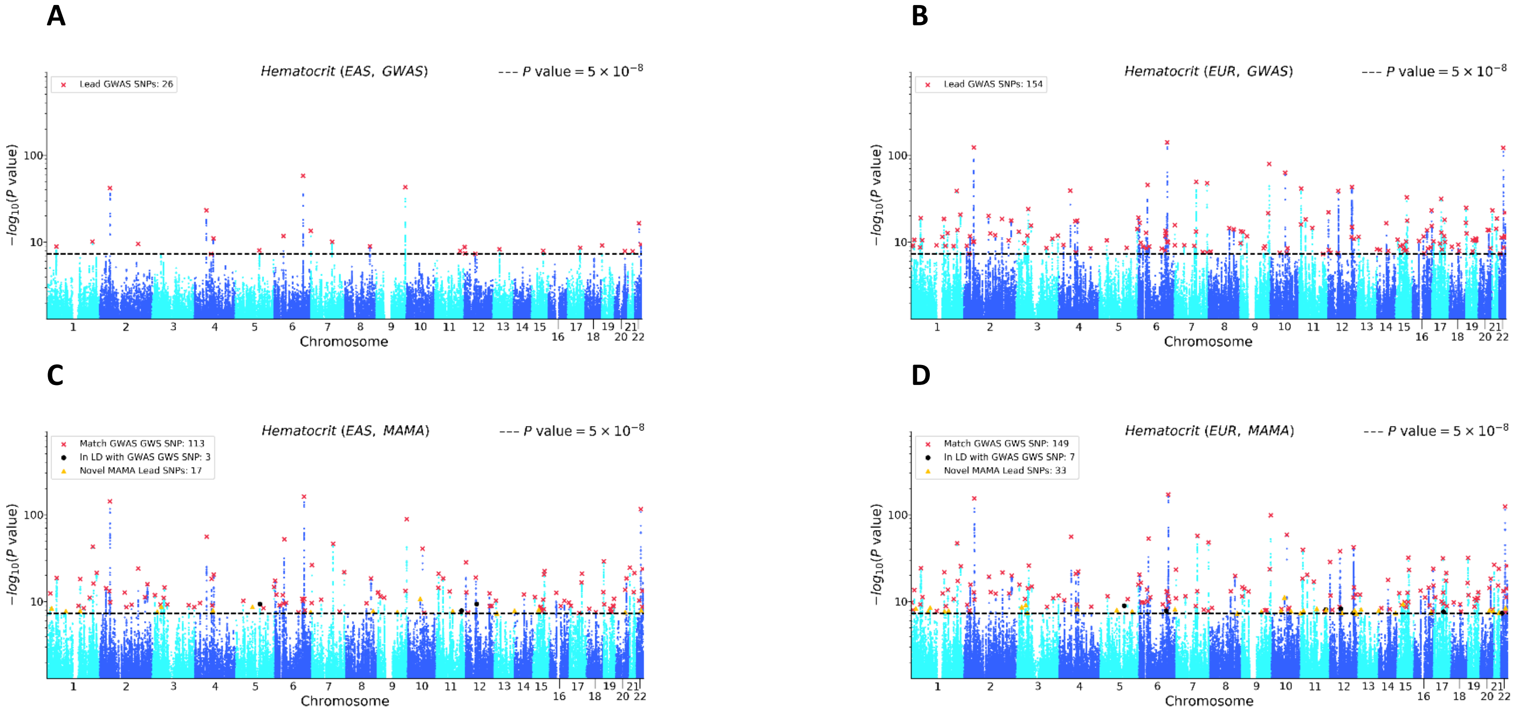


Note: (a) EAS GWAS, (b) EUR GWAS, (c) EAS MAMA, and (d) EUR MAMA. The x-axis is chromosomal position and the y-axis is the *P* value on a -$\log_{10}$ scale (note the y-axes scale logarithmically). The dashed line marks the threshold for genome-wide significance ($P = 5 \times 10^{-8}$). For the GWAS Manhattan plots, lead SNPs (i.e., approximately independent SNPs surpassing the significance threshold) are marked with a red ×. For the MAMA Manhattan plots, lead SNPs are binned into one of three mutually exclusive categories: matching a GWAS genome-wide significant (GWS) SNP in either population (marked with a red ×), in LD with a GWAS gnome-wide significant (GWS) SNP in either population (marked with a black •), or a novel SNP that is independent of any GWAS lead SNP in either population (marked with a yellow ▲). Details on lead SNP and novel SNP identification can be found in Online Methods.

Figure S16. GWAS and MAMA Manhattan Plots for Hemoglobin

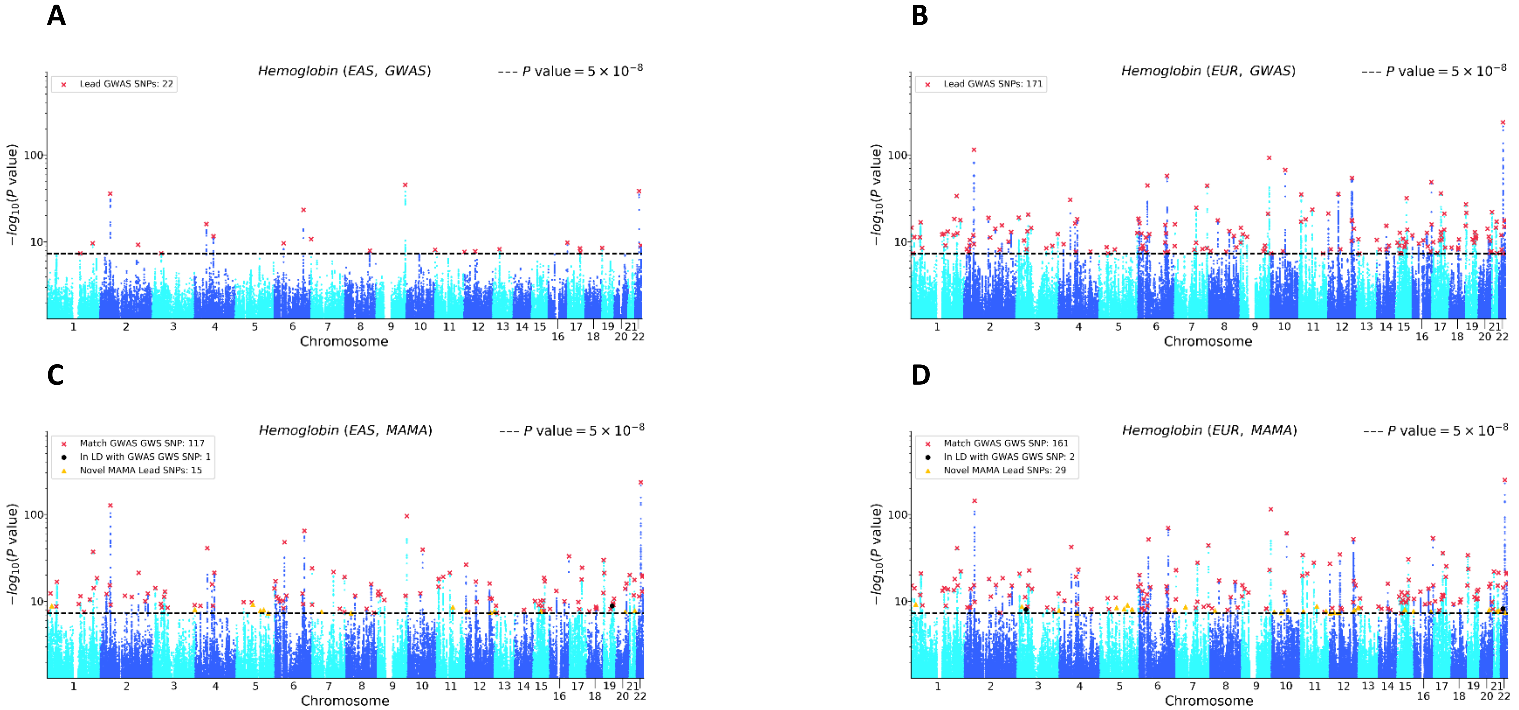


Note: (a) EAS GWAS, (b) EUR GWAS, (c) EAS MAMA, and (d) EUR MAMA. The x-axis is chromosomal position and the y-axis is the *P* value on a $-\log_{10}$ scale (note the y-axes scale logarithmically). The dashed line marks the threshold for genome-wide significance ($P = 5 \times 10^{-8}$). For the GWAS Manhattan plots, lead SNPs (i.e., approximately independent SNPs surpassing the significance threshold) are marked with a red ×. For the MAMA Manhattan plots, lead SNPs are binned into one of three mutually exclusive categories: matching a GWAS genome-wide significant (GWS) SNP in either population (marked with a red ×), in LD with a GWAS gnome-wide significant (GWS) SNP in either population (marked with a black •), or a novel SNP that is independent of any GWAS lead SNP in either population (marked with a yellow ▲). Details on lead SNP and novel SNP identification can be found in Online Methods.

Figure S17. GWAS and MAMA Manhattan Plots for Lymphocyte Count

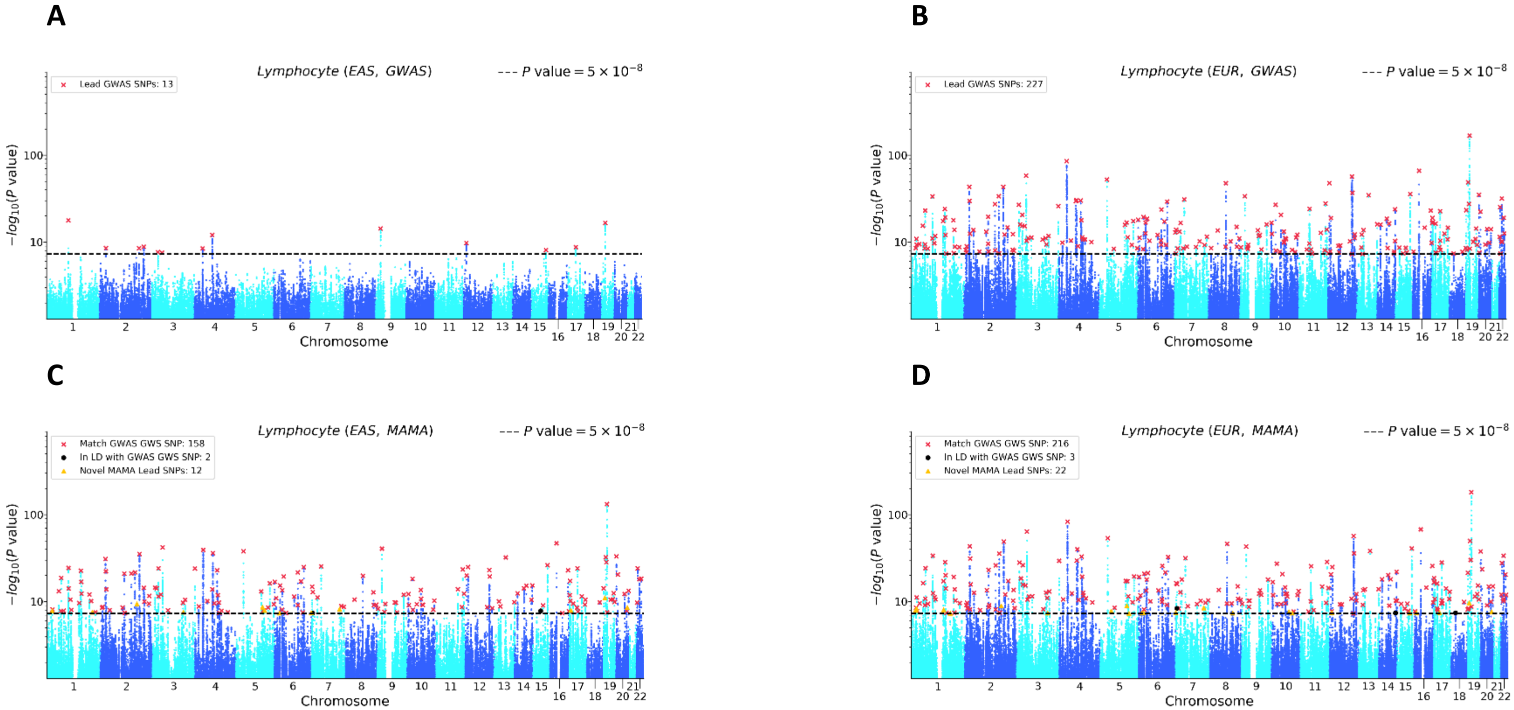


Note: (a) EAS GWAS, (b) EUR GWAS, (c) EAS MAMA, and (d) EUR MAMA. The x-axis is chromosomal position and the y-axis is the *P* value on a $-\log_{10}$ scale (note the y-axes scale logarithmically). The dashed line marks the threshold for genome-wide significance ($P = 5 \times 10^{-8}$). For the GWAS Manhattan plots, lead SNPs (i.e., approximately independent SNPs surpassing the significance threshold) are marked with a red ×. For the MAMA Manhattan plots, lead SNPs are binned into one of three mutually exclusive categories: matching a GWAS genome-wide significant (GWS) SNP in either population (marked with a red ×), in LD with a GWAS gnome-wide significant (GWS) SNP in either population (marked with a black •), or a novel SNP that is independent of any GWAS lead SNP in either population (marked with a yellow ▲). Details on lead SNP and novel SNP identification can be found in Online Methods.

Figure S18. GWAS and MAMA Manhattan Plots for Mean Corpuscular Hemoglobin

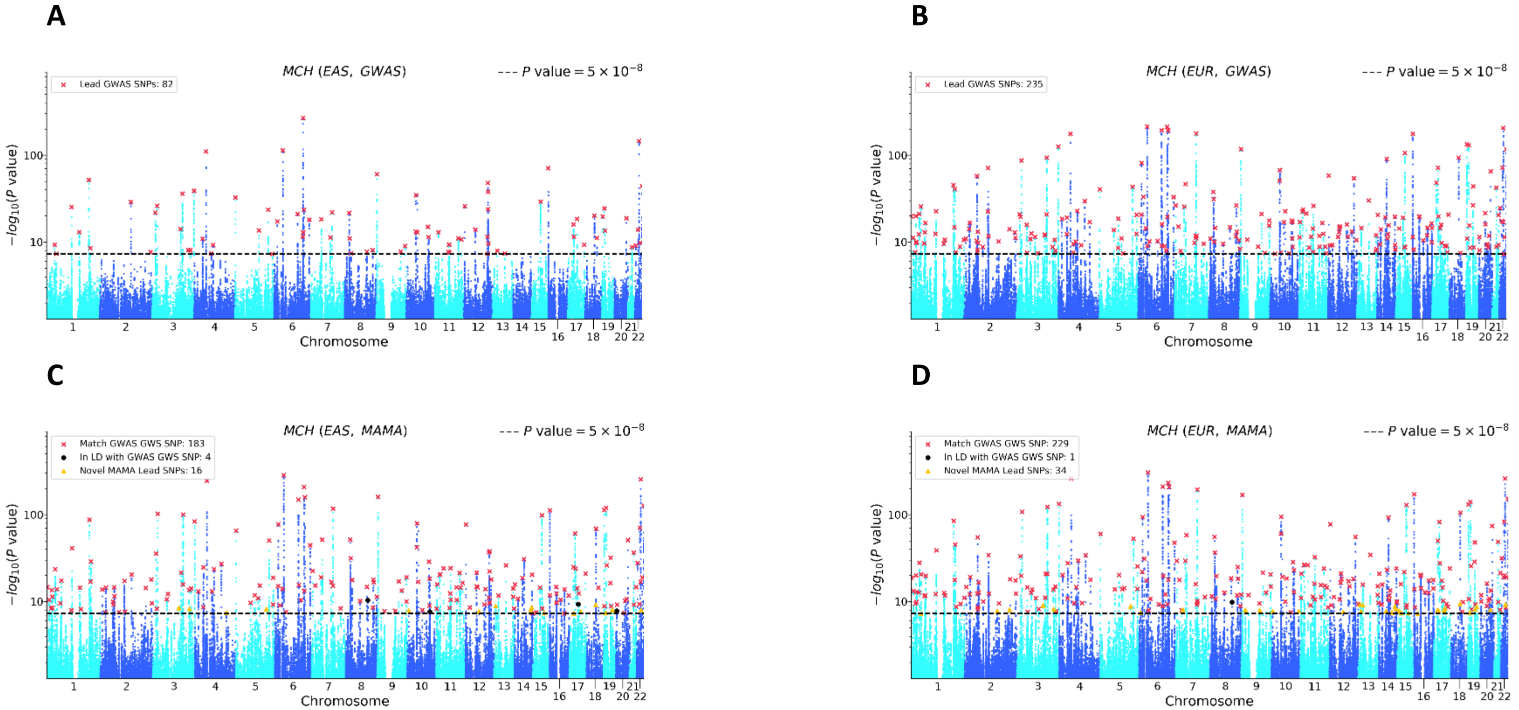


Note: (a) EAS GWAS, (b) EUR GWAS, (c) EAS MAMA, and (d) EUR MAMA. The x-axis is chromosomal position and the y-axis is the *P* value on a $-\log_{10}$ scale (note the y-axes scale logarithmically). The dashed line marks the threshold for genome-wide significance ($P = 5 \times 10^{-8}$). For the GWAS Manhattan plots, lead SNPs (i.e., approximately independent SNPs surpassing the significance threshold) are marked with a red ×. For the MAMA Manhattan plots, lead SNPs are binned into one of three mutually exclusive categories: matching a GWAS genome-wide significant (GWS) SNP in either population (marked with a red ×), in LD with a GWAS gnome-wide significant (GWS) SNP in either population (marked with a black •), or a novel SNP that is independent of any GWAS lead SNP in either population (marked with a yellow ▲). Details on lead SNP and novel SNP identification can be found in Online Methods.

Figure S19. GWAS and MAMA Manhattan Plots for Mean Corpuscular Hemoglobin Concentration

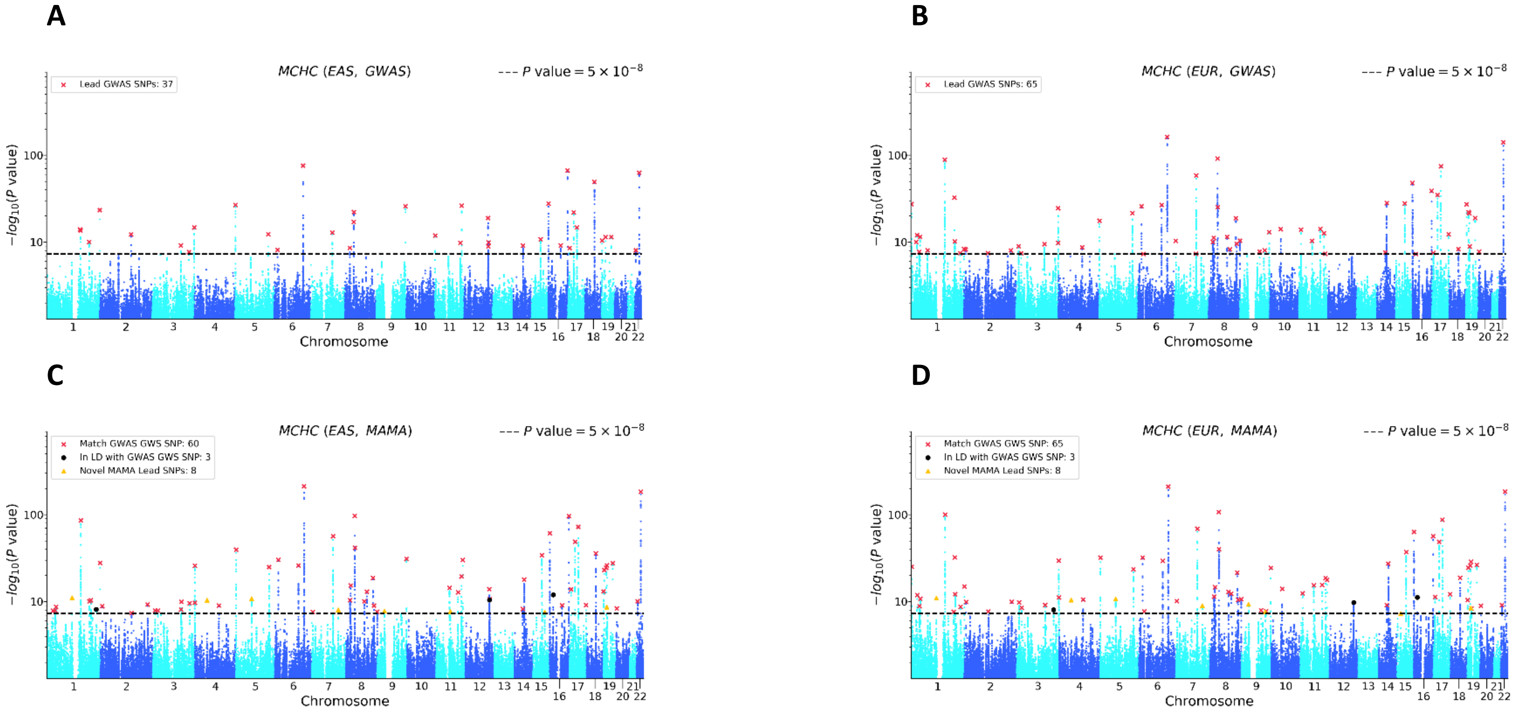


Note: (a) EAS GWAS, (b) EUR GWAS, (c) EAS MAMA, and (d) EUR MAMA. The x-axis is chromosomal position and the y-axis is the *P* value on a $-\log_{10}$ scale (note the y-axes scale logarithmically). The dashed line marks the threshold for genome-wide significance ($P = 5 \times 10^{-8}$). For the GWAS Manhattan plots, lead SNPs (i.e., approximately independent SNPs surpassing the significance threshold) are marked with a red ×. For the MAMA Manhattan plots, lead SNPs are binned into one of three mutually exclusive categories: matching a GWAS genome-wide significant (GWS) SNP in either population (marked with a red ×), in LD with a GWAS gnome-wide significant (GWS) SNP in either population (marked with a black •), or a novel SNP that is independent of any GWAS lead SNP in either population (marked with a yellow ▲). Details on lead SNP and novel SNP identification can be found in Online Methods.

Figure S20. GWAS and MAMA Manhattan Plots for Mean Corpuscular Volume

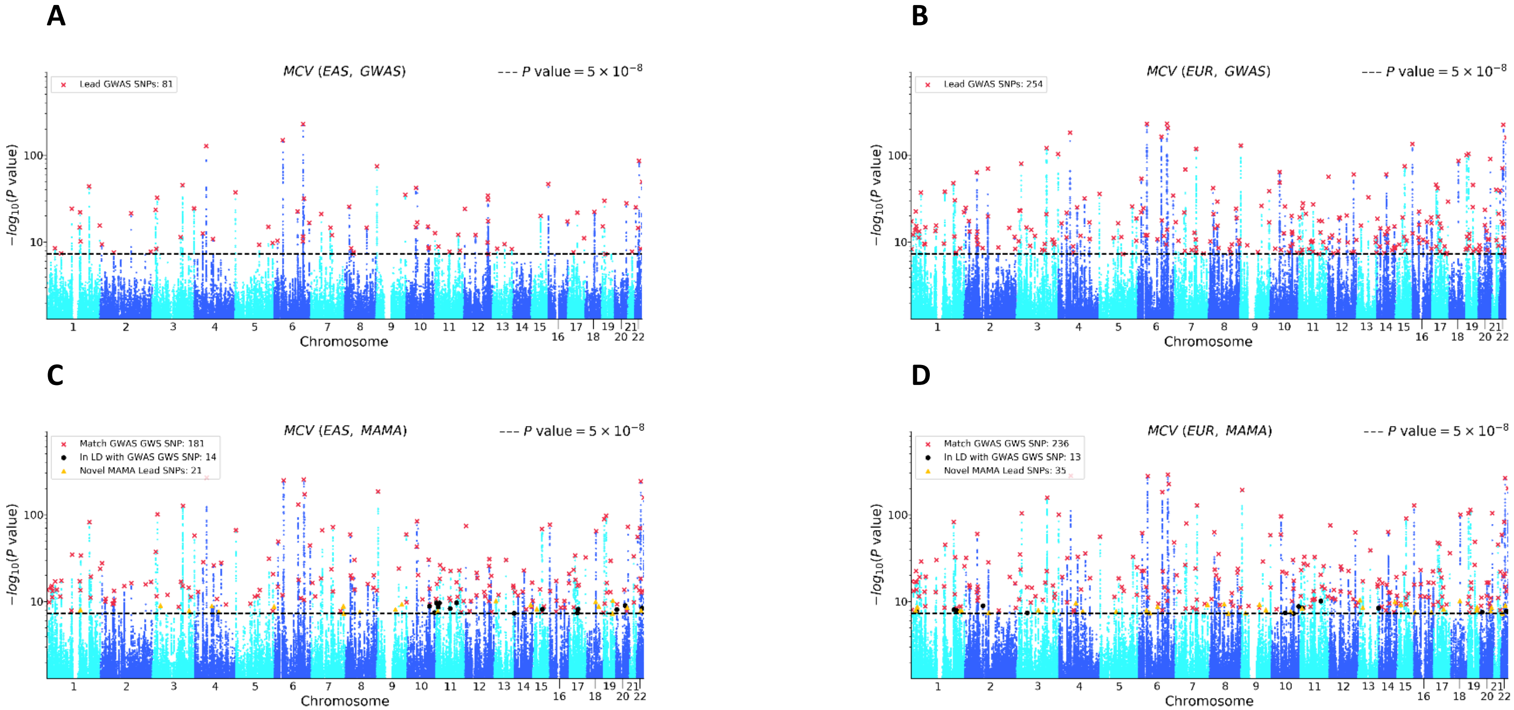


Note: (a) EAS GWAS, (b) EUR GWAS, (c) EAS MAMA, and (d) EUR MAMA. The x-axis is chromosomal position and the y-axis is the *P* value on a $-\log_{10}$ scale (note the y-axes scale logarithmically). The dashed line marks the threshold for genome-wide significance ($P = 5 \times 10^{-8}$). For the GWAS Manhattan plots, lead SNPs (i.e., approximately independent SNPs surpassing the significance threshold) are marked with a red ×. For the MAMA Manhattan plots, lead SNPs are binned into one of three mutually exclusive categories: matching a GWAS genome-wide significant (GWS) SNP in either population (marked with a red ×), in LD with a GWAS gnome-wide significant (GWS) SNP in either population (marked with a black •), or a novel SNP that is independent of any GWAS lead SNP in either population (marked with a yellow ▲). Details on lead SNP and novel SNP identification can be found in Online Methods.

Figure S21. GWAS and MAMA Manhattan Plots for Monocyte Count

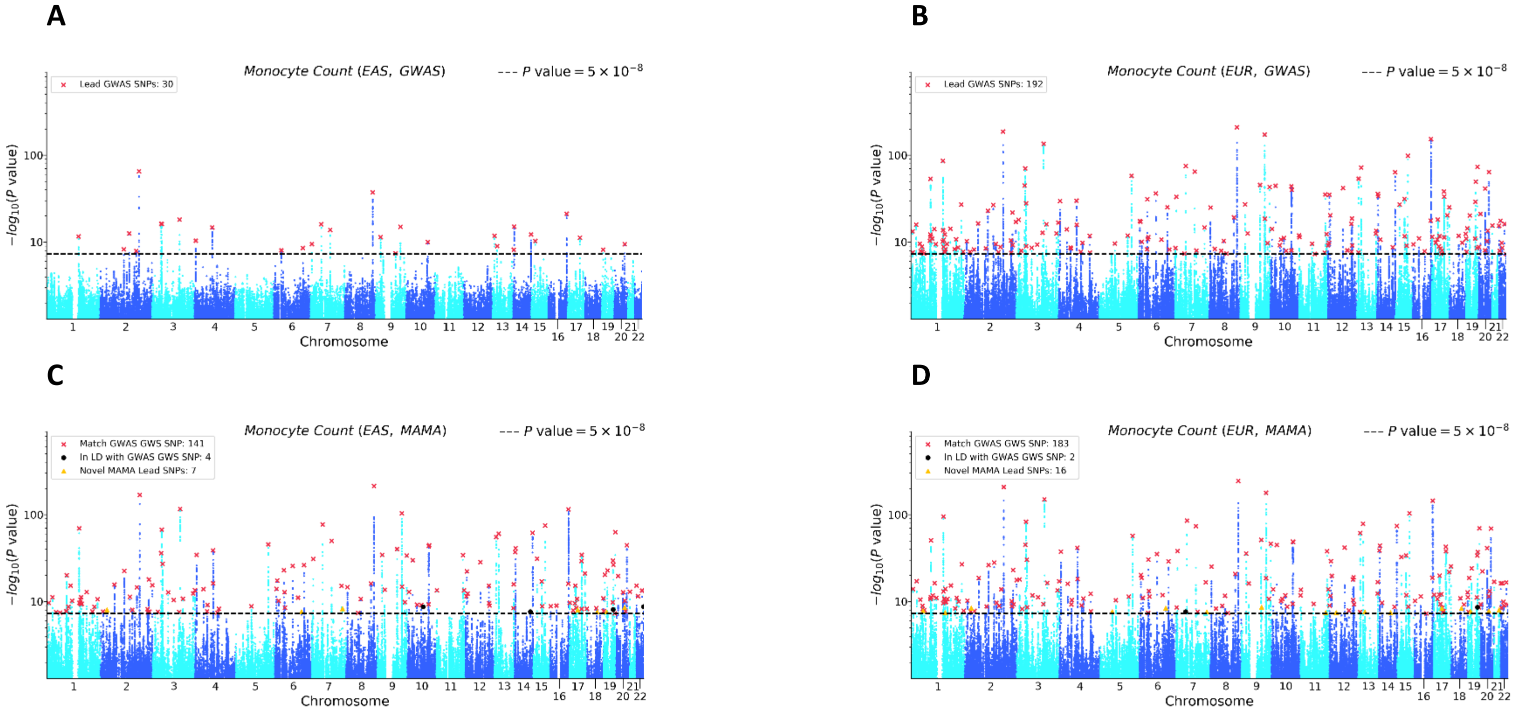


Note: (a) EAS GWAS, (b) EUR GWAS, (c) EAS MAMA, and (d) EUR MAMA. The x-axis is chromosomal position and the y-axis is the *P* value on a -$\log_{10}$ scale (note the y-axes scale logarithmically). The dashed line marks the threshold for genome-wide significance ($P = 5 \times 10^{-8}$). For the GWAS Manhattan plots, lead SNPs (i.e., approximately independent SNPs surpassing the significance threshold) are marked with a red ×. For the MAMA Manhattan plots, lead SNPs are binned into one of three mutually exclusive categories: matching a GWAS genome-wide significant (GWS) SNP in either population (marked with a red ×), in LD with a GWAS gnome-wide significant (GWS) SNP in either population (marked with a black •), or a novel SNP that is independent of any GWAS lead SNP in either population (marked with a yellow ▲). Details on lead SNP and novel SNP identification can be found in Online Methods.

Figure S22. GWAS and MAMA Manhattan Plots for Neutrophil Count

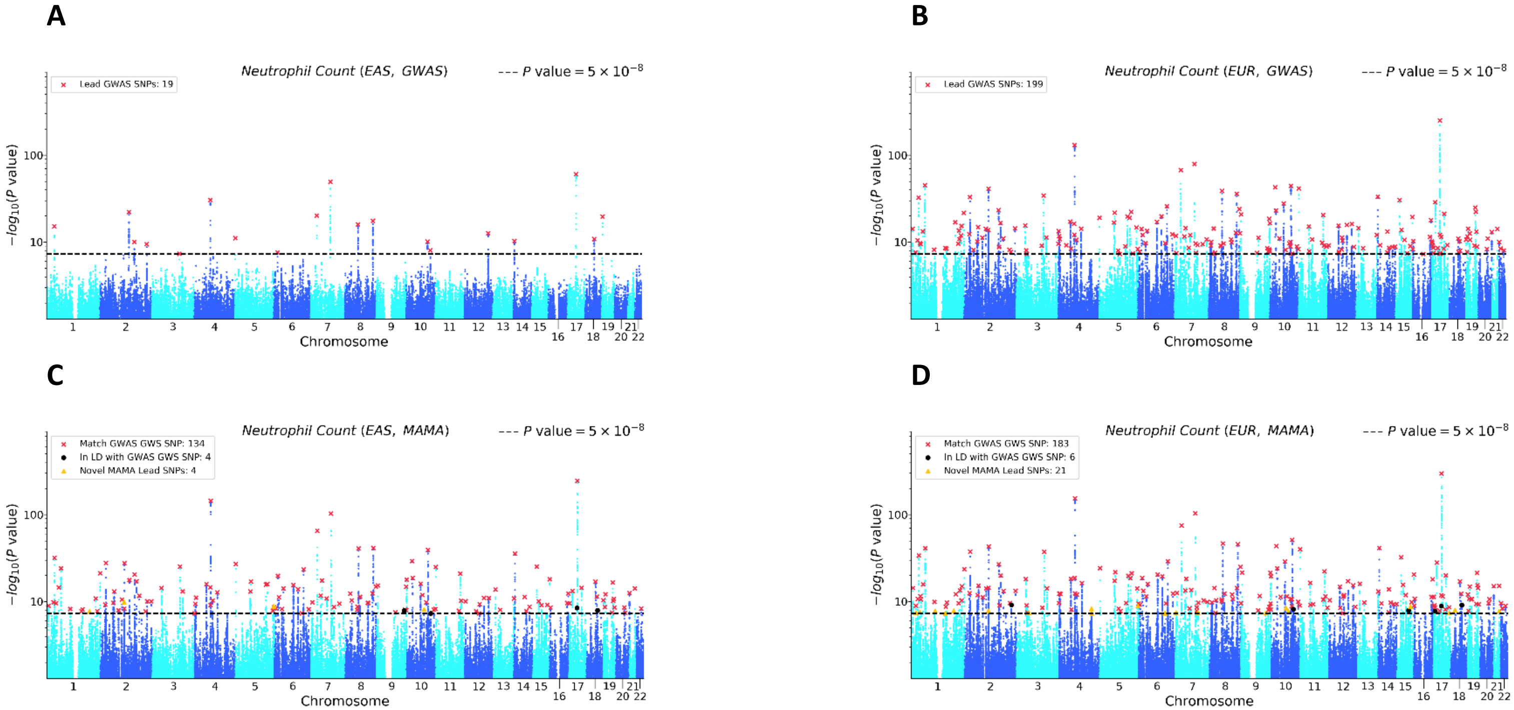


Note: (a) EAS GWAS, (b) EUR GWAS, (c) EAS MAMA, and (d) EUR MAMA. The x-axis is chromosomal position and the y-axis is the *P* value on a $-\log_{10}$ scale (note the y-axes scale logarithmically). The dashed line marks the threshold for genome-wide significance ($P = 5 \times 10^{-8}$). For the GWAS Manhattan plots, lead SNPs (i.e., approximately independent SNPs surpassing the significance threshold) are marked with a red ×. For the MAMA Manhattan plots, lead SNPs are binned into one of three mutually exclusive categories: matching a GWAS genome-wide significant (GWS) SNP in either population (marked with a red ×), in LD with a GWAS gnome-wide significant (GWS) SNP in either population (marked with a black •), or a novel SNP that is independent of any GWAS lead SNP in either population (marked with a yellow ▲). Details on lead SNP and novel SNP identification can be found in Online Methods.

Figure S23. GWAS and MAMA Manhattan Plots for Physical Activity

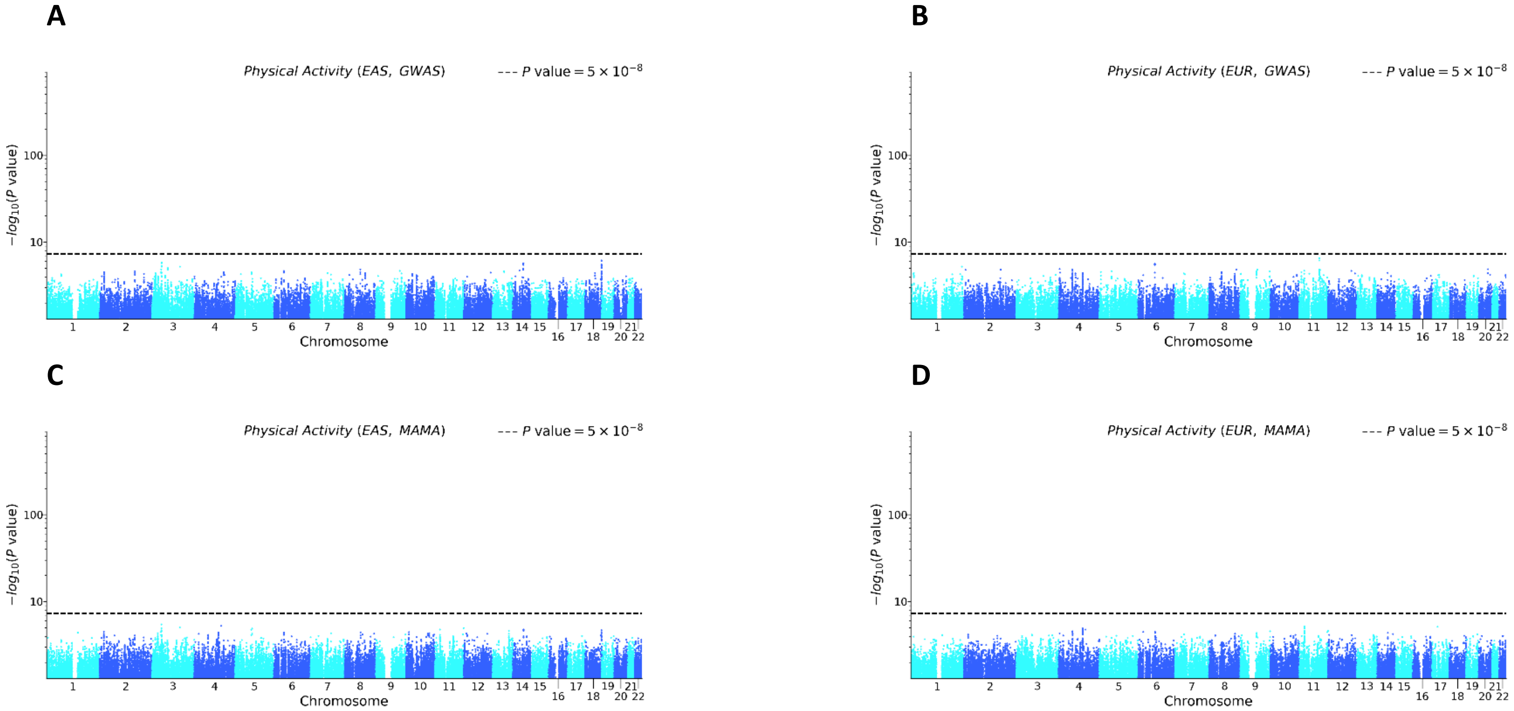


Note: (a) EAS GWAS, (b) EUR GWAS, (c) EAS MAMA, and (d) EUR MAMA. The x-axis is chromosomal position and the y-axis is the *P* value on a -$\log_{10}$ scale (note the y-axes scale logarithmically). The dashed line marks the threshold for genome-wide significance ($P = 5 \times 10^{-8}$). For the GWAS Manhattan plots, lead SNPs (i.e., approximately independent SNPs surpassing the significance threshold) are marked with a red ×. For the MAMA Manhattan plots, lead SNPs are binned into one of three mutually exclusive categories: matching a GWAS genome-wide significant (GWS) SNP in either population (marked with a red ×), in LD with a GWAS gnome-wide significant (GWS) SNP in either population (marked with a black •), or a novel SNP that is independent of any GWAS lead SNP in either population (marked with a yellow ▲). Details on lead SNP and novel SNP identification can be found in Online Methods.

Figure S24. GWAS and MAMA Manhattan Plots for Platelet Count

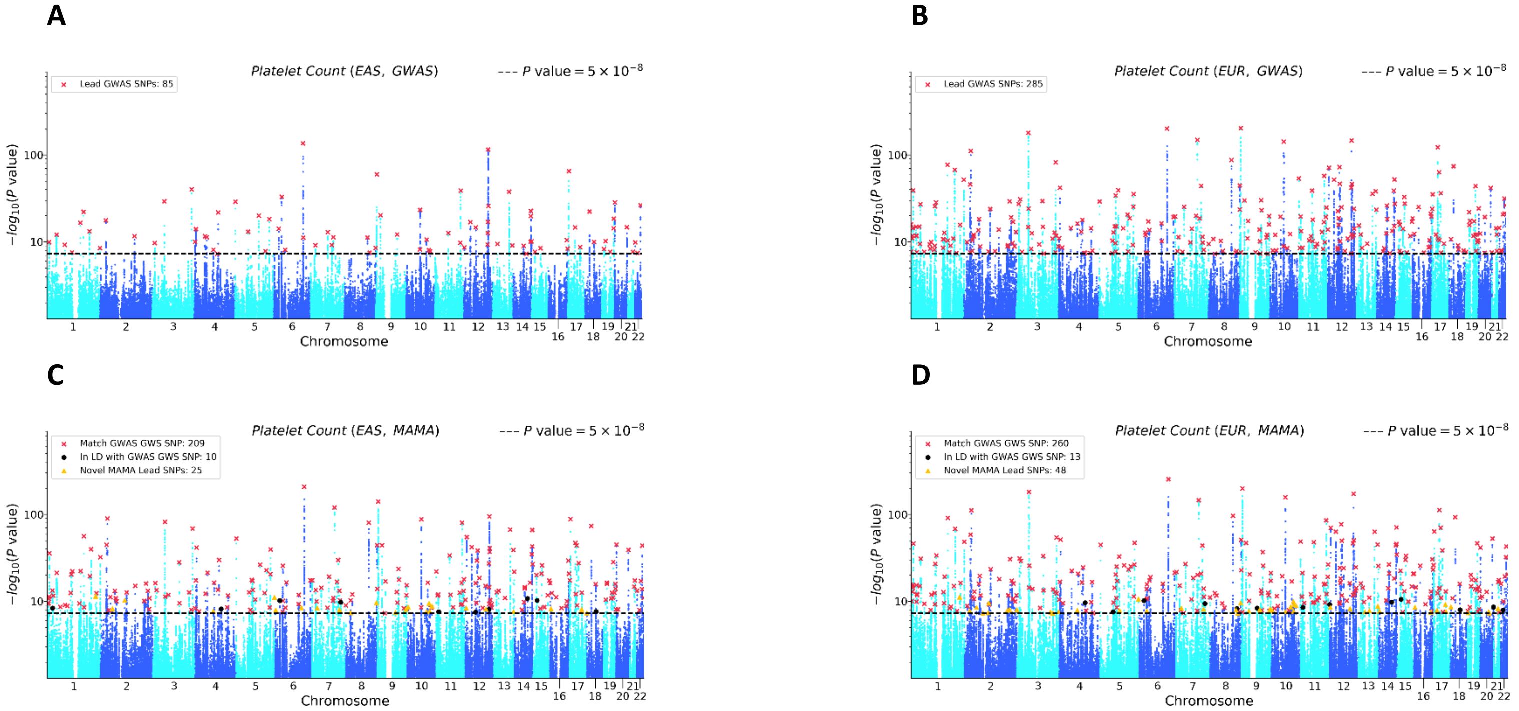


Note: (a) EAS GWAS, (b) EUR GWAS, (c) EAS MAMA, and (d) EUR MAMA. The x-axis is chromosomal position and the y-axis is the *P* value on a $-\log_{10}$ scale (note the y-axes scale logarithmically). The dashed line marks the threshold for genome-wide significance ($P = 5 \times 10^{-8}$). For the GWAS Manhattan plots, lead SNPs (i.e., approximately independent SNPs surpassing the significance threshold) are marked with a red ×. For the MAMA Manhattan plots, lead SNPs are binned into one of three mutually exclusive categories: matching a GWAS genome-wide significant (GWS) SNP in either population (marked with a red ×), in LD with a GWAS gnome-wide significant (GWS) SNP in either population (marked with a black •), or a novel SNP that is independent of any GWAS lead SNP in either population (marked with a yellow ▲). Details on lead SNP and novel SNP identification can be found in Online Methods.

Figure S25. GWAS and MAMA Manhattan Plots for Red Blood Cell Count

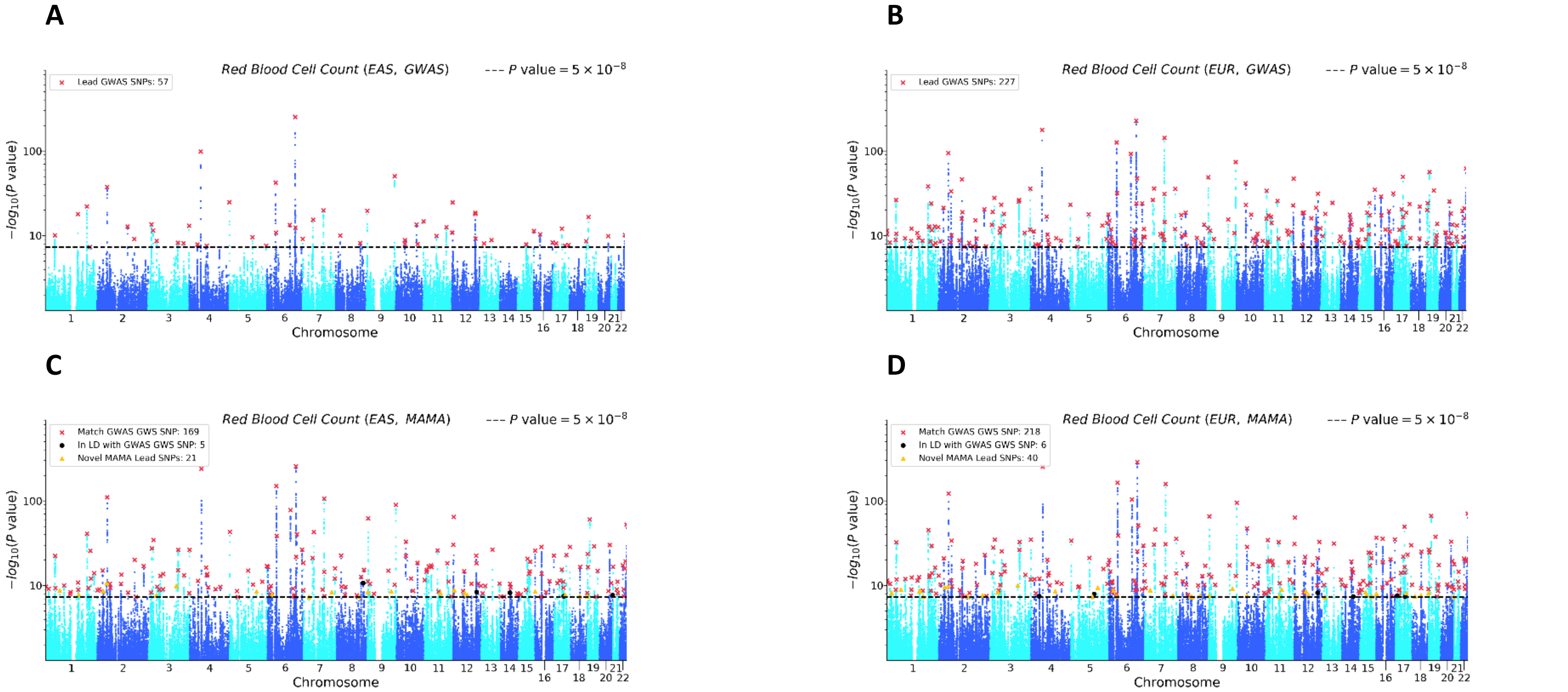


Note: (a) EAS GWAS, (b) EUR GWAS, (c) EAS MAMA, and (d) EUR MAMA. The x-axis is chromosomal position and the y-axis is the *P* value on a $-\log_{10}$ scale (note the y-axes scale logarithmically). The dashed line marks the threshold for genome-wide significance ($P = 5 \times 10^{-8}$). For the GWAS Manhattan plots, lead SNPs (i.e., approximately independent SNPs surpassing the significance threshold) are marked with a red ×. For the MAMA Manhattan plots, lead SNPs are binned into one of three mutually exclusive categories: matching a GWAS genome-wide significant (GWS) SNP in either population (marked with a red ×), in LD with a GWAS gnome-wide significant (GWS) SNP in either population (marked with a black •), or a novel SNP that is independent of any GWAS lead SNP in either population (marked with a yellow ▲). Details on lead SNP and novel SNP identification can be found in Online Methods.

Figure S26. GWAS and MAMA Manhattan Plots for Schizophrenia

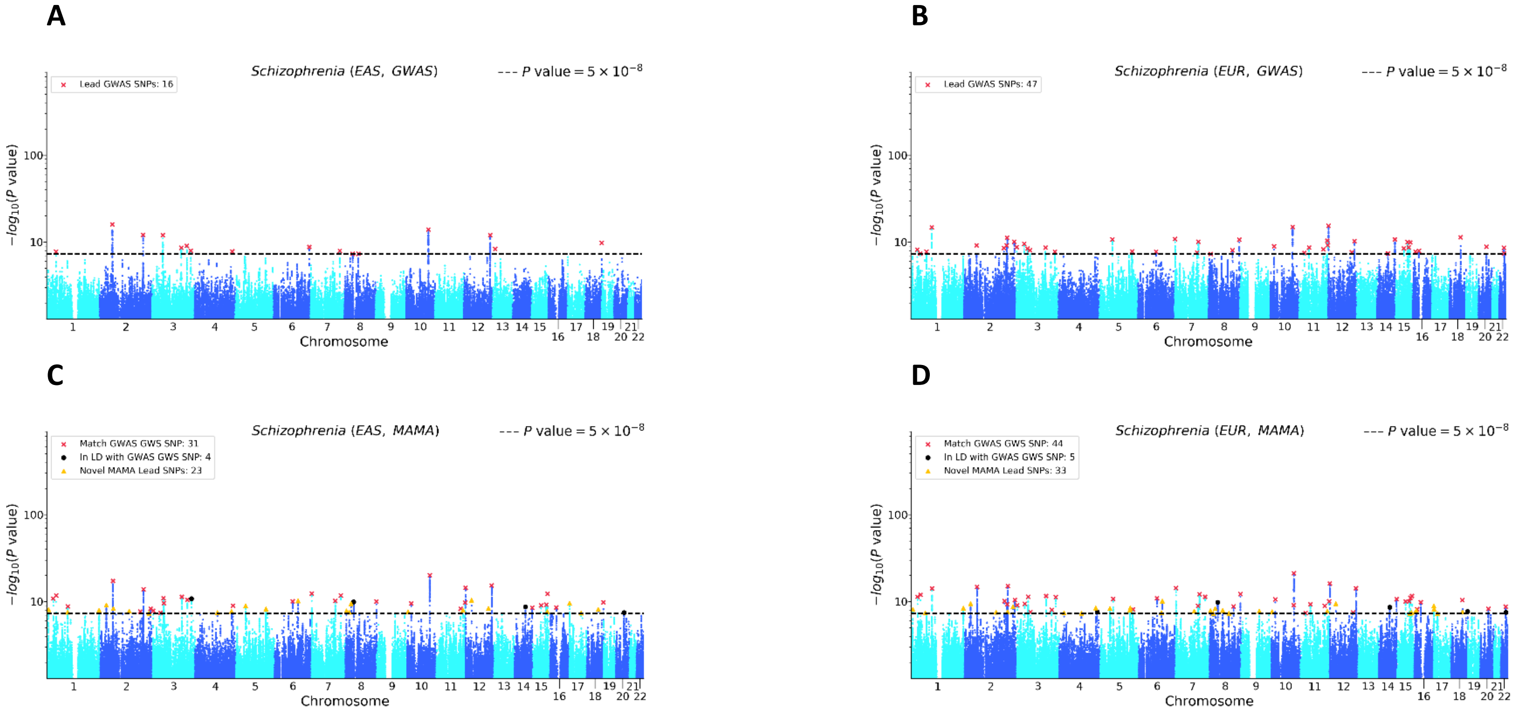


Note: (a) EAS GWAS, (b) EUR GWAS, (c) EAS MAMA, and (d) EUR MAMA. The x-axis is chromosomal position and the y-axis is the *P* value on a $-\log_{10}$ scale (note the y-axes scale logarithmically). The dashed line marks the threshold for genome-wide significance ($P = 5 \times 10^{-8}$). For the GWAS Manhattan plots, lead SNPs (i.e., approximately independent SNPs surpassing the significance threshold) are marked with a red ×. For the MAMA Manhattan plots, lead SNPs are binned into one of three mutually exclusive categories: matching a GWAS genome-wide significant (GWS) SNP in either population (marked with a red ×), in LD with a GWAS gnome-wide significant (GWS) SNP in either population (marked with a black •), or a novel SNP that is independent of any GWAS lead SNP in either population (marked with a yellow ▲). Details on lead SNP and novel SNP identification can be found in Online Methods.

Figure S27. GWAS and MAMA Manhattan Plots for Smoking Initiation

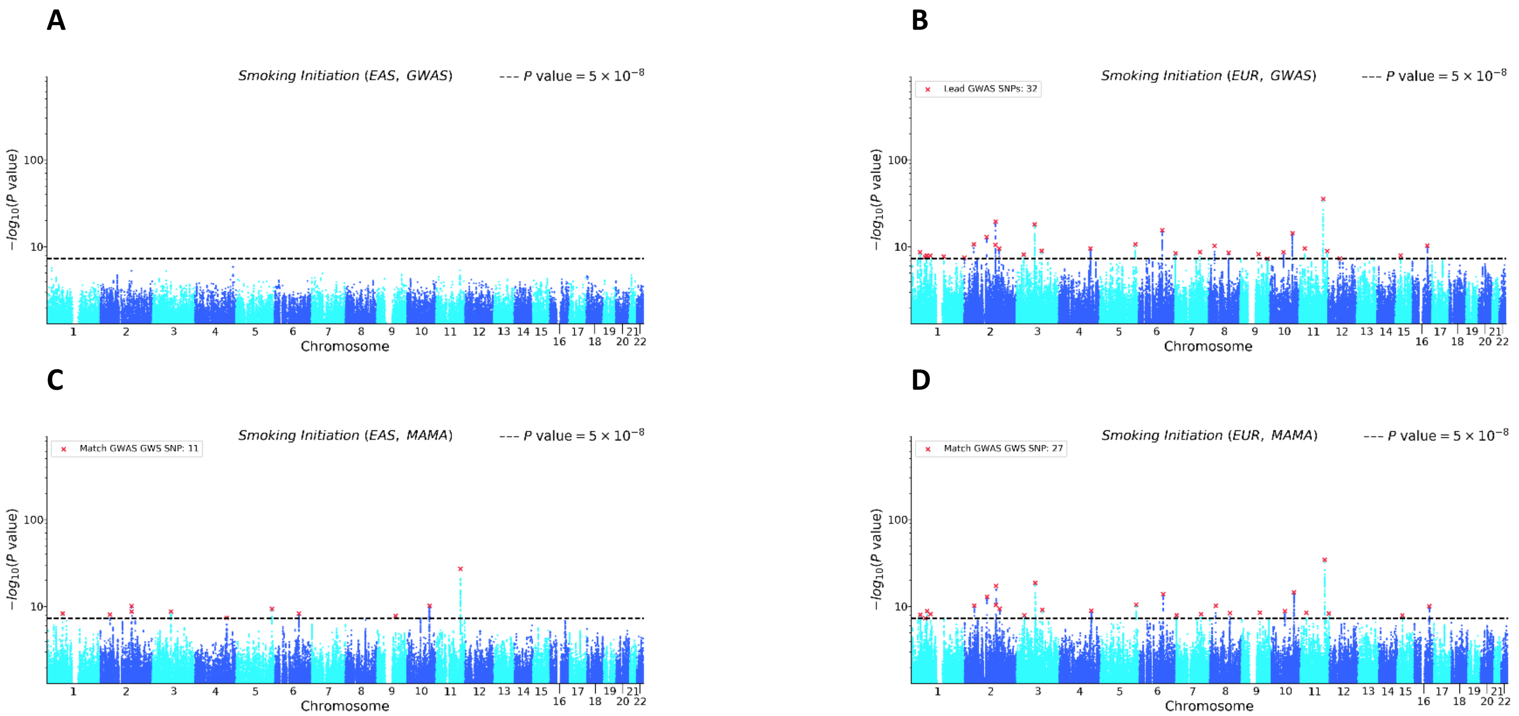


Note: (a) EAS GWAS, (b) EUR GWAS, (c) EAS MAMA, and (d) EUR MAMA. The x-axis is chromosomal position and the y-axis is the *P* value on a -$\log_{10}$ scale (note the y-axes scale logarithmically). The dashed line marks the threshold for genome-wide significance ($P = 5 \times 10^{-8}$). For the GWAS Manhattan plots, lead SNPs (i.e., approximately independent SNPs surpassing the significance threshold) are marked with a red ×. For the MAMA Manhattan plots, lead SNPs are binned into one of three mutually exclusive categories: matching a GWAS genome-wide significant (GWS) SNP in either population (marked with a red ×), in LD with a GWAS gnome-wide significant (GWS) SNP in either population (marked with a black •), or a novel SNP that is independent of any GWAS lead SNP in either population (marked with a yellow ▲). Details on lead SNP and novel SNP identification can be found in Online Methods.

Figure S28. GWAS and MAMA Manhattan Plots for Weight

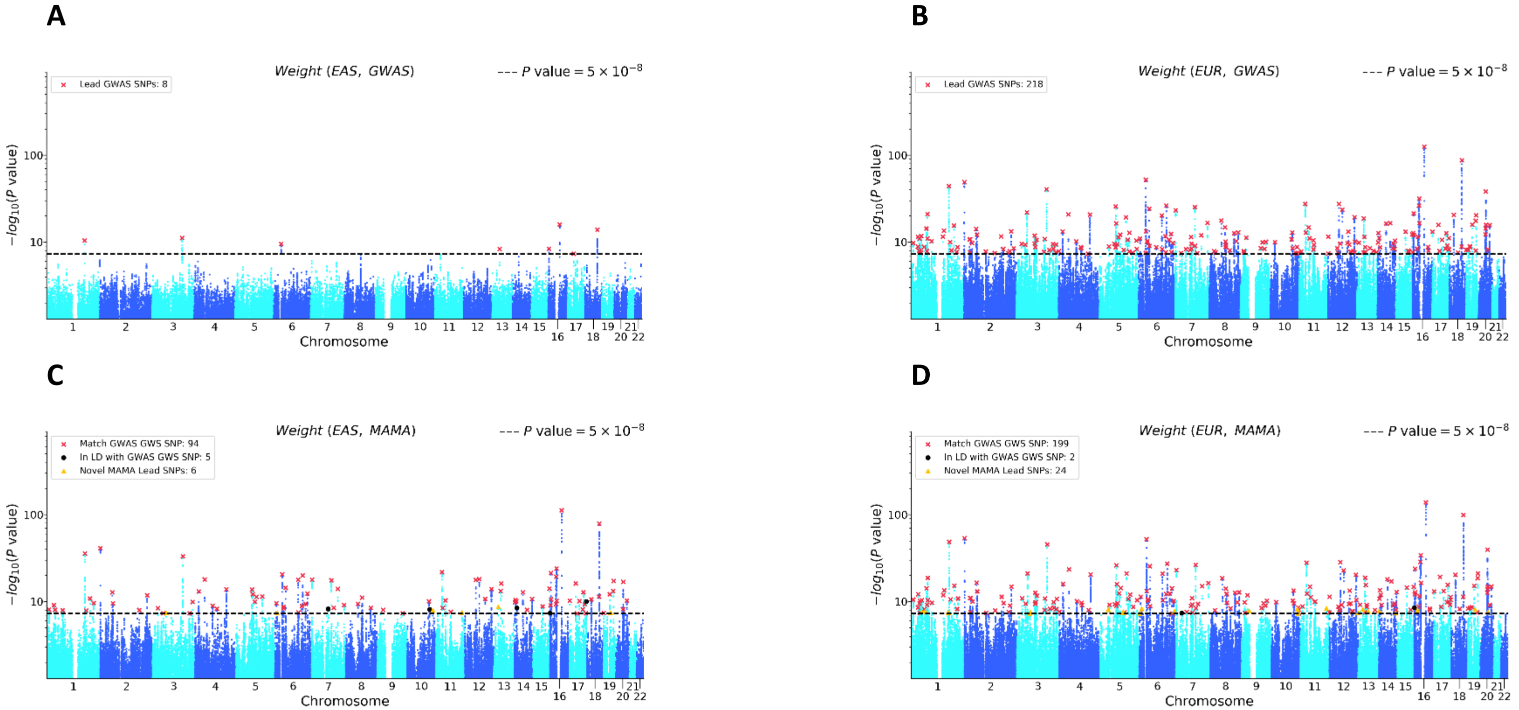


Note: (a) EAS GWAS, (b) EUR GWAS, (c) EAS MAMA, and (d) EUR MAMA. The x-axis is chromosomal position and the y-axis is the *P* value on a $-\log_{10}$ scale (note the y-axes scale logarithmically). The dashed line marks the threshold for genome-wide significance ($P = 5 \times 10^{-8}$). For the GWAS Manhattan plots, lead SNPs (i.e., approximately independent SNPs surpassing the significance threshold) are marked with a red ×. For the MAMA Manhattan plots, lead SNPs are binned into one of three mutually exclusive categories: matching a GWAS genome-wide significant (GWS) SNP in either population (marked with a red ×), in LD with a GWAS gnome-wide significant (GWS) SNP in either population (marked with a black •), or a novel SNP that is independent of any GWAS lead SNP in either population (marked with a yellow ▲). Details on lead SNP and novel SNP identification can be found in Online Methods.

Figure S29. GWAS and MAMA Manhattan Plots for White Blood Cell Count

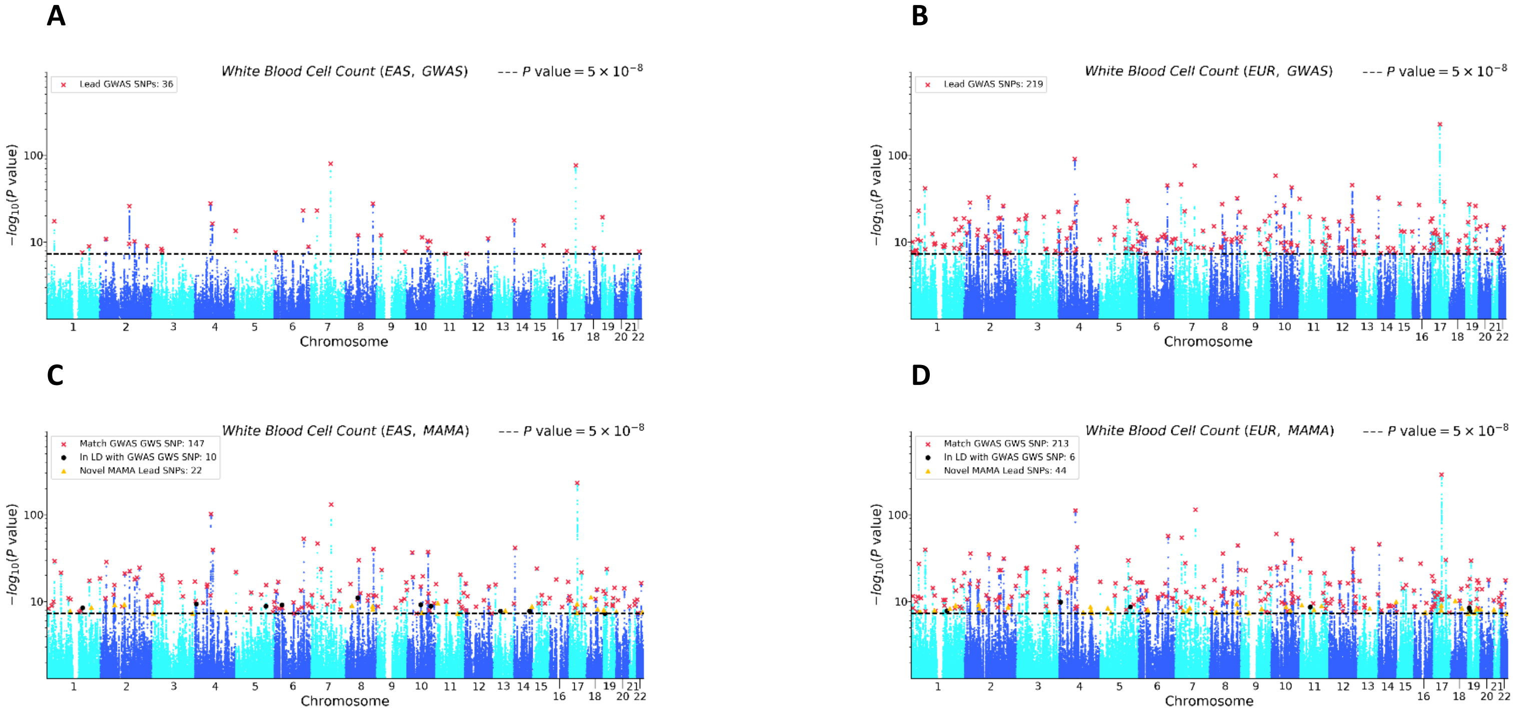


Note: (a) EAS GWAS, (b) EUR GWAS, (c) EAS MAMA, and (d) EUR MAMA. The x-axis is chromosomal position and the y-axis is the *P* value on a $-\log_{10}$ scale (note the y-axes scale logarithmically). The dashed line marks the threshold for genome-wide significance ($P = 5 \times 10^{-8}$). For the GWAS Manhattan plots, lead SNPs (i.e., approximately independent SNPs surpassing the significance threshold) are marked with a red ×. For the MAMA Manhattan plots, lead SNPs are binned into one of three mutually exclusive categories: matching a GWAS genome-wide significant (GWS) SNP in either population (marked with a red ×), in LD with a GWAS gnome-wide significant (GWS) SNP in either population (marked with a black •), or a novel SNP that is independent of any GWAS lead SNP in either population (marked with a yellow ▲). Details on lead SNP and novel SNP identification can be found in Online Methods.

Figure S30. Replication of MAMA Lead SNPs for Age at First Birth

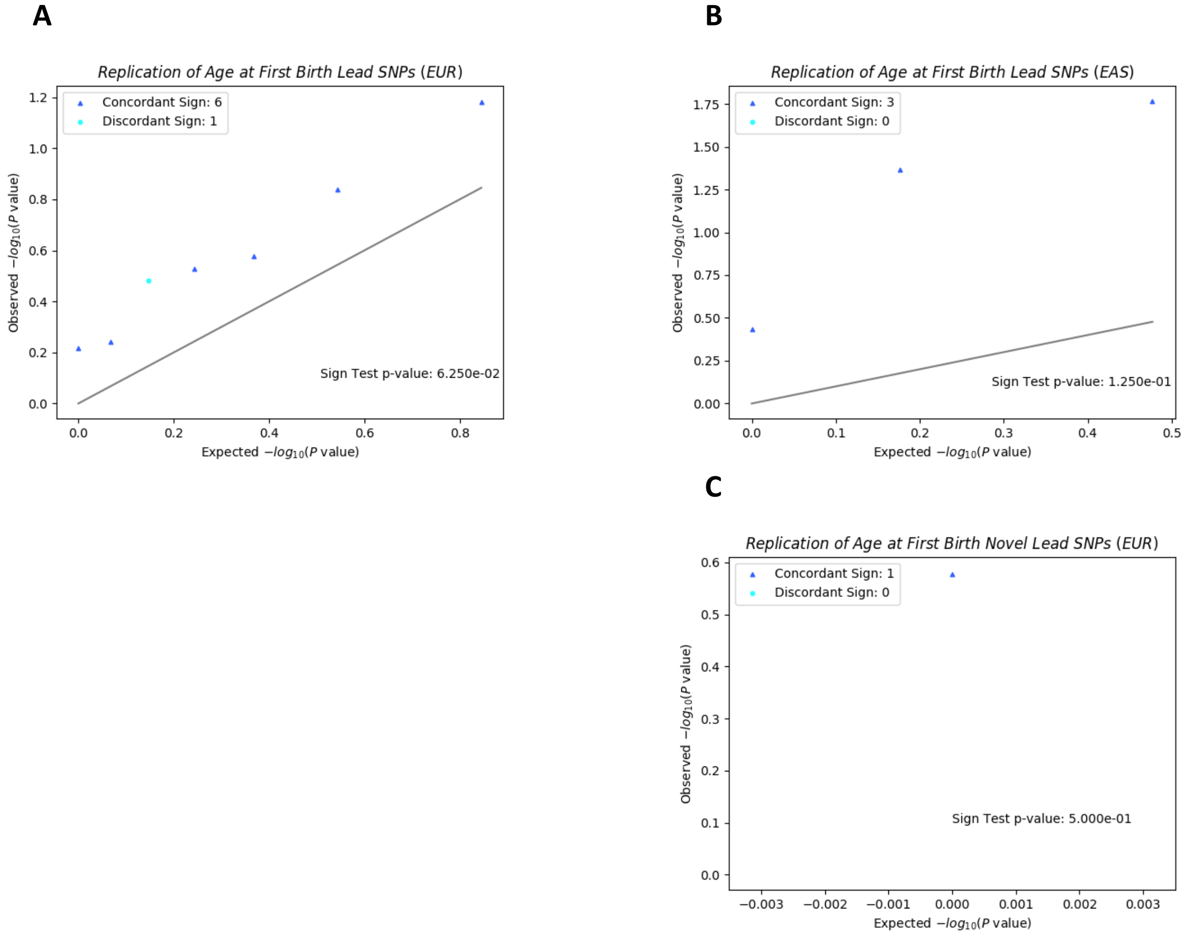


Note: QQ plot and sign test replication of (a) all lead SNPs in MAMA EAS, (b) all lead SNPs in MAMA EUR, (c) and novel lead SNPs in MAMA EUR. There are no novel lead SNPs for EAS. The x-axis corresponds to the uniform distribution of *P* values expected under the null hypothesis, and the y-axis corresponds to the observed *P* values in the replication GWAS output. *P* values are reported on the $-\log_{10}$ scale. The 45-degree line is plotted for reference. The sign of each replication GWAS estimate is compared against the sign of the MAMA estimate for that SNP. SNPs whose sign matches the sign reported in the replication GWAS are marked with a dark blue ▲, and SNPs whose sign does not match the sign reported in the replication GWAS are marked with a light blue •. A sign-test *P* value based on the sign concordance of the replication GWAS estimates and the MAMA estimates is reported in each figure. Details on lead SNP and novel SNP identification and on calculating the sign-test *P* value can be found in Online Methods.

Figure S31. Replication of MAMA Lead SNPs for Basophil Count

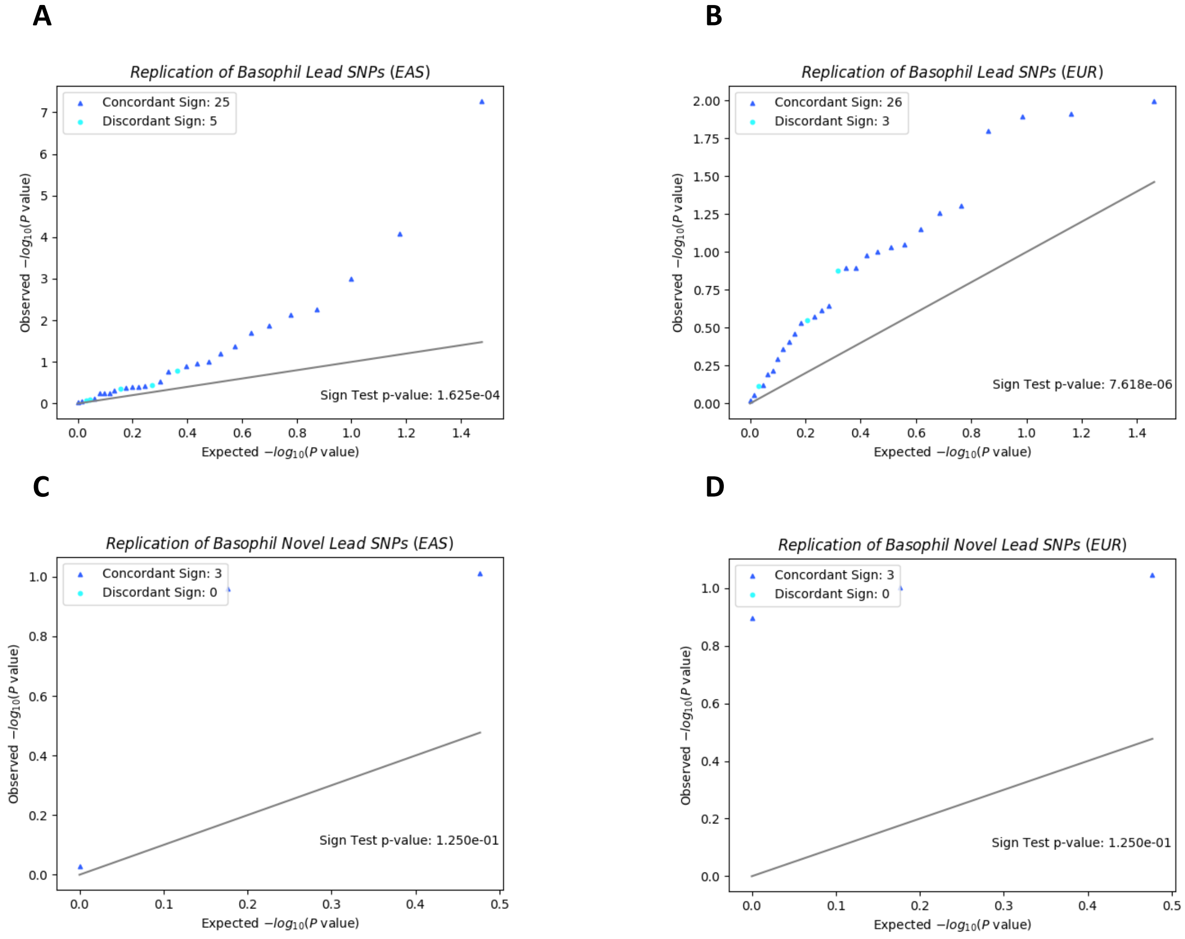


Note: QQ plot and sign test replication of (a) all lead SNPs in MAMA EAS, (b) all lead SNPs in MAMA EUR, (c) novel lead SNPs in MAMA EAS, and (d) novel lead SNPs in MAMA EUR. The x-axis corresponds to the uniform distribution of P values expected under the null hypothesis, and the y-axis corresponds to the observed P values in the MAMA output. P values are reported on the $-\log_{10}$ scale. The 45-degree line is plotted for reference. The sign of each MAMA SNP is compared against an independent replication GWAS. SNPs whose sign matches the sign reported in the replication GWAS are marked with a dark blue ▲, and SNPs whose sign does not match the sign reported in the replication GWAS are marked with a light blue •. Sign test P values are drawn from the binomial distribution with success probability 1/2. Details on lead SNP and novel SNP identification can be found in Online Methods.

Figure S32. Replication of MAMA Lead SNPs for Blood Pressure (Diastolic)

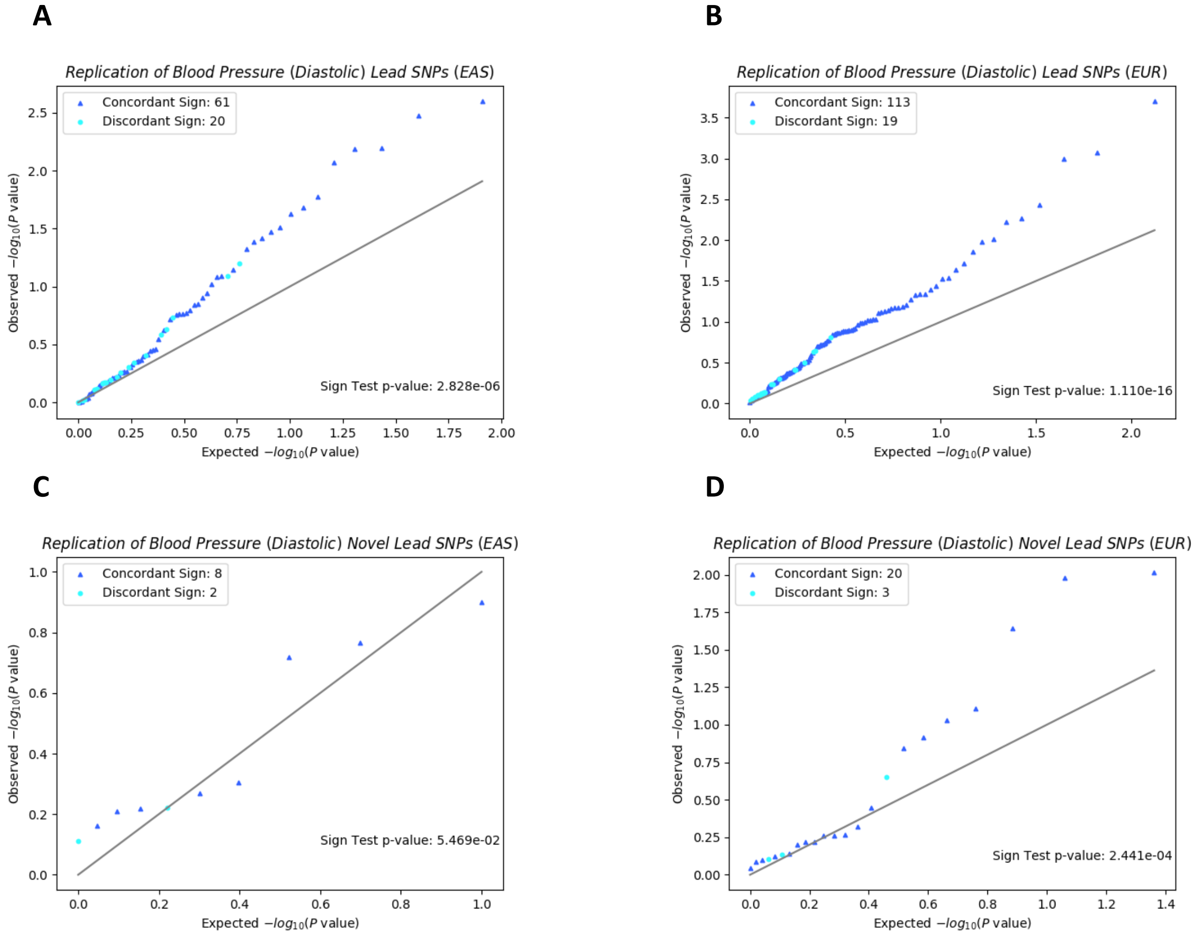


Note: QQ plot and sign test replication of (a) all lead SNPs in MAMA EAS, (b) all lead SNPs in MAMA EUR, (c) novel lead SNPs in MAMA EAS, and (d) novel lead SNPs in MAMA EUR. The x-axis corresponds to the uniform distribution of *P* values expected under the null hypothesis, and the y-axis corresponds to the observed *P* values in the replication GWAS output. *P* values are reported on the $-\log_{10}$ scale. The 45-degree line is plotted for reference. The sign of each replication GWAS estimate is compared against the sign of the MAMA estimate for that SNP. SNPs whose sign matches the sign reported in the replication GWAS are marked with a dark blue ▲, and SNPs whose sign does not match the sign reported in the replication GWAS are marked with a light blue •. A sign-test *P* value based on the sign concordance of the replication GWAS estimates and the MAMA estimates is reported in each figure. Details on lead SNP and novel SNP identification and on calculating the sign-test *P* value can be found in Online Methods.

Figure S33. Replication of MAMA Lead SNPs for Blood Pressure (Systolic)

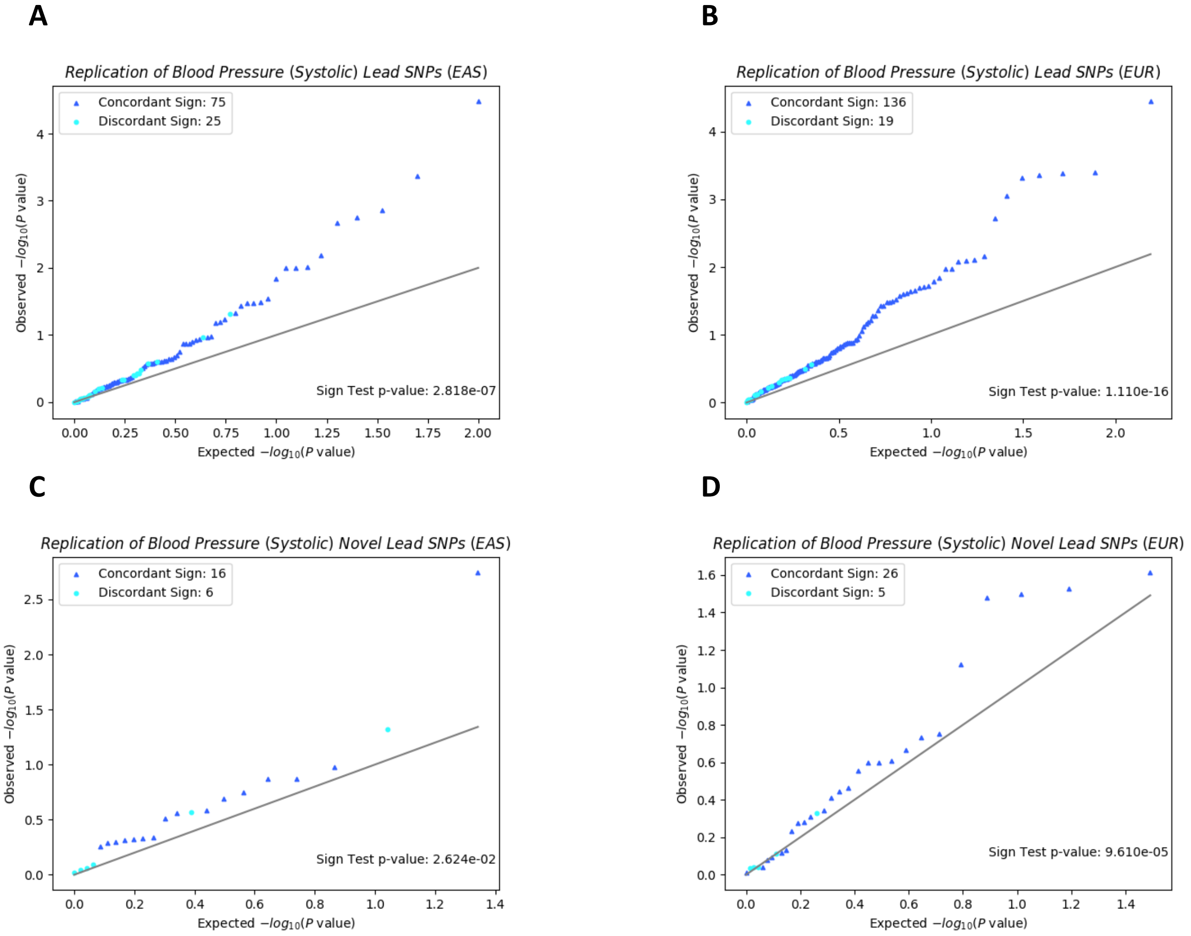


Note: QQ plot and sign test replication of (a) all lead SNPs in MAMA EAS, (b) all lead SNPs in MAMA EUR, (c) novel lead SNPs in MAMA EAS, and (d) novel lead SNPs in MAMA EUR. The x-axis corresponds to the uniform distribution of *P* values expected under the null hypothesis, and the y-axis corresponds to the observed *P* values in the replication GWAS output. *P* values are reported on the $-\log_{10}$ scale. The 45-degree line is plotted for reference. The sign of each replication GWAS estimate is compared against the sign of the MAMA estimate for that SNP. SNPs whose sign matches the sign reported in the replication GWAS are marked with a dark blue ▲, and SNPs whose sign does not match the sign reported in the replication GWAS are marked with a light blue •. A sign-test *P* value based on the sign concordance of the replication GWAS estimates and the MAMA estimates is reported in each figure. Details on lead SNP and novel SNP identification and on calculating the sign-test *P* value can be found in Online Methods.

Figure S34. Replication of MAMA Lead SNPs for BMI

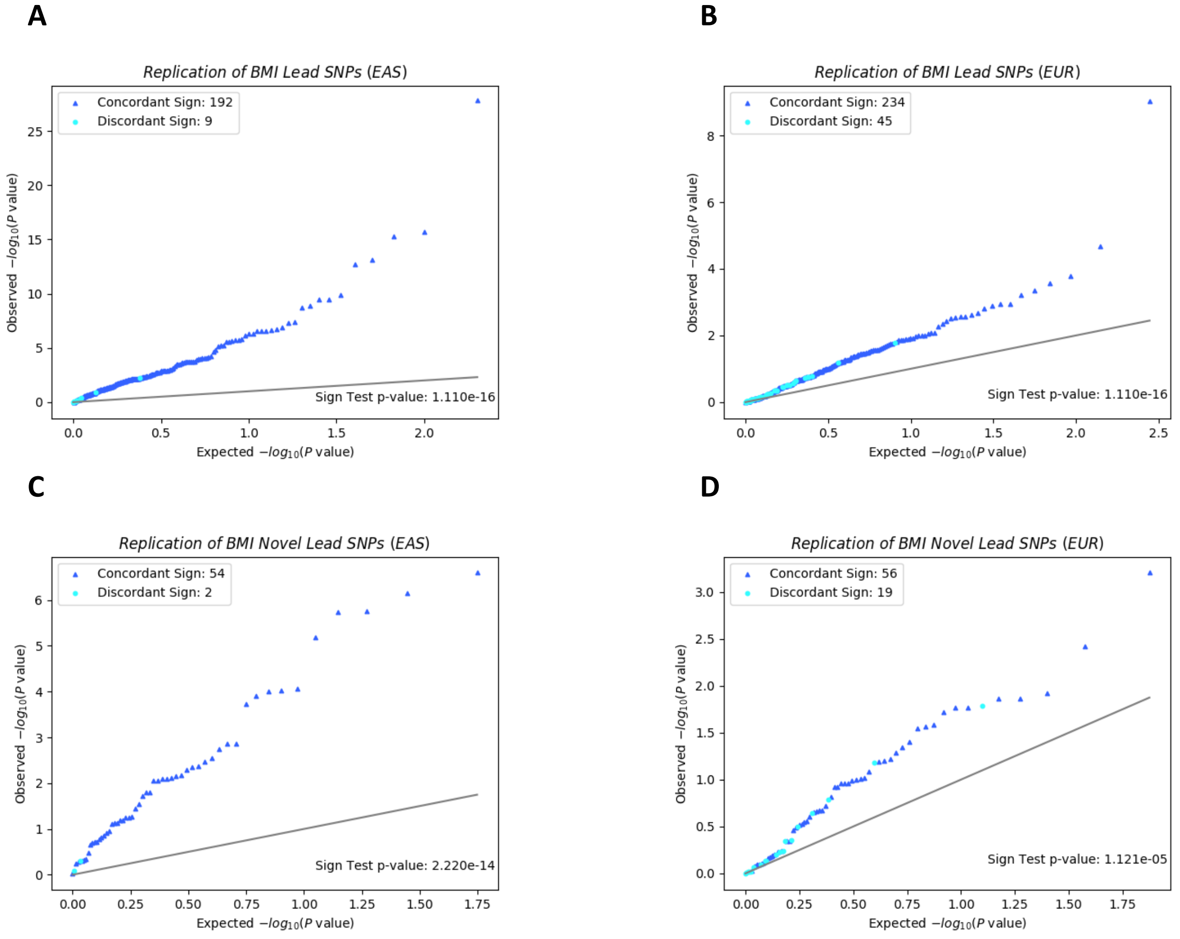


Note: QQ plot and sign test replication of (a) all lead SNPs in MAMA EAS, (b) all lead SNPs in MAMA EUR, (c) novel lead SNPs in MAMA EAS, and (d) novel lead SNPs in MAMA EUR. The x-axis corresponds to the uniform distribution of *P* values expected under the null hypothesis, and the y-axis corresponds to the observed *P* values in the replication GWAS output. *P* values are reported on the $-\log_{10}$ scale. The 45-degree line is plotted for reference. The sign of each replication GWAS estimate is compared against the sign of the MAMA estimate for that SNP. SNPs whose sign matches the sign reported in the replication GWAS are marked with a dark blue ▲, and SNPs whose sign does not match the sign reported in the replication GWAS are marked with a light blue •. A sign-test *P* value based on the sign concordance of the replication GWAS estimates and the MAMA estimates is reported in each figure. Details on lead SNP and novel SNP identification and on calculating the sign-test *P* value can be found in Online Methods.

Figure S35. Replication of MAMA Lead SNPs for Eosinophil Count

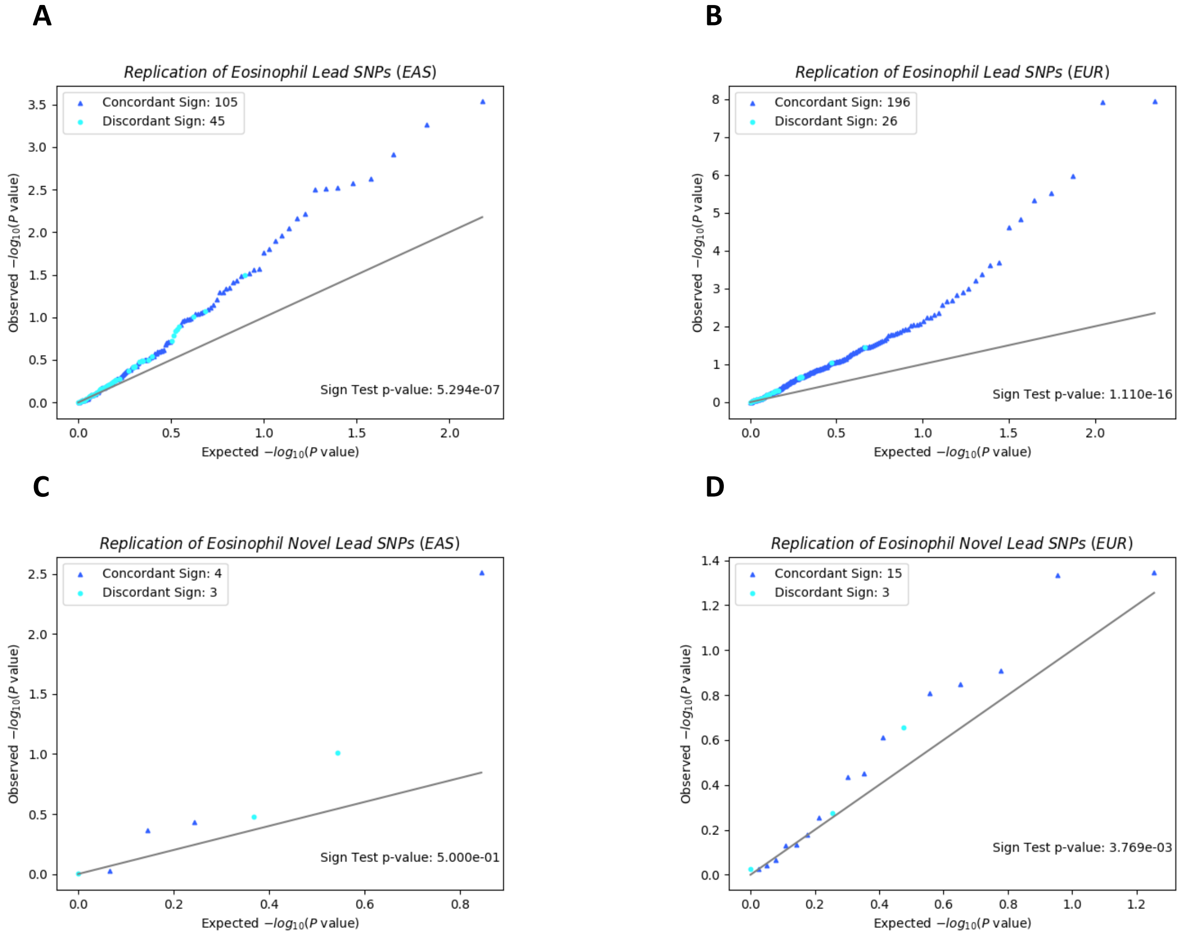


Note: QQ plot and sign test replication of (a) all lead SNPs in MAMA EAS, (b) all lead SNPs in MAMA EUR, (c) novel lead SNPs in MAMA EAS, and (d) novel lead SNPs in MAMA EUR. The x-axis corresponds to the uniform distribution of *P* values expected under the null hypothesis, and the y-axis corresponds to the observed *P* values in the replication GWAS output. *P* values are reported on the $-\log_{10}$ scale. The 45-degree line is plotted for reference. The sign of each replication GWAS estimate is compared against the sign of the MAMA estimate for that SNP. SNPs whose sign matches the sign reported in the replication GWAS are marked with a dark blue ▲, and SNPs whose sign does not match the sign reported in the replication GWAS are marked with a light blue •. A sign-test *P* value based on the sign concordance of the replication GWAS estimates and the MAMA estimates is reported in each figure. Details on lead SNP and novel SNP identification and on calculating the sign-test *P* value can be found in Online Methods.

Figure S36. Replication of MAMA Lead SNPs for Height

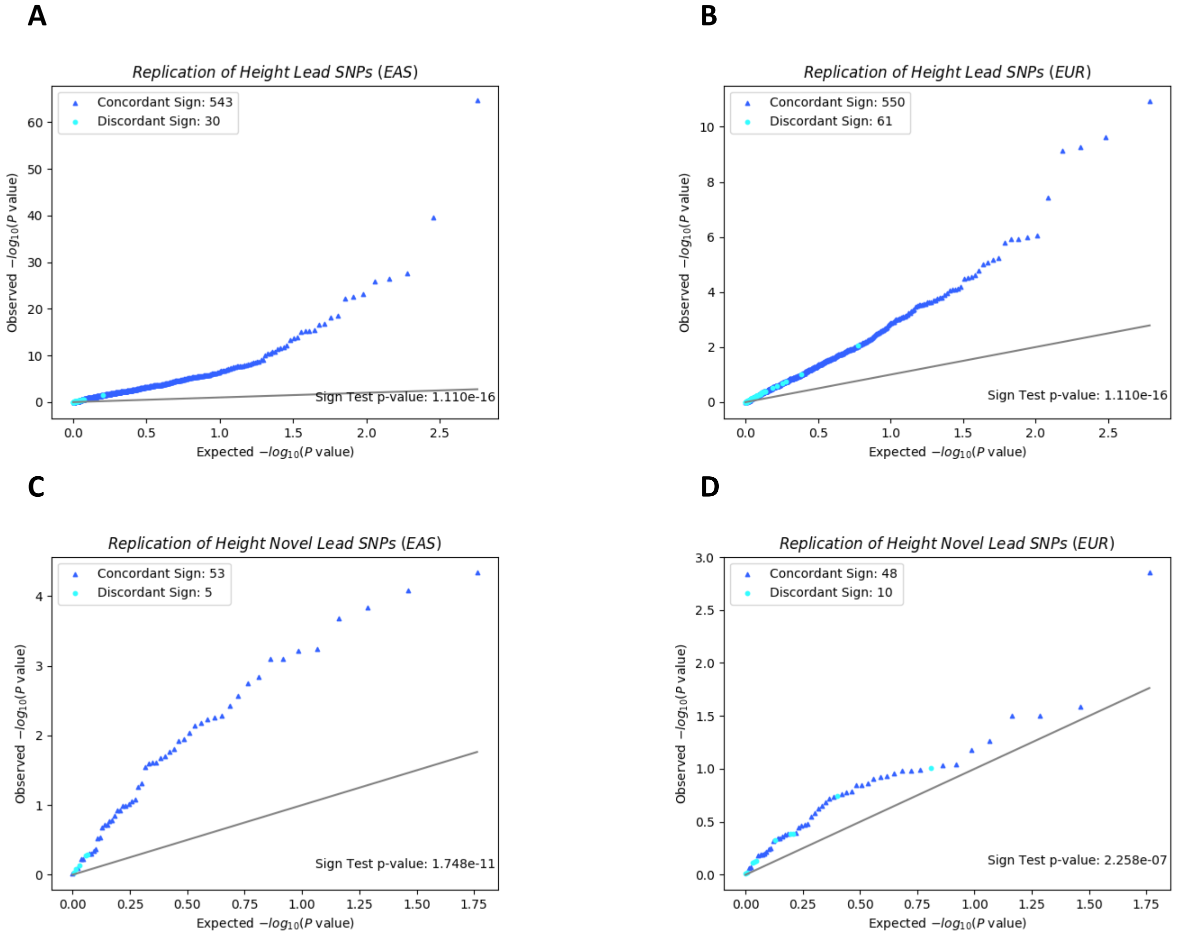


Note: QQ plot and sign test replication of (a) all lead SNPs in MAMA EAS, (b) all lead SNPs in MAMA EUR, (c) novel lead SNPs in MAMA EAS, and (d) novel lead SNPs in MAMA EUR. The x-axis corresponds to the uniform distribution of *P* values expected under the null hypothesis, and the y-axis corresponds to the observed *P* values in the replication GWAS output. *P* values are reported on the $-\log_{10}$ scale. The 45-degree line is plotted for reference. The sign of each replication GWAS estimate is compared against the sign of the MAMA estimate for that SNP. SNPs whose sign matches the sign reported in the replication GWAS are marked with a dark blue ▲, and SNPs whose sign does not match the sign reported in the replication GWAS are marked with a light blue •. A sign-test *P* value based on the sign concordance of the replication GWAS estimates and the MAMA estimates is reported in each figure. Details on lead SNP and novel SNP identification and on calculating the sign-test *P* value can be found in Online Methods.

Figure S37. Replication of MAMA Lead SNPs for Hematocrit

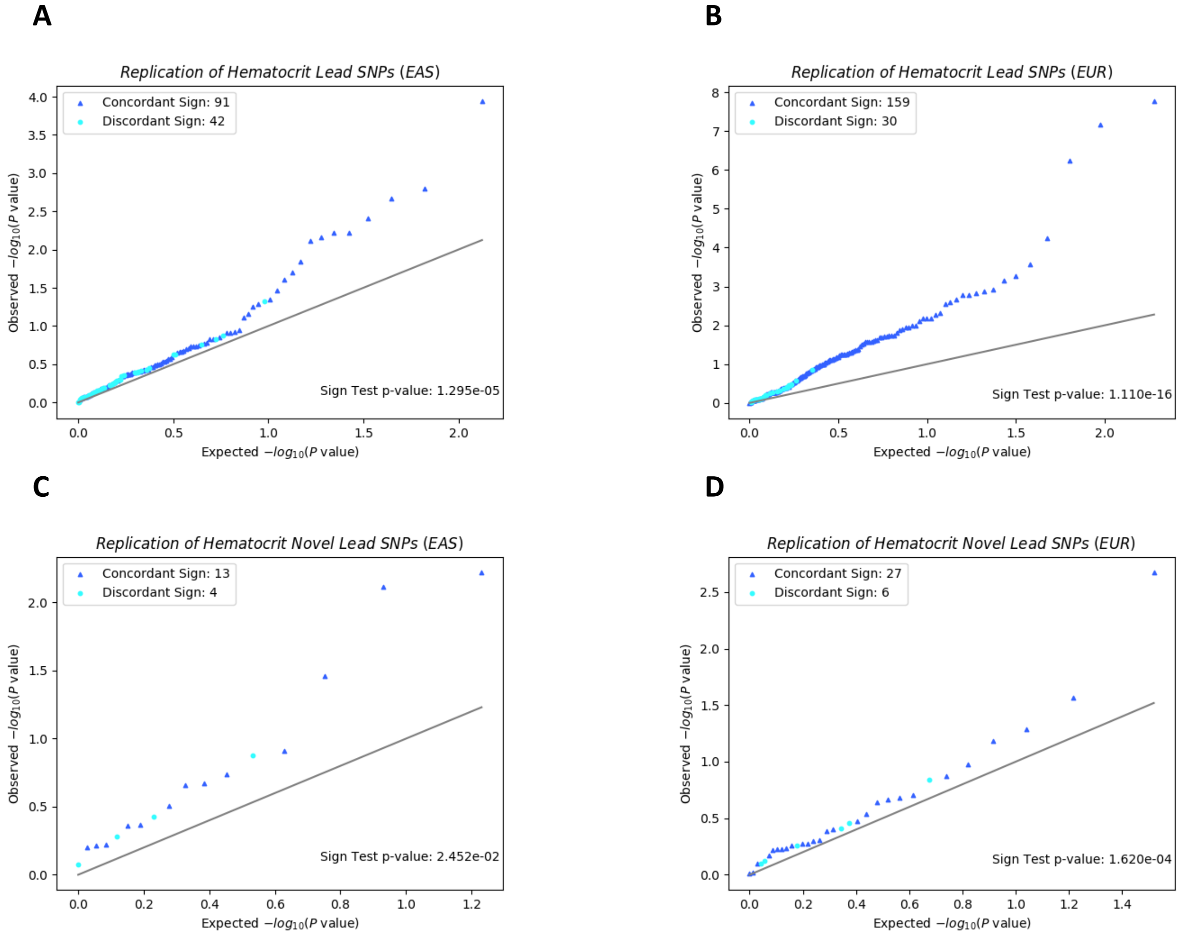


Note: QQ plot and sign test replication of (a) all lead SNPs in MAMA EAS, (b) all lead SNPs in MAMA EUR, (c) novel lead SNPs in MAMA EAS, and (d) novel lead SNPs in MAMA EUR. The x-axis corresponds to the uniform distribution of *P* values expected under the null hypothesis, and the y-axis corresponds to the observed *P* values in the replication GWAS output. *P* values are reported on the $-\log_{10}$ scale. The 45-degree line is plotted for reference. The sign of each replication GWAS estimate is compared against the sign of the MAMA estimate for that SNP. SNPs whose sign matches the sign reported in the replication GWAS are marked with a dark blue ▲, and SNPs whose sign does not match the sign reported in the replication GWAS are marked with a light blue •. A sign-test *P* value based on the sign concordance of the replication GWAS estimates and the MAMA estimates is reported in each figure. Details on lead SNP and novel SNP identification and on calculating the sign-test *P* value can be found in Online Methods.

Figure S38. Replication of MAMA Lead SNPs for Hemoglobin

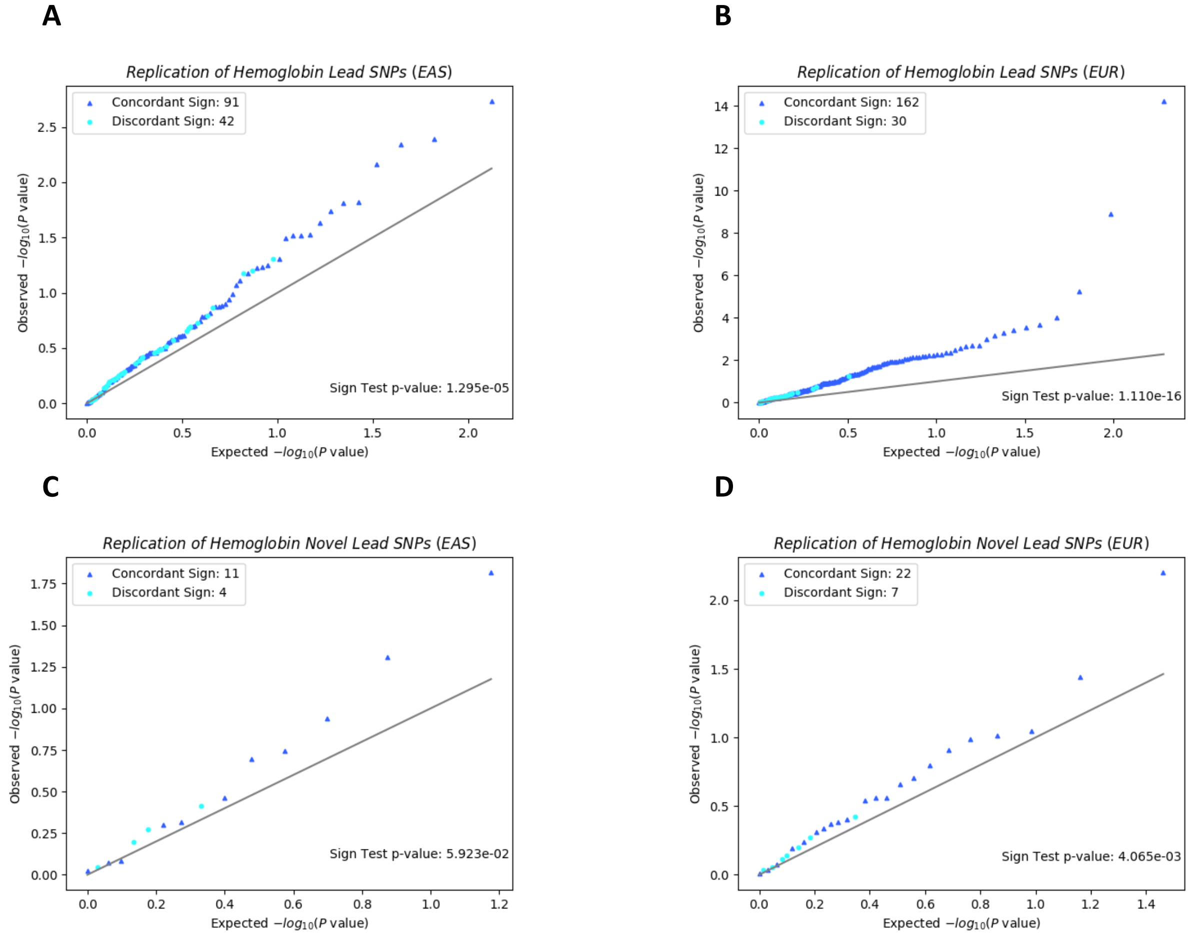


Note: QQ plot and sign test replication of (a) all lead SNPs in MAMA EAS, (b) all lead SNPs in MAMA EUR, (c) novel lead SNPs in MAMA EAS, and (d) novel lead SNPs in MAMA EUR. The x-axis corresponds to the uniform distribution of *P* values expected under the null hypothesis, and the y-axis corresponds to the observed *P* values in the replication GWAS output. *P* values are reported on the $-\log_{10}$ scale. The 45-degree line is plotted for reference. The sign of each replication GWAS estimate is compared against the sign of the MAMA estimate for that SNP. SNPs whose sign matches the sign reported in the replication GWAS are marked with a dark blue ▲, and SNPs whose sign does not match the sign reported in the replication GWAS are marked with a light blue •. A sign-test *P* value based on the sign concordance of the replication GWAS estimates and the MAMA estimates is reported in each figure. Details on lead SNP and novel SNP identification and on calculating the sign-test *P* value can be found in Online Methods.

Figure S39. Replication of MAMA Lead SNPs for Lymphocyte Count

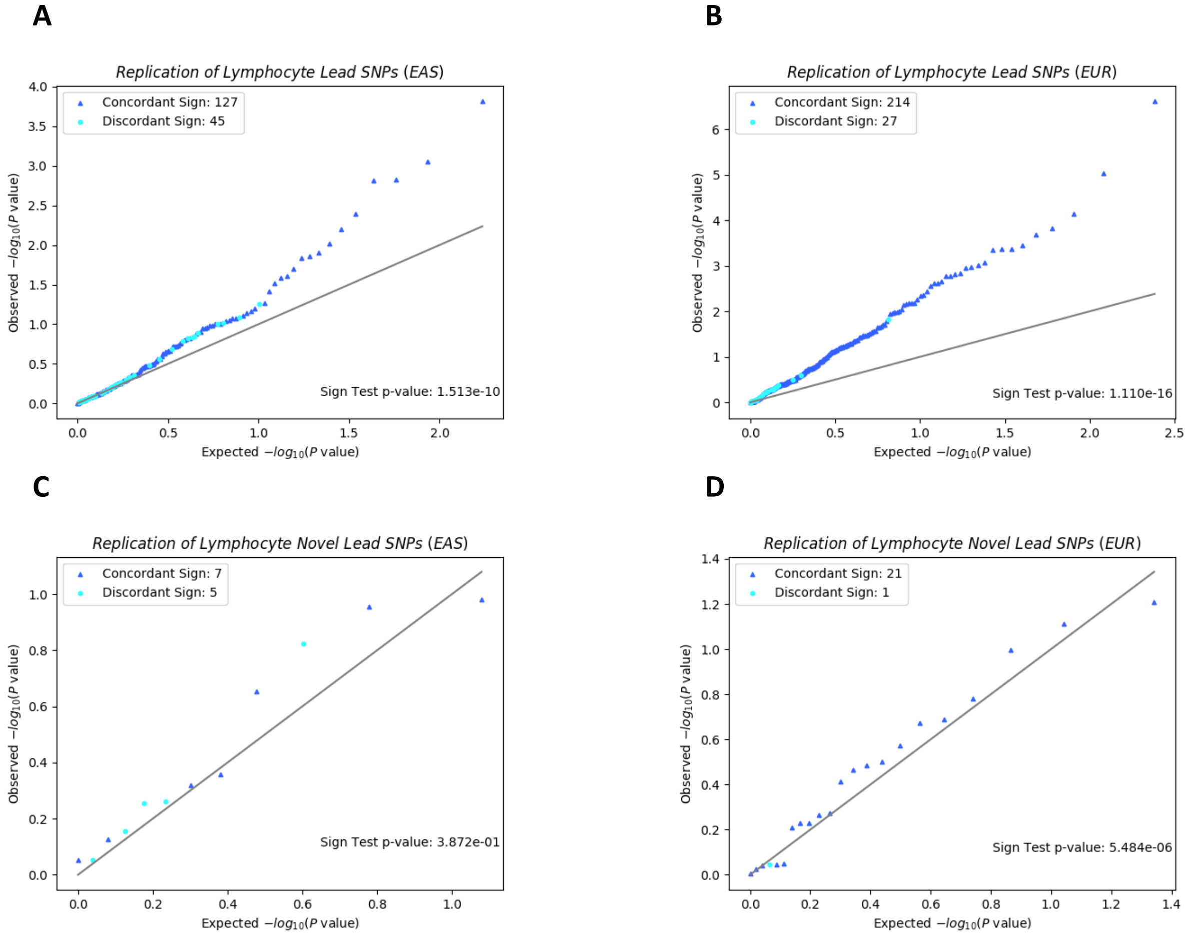


Note: QQ plot and sign test replication of (a) all lead SNPs in MAMA EAS, (b) all lead SNPs in MAMA EUR, (c) novel lead SNPs in MAMA EAS, and (d) novel lead SNPs in MAMA EUR. The x-axis corresponds to the uniform distribution of *P* values expected under the null hypothesis, and the y-axis corresponds to the observed *P* values in the replication GWAS output. *P* values are reported on the $-\log_{10}$ scale. The 45-degree line is plotted for reference. The sign of each replication GWAS estimate is compared against the sign of the MAMA estimate for that SNP. SNPs whose sign matches the sign reported in the replication GWAS are marked with a dark blue ▲, and SNPs whose sign does not match the sign reported in the replication GWAS are marked with a light blue •. A sign-test *P* value based on the sign concordance of the replication GWAS estimates and the MAMA estimates is reported in each figure. Details on lead SNP and novel SNP identification and on calculating the sign-test *P* value can be found in Online Methods.

Figure S40. Replication of MAMA Lead SNPs for Mean Corpuscular Hemoglobin

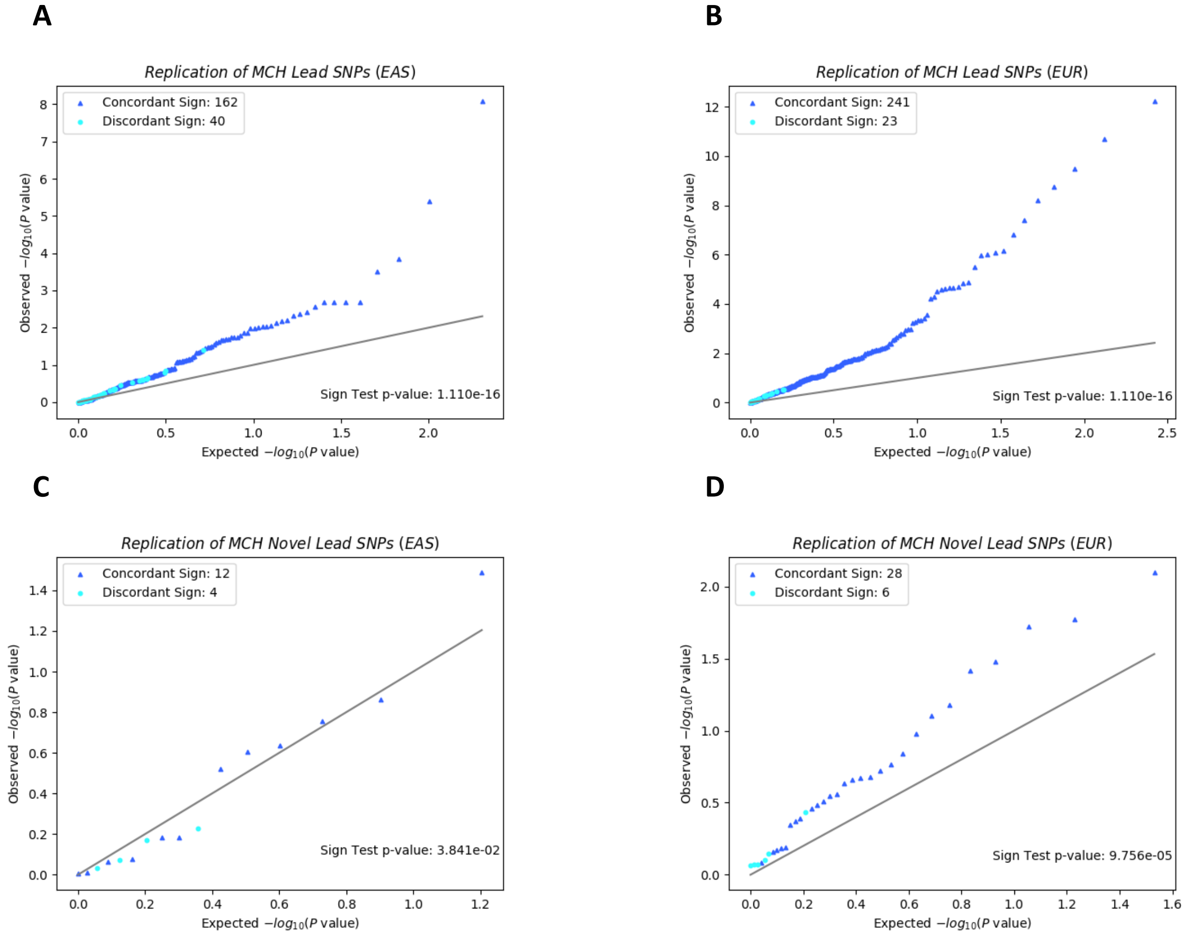


Note: QQ plot and sign test replication of (a) all lead SNPs in MAMA EAS, (b) all lead SNPs in MAMA EUR, (c) novel lead SNPs in MAMA EAS, and (d) novel lead SNPs in MAMA EUR. The x-axis corresponds to the uniform distribution of *P* values expected under the null hypothesis, and the y-axis corresponds to the observed *P* values in the replication GWAS output. *P* values are reported on the $-\log_{10}$ scale. The 45-degree line is plotted for reference. The sign of each replication GWAS estimate is compared against the sign of the MAMA estimate for that SNP. SNPs whose sign matches the sign reported in the replication GWAS are marked with a dark blue ▲, and SNPs whose sign does not match the sign reported in the replication GWAS are marked with a light blue •. A sign-test *P* value based on the sign concordance of the replication GWAS estimates and the MAMA estimates is reported in each figure. Details on lead SNP and novel SNP identification and on calculating the sign-test *P* value can be found in Online Methods.

Figure S41. Replication of MAMA Lead SNPs for Mean Corpuscular Hemoglobin Concentration

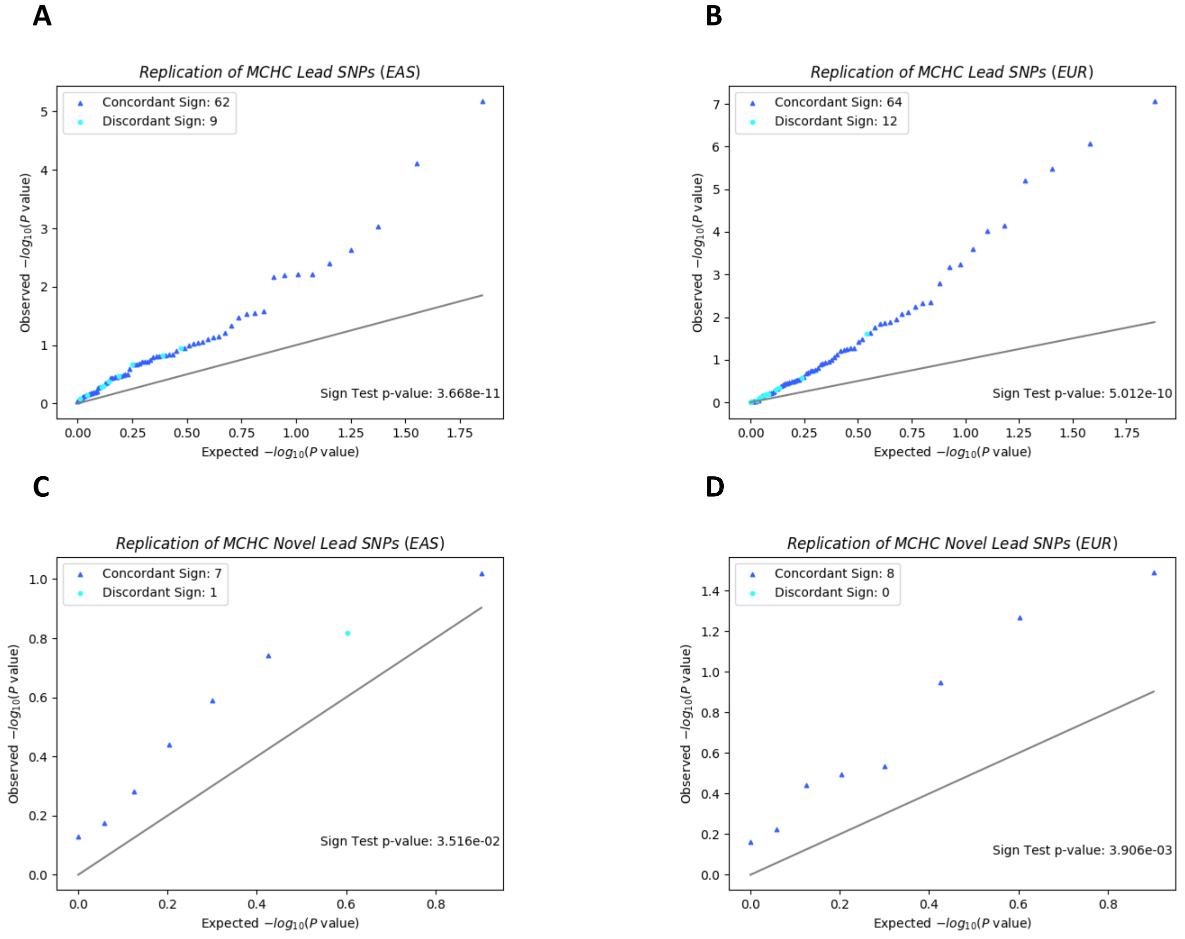


Note: QQ plot and sign test replication of (a) all lead SNPs in MAMA EAS, (b) all lead SNPs in MAMA EUR, (c) novel lead SNPs in MAMA EAS, and (d) novel lead SNPs in MAMA EUR. The x-axis corresponds to the uniform distribution of *P* values expected under the null hypothesis, and the y-axis corresponds to the observed *P* values in the replication GWAS output. *P* values are reported on the -$log_{10}$ scale. The 45-degree line is plotted for reference. The sign of each replication GWAS estimate is compared against the sign of the MAMA estimate for that SNP. SNPs whose sign matches the sign reported in the replication GWAS are marked with a dark blue ▲, and SNPs whose sign does not match the sign reported in the replication GWAS are marked with a light blue •. A sign-test *P* value based on the sign concordance of the replication GWAS estimates and the MAMA estimates is reported in each figure. Details on lead SNP and novel SNP identification and on calculating the sign-test *P* value can be found in Online Methods.

Figure S42. Replication of MAMA Lead SNPs for Mean Corpuscular Volume

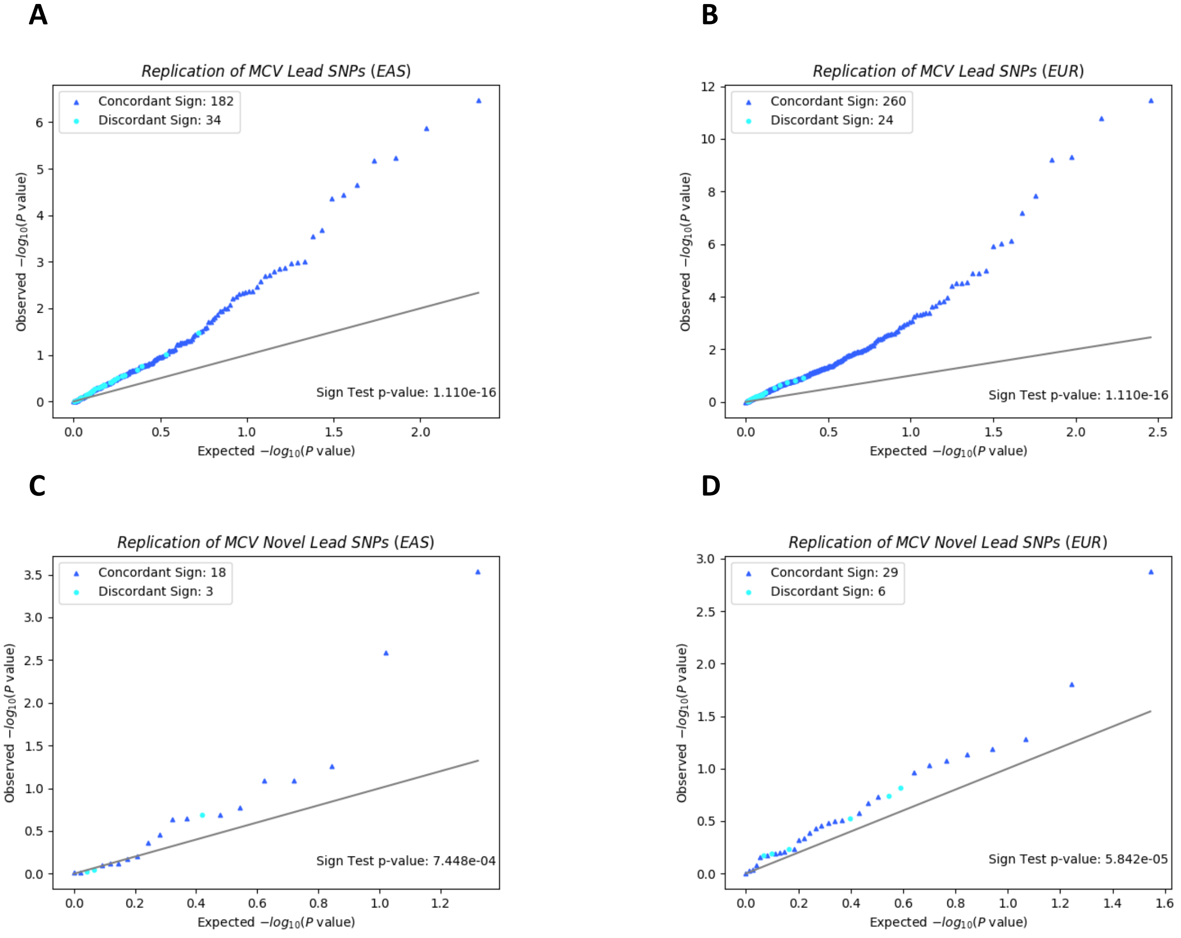


Note: QQ plot and sign test replication of (a) all lead SNPs in MAMA EAS, (b) all lead SNPs in MAMA EUR, (c) novel lead SNPs in MAMA EAS, and (d) novel lead SNPs in MAMA EUR. The x-axis corresponds to the uniform distribution of *P* values expected under the null hypothesis, and the y-axis corresponds to the observed *P* values in the replication GWAS output. *P* values are reported on the -$\log_{10}$ scale. The 45-degree line is plotted for reference. The sign of each replication GWAS estimate is compared against the sign of the MAMA estimate for that SNP. SNPs whose sign matches the sign reported in the replication GWAS are marked with a dark blue ▲, and SNPs whose sign does not match the sign reported in the replication GWAS are marked with a light blue •. A sign-test *P* value based on the sign concordance of the replication GWAS estimates and the MAMA estimates is reported in each figure. Details on lead SNP and novel SNP identification and on calculating the sign-test *P* value can be found in Online Methods.

Figure S43. Replication of MAMA Lead SNPs for Monocyte Count

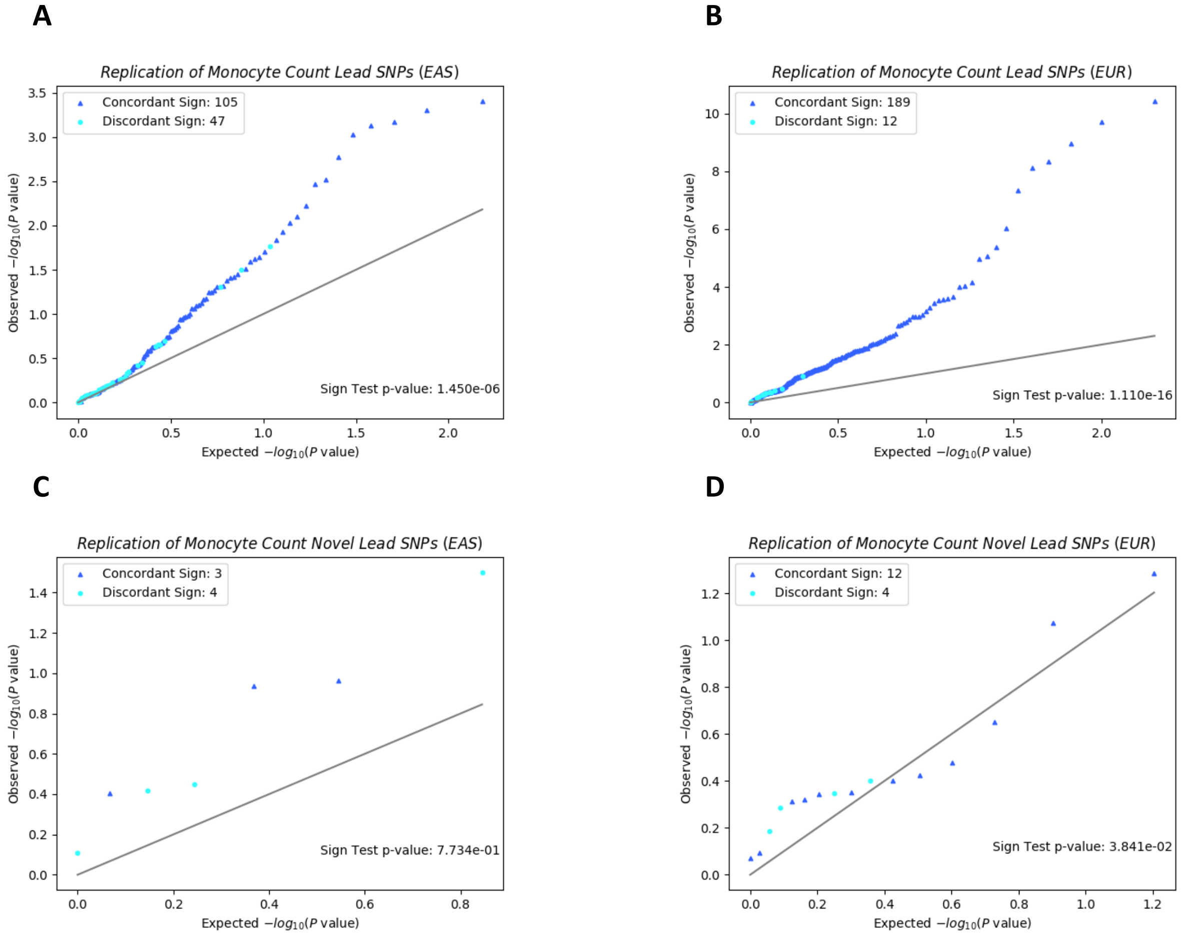


Note: QQ plot and sign test replication of (a) all lead SNPs in MAMA EAS, (b) all lead SNPs in MAMA EUR, (c) novel lead SNPs in MAMA EAS, and (d) novel lead SNPs in MAMA EUR. The x-axis corresponds to the uniform distribution of *P* values expected under the null hypothesis, and the y-axis corresponds to the observed *P* values in the replication GWAS output. *P* values are reported on the $-\log_{10}$ scale. The 45-degree line is plotted for reference. The sign of each replication GWAS estimate is compared against the sign of the MAMA estimate for that SNP. SNPs whose sign matches the sign reported in the replication GWAS are marked with a dark blue ▲, and SNPs whose sign does not match the sign reported in the replication GWAS are marked with a light blue •. A sign-test *P* value based on the sign concordance of the replication GWAS estimates and the MAMA estimates is reported in each figure. Details on lead SNP and novel SNP identification and on calculating the sign-test *P* value can be found in Online Methods.

Figure S44. Replication of MAMA Lead SNPs for Neutrophil Count

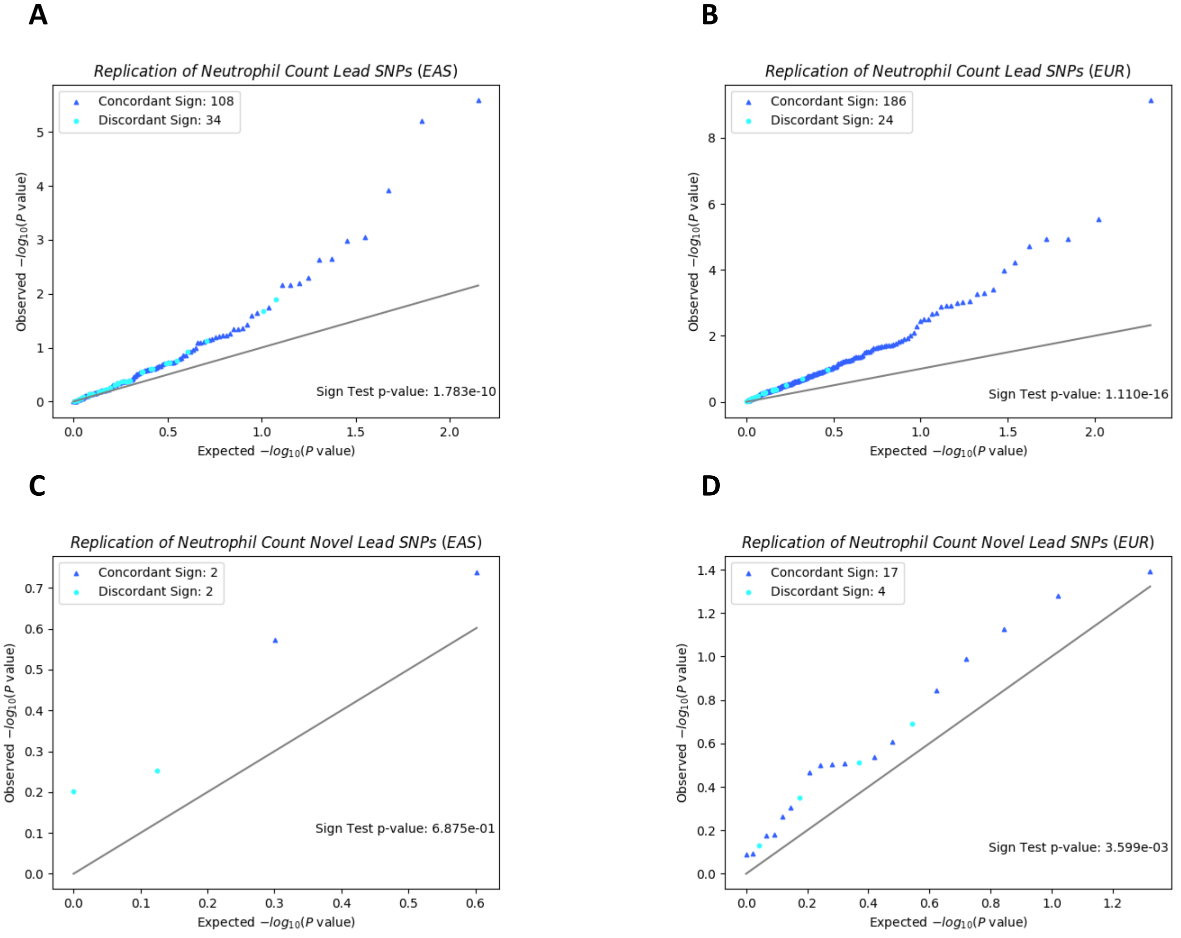


Note: QQ plot and sign test replication of (a) all lead SNPs in MAMA EAS, (b) all lead SNPs in MAMA EUR, (c) novel lead SNPs in MAMA EAS, and (d) novel lead SNPs in MAMA EUR. The x-axis corresponds to the uniform distribution of *P* values expected under the null hypothesis, and the y-axis corresponds to the observed *P* values in the replication GWAS output. *P* values are reported on the $-\log_{10}$ scale. The 45-degree line is plotted for reference. The sign of each replication GWAS estimate is compared against the sign of the MAMA estimate for that SNP. SNPs whose sign matches the sign reported in the replication GWAS are marked with a dark blue ▲, and SNPs whose sign does not match the sign reported in the replication GWAS are marked with a light blue •. A sign-test *P* value based on the sign concordance of the replication GWAS estimates and the MAMA estimates is reported in each figure. Details on lead SNP and novel SNP identification and on calculating the sign-test *P* value can be found in Online Methods.

Figure S45. Replication of MAMA Lead SNPs for Platelet Count

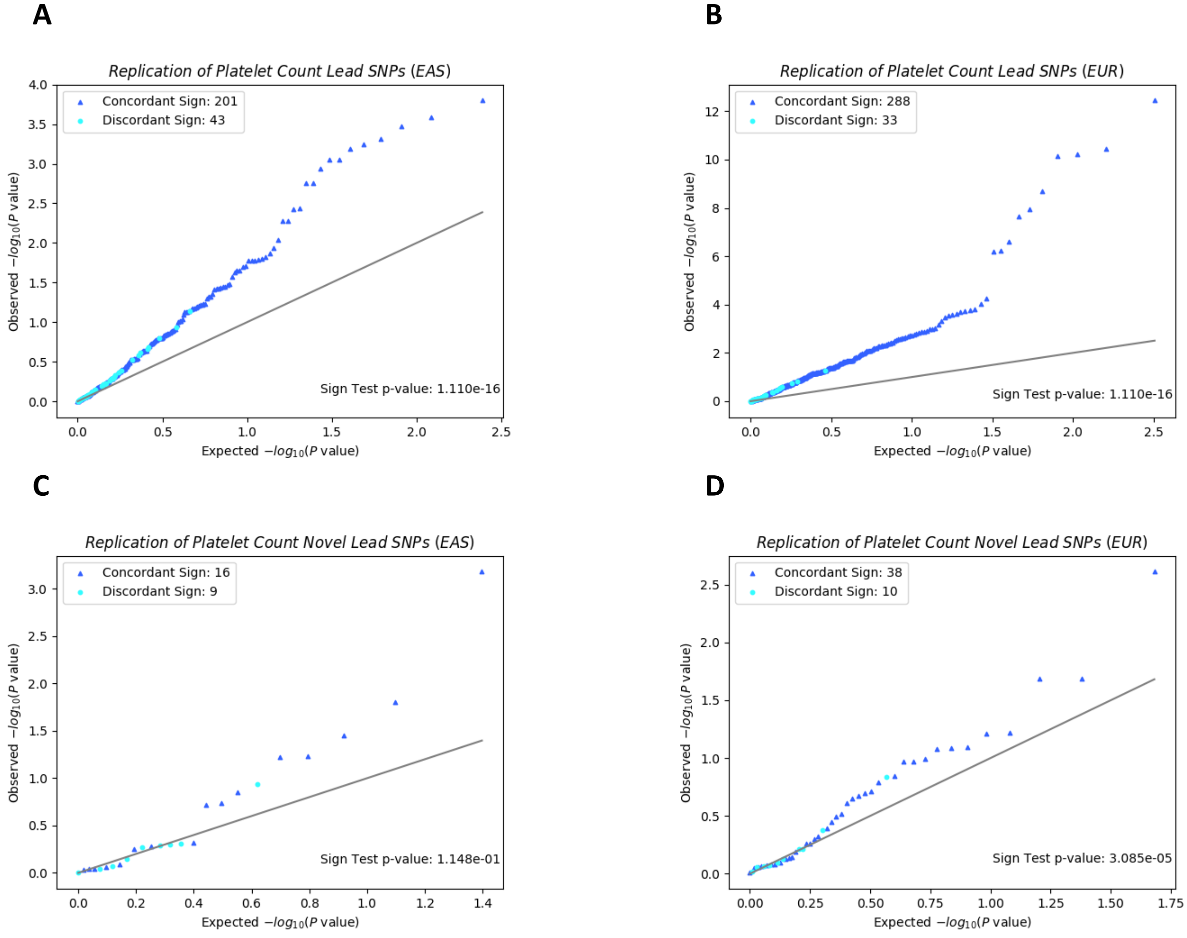


Note: QQ plot and sign test replication of (a) all lead SNPs in MAMA EAS, (b) all lead SNPs in MAMA EUR, (c) novel lead SNPs in MAMA EAS, and (d) novel lead SNPs in MAMA EUR. The x-axis corresponds to the uniform distribution of *P* values expected under the null hypothesis, and the y-axis corresponds to the observed *P* values in the replication GWAS output. *P* values are reported on the $-\log_{10}$ scale. The 45-degree line is plotted for reference. The sign of each replication GWAS estimate is compared against the sign of the MAMA estimate for that SNP. SNPs whose sign matches the sign reported in the replication GWAS are marked with a dark blue ▲, and SNPs whose sign does not match the sign reported in the replication GWAS are marked with a light blue •. A sign-test *P* value based on the sign concordance of the replication GWAS estimates and the MAMA estimates is reported in each figure. Details on lead SNP and novel SNP identification and on calculating the sign-test *P* value can be found in Online Methods.

Figure S46. Replication of MAMA Lead SNPs for Red Blood Cell Count

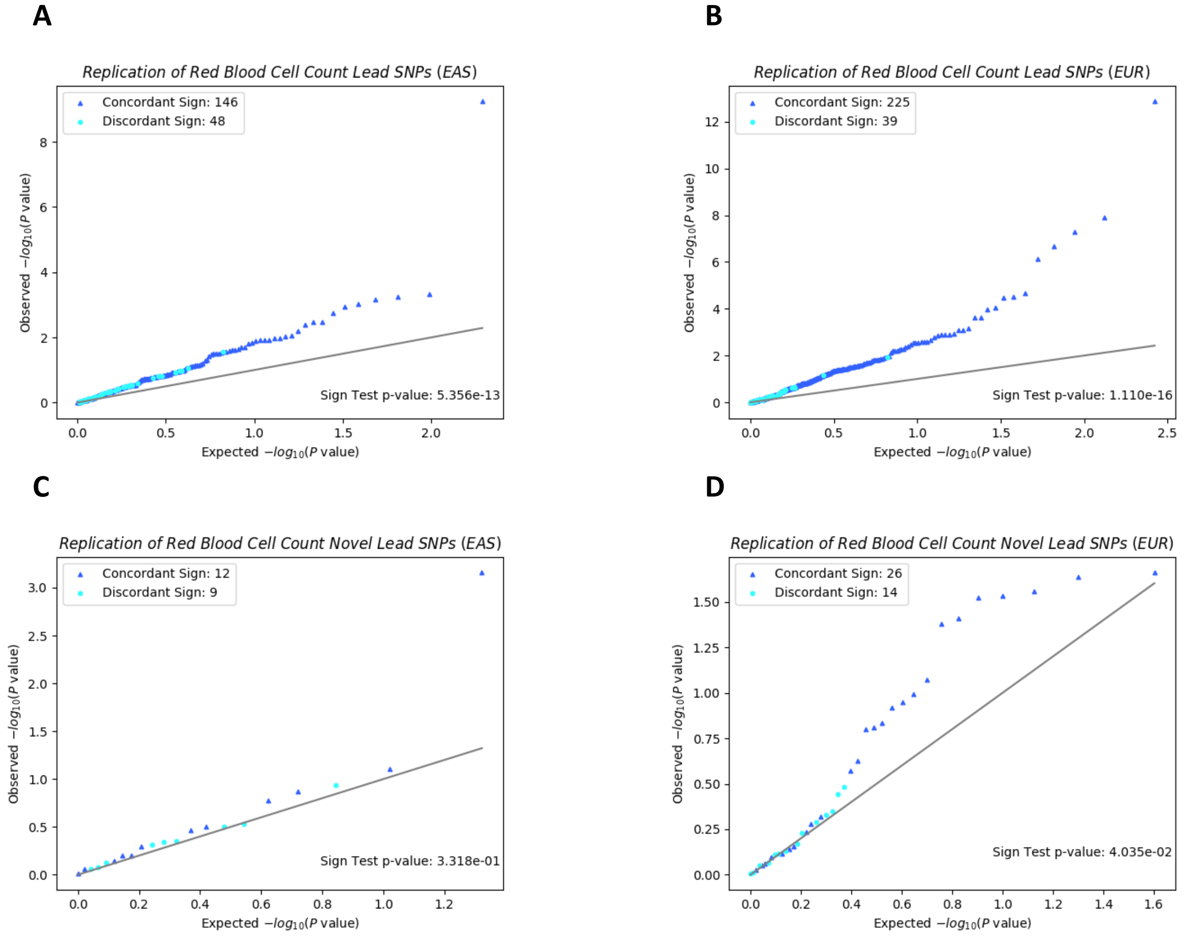


Note: QQ plot and sign test replication of (a) all lead SNPs in MAMA EAS, (b) all lead SNPs in MAMA EUR, (c) novel lead SNPs in MAMA EAS, and (d) novel lead SNPs in MAMA EUR. The x-axis corresponds to the uniform distribution of *P* values expected under the null hypothesis, and the y-axis corresponds to the observed *P* values in the replication GWAS output. *P* values are reported on the -$\log_{10}$ scale. The 45-degree line is plotted for reference. The sign of each replication GWAS estimate is compared against the sign of the MAMA estimate for that SNP. SNPs whose sign matches the sign reported in the replication GWAS are marked with a dark blue ▲, and SNPs whose sign does not match the sign reported in the replication GWAS are marked with a light blue •. A sign-test *P* value based on the sign concordance of the replication GWAS estimates and the MAMA estimates is reported in each figure. Details on lead SNP and novel SNP identification and on calculating the sign-test *P* value can be found in Online Methods.

Figure S47. Replication of MAMA Lead SNPs for Smoking Initiation

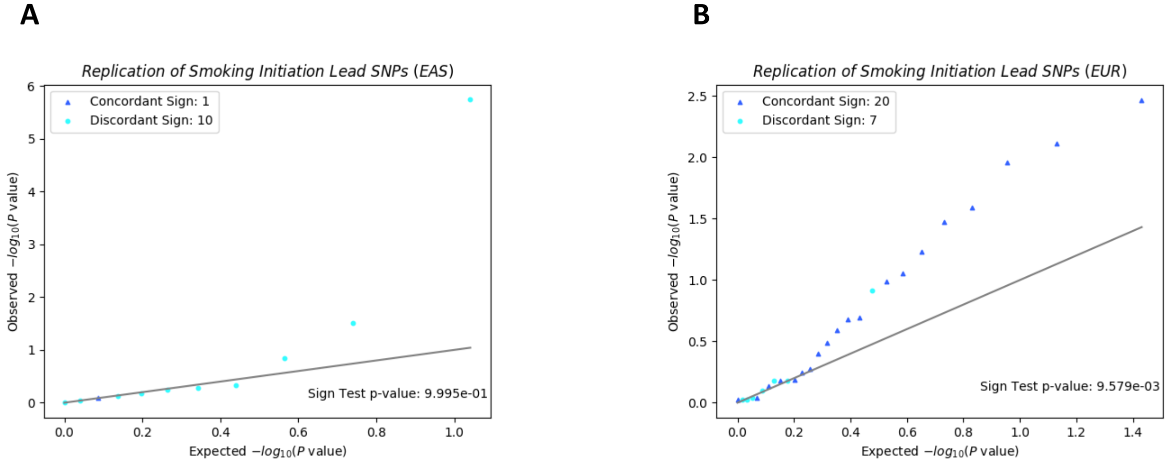


Note: QQ plot and sign test replication of (a) all lead SNPs in MAMA EAS and (b) all lead SNPs in MAMA EUR. There are no novel lead SNPs for EAS or EUR. The x-axis corresponds to the uniform distribution of *P* values expected under the null hypothesis, and the y-axis corresponds to the observed *P* values in the replication GWAS output. *P* values are reported on the $-\log_{10}$ scale. The 45-degree line is plotted for reference. The sign of each replication GWAS estimate is compared against the sign of the MAMA estimate for that SNP. SNPs whose sign matches the sign reported in the replication GWAS are marked with a dark blue ▲, and SNPs whose sign does not match the sign reported in the replication GWAS are marked with a light blue •. A sign-test *P* value based on the sign concordance of the replication GWAS estimates and the MAMA estimates is reported in each figure. Details on lead SNP and novel SNP identification and on calculating the sign-test *P* value can be found in Online Methods.

Figure S48. Replication of MAMA Lead SNPs for Weight

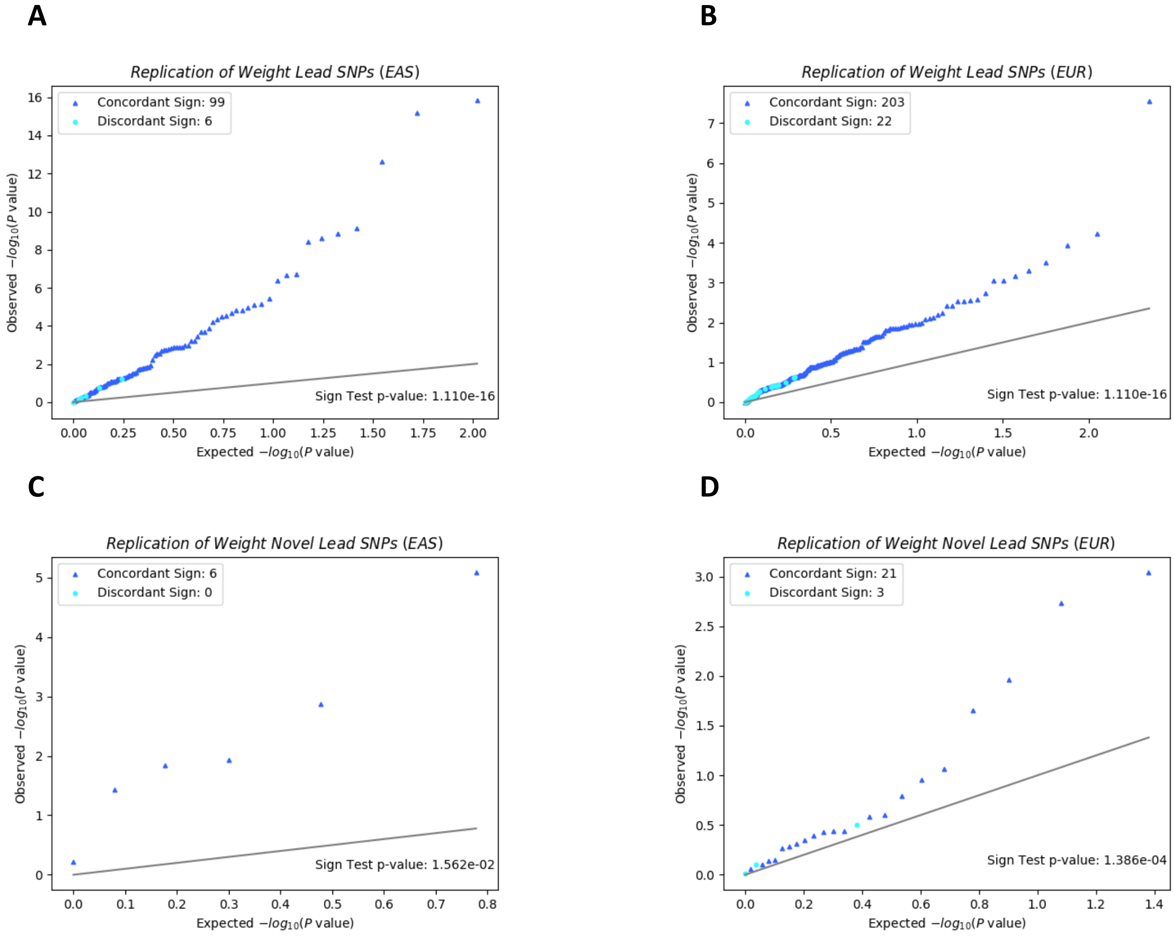


Note: QQ plot and sign test replication of (a) all lead SNPs in MAMA EAS, (b) all lead SNPs in MAMA EUR, (c) novel lead SNPs in MAMA EAS, and (d) novel lead SNPs in MAMA EUR. The x-axis corresponds to the uniform distribution of *P* values expected under the null hypothesis, and the y-axis corresponds to the observed *P* values in the replication GWAS output. *P* values are reported on the $-\log_{10}$ scale. The 45-degree line is plotted for reference. The sign of each replication GWAS estimate is compared against the sign of the MAMA estimate for that SNP. SNPs whose sign matches the sign reported in the replication GWAS are marked with a dark blue ▲, and SNPs whose sign does not match the sign reported in the replication GWAS are marked with a light blue •. A sign-test *P* value based on the sign concordance of the replication GWAS estimates and the MAMA estimates is reported in each figure. Details on lead SNP and novel SNP identification and on calculating the sign-test *P* value can be found in Online Methods.

Figure S49. Replication of MAMA Lead SNPs for White Blood Cell Count

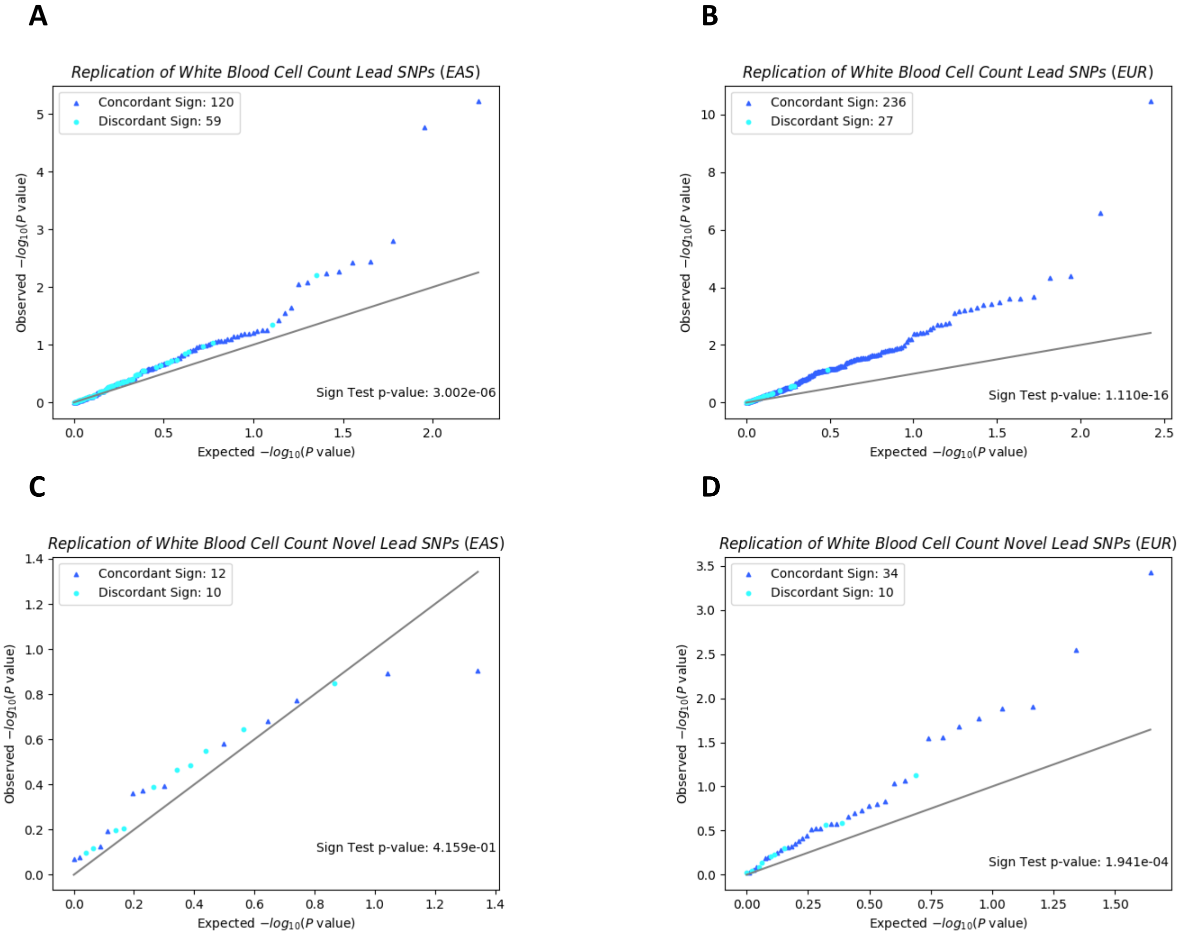


Note: QQ plot and sign test replication of (a) all lead SNPs in MAMA EAS, (b) all lead SNPs in MAMA EUR, (c) novel lead SNPs in MAMA EAS, and (d) novel lead SNPs in MAMA EUR. The x-axis corresponds to the uniform distribution of *P* values expected under the null hypothesis, and the y-axis corresponds to the observed *P* values in the replication GWAS output. *P* values are reported on the $-\log_{10}$ scale. The 45-degree line is plotted for reference. The sign of each replication GWAS estimate is compared against the sign of the MAMA estimate for that SNP. SNPs whose sign matches the sign reported in the replication GWAS are marked with a dark blue ▲, and SNPs whose sign does not match the sign reported in the replication GWAS are marked with a light blue •. A sign-test *P* value based on the sign concordance of the replication GWAS estimates and the MAMA estimates is reported in each figure. Details on lead SNP and novel SNP identification and on calculating the sign-test *P* value can be found in Online Methods.

Figure S50. LD Score Histograms and LD Score Correlation in Two Populations.

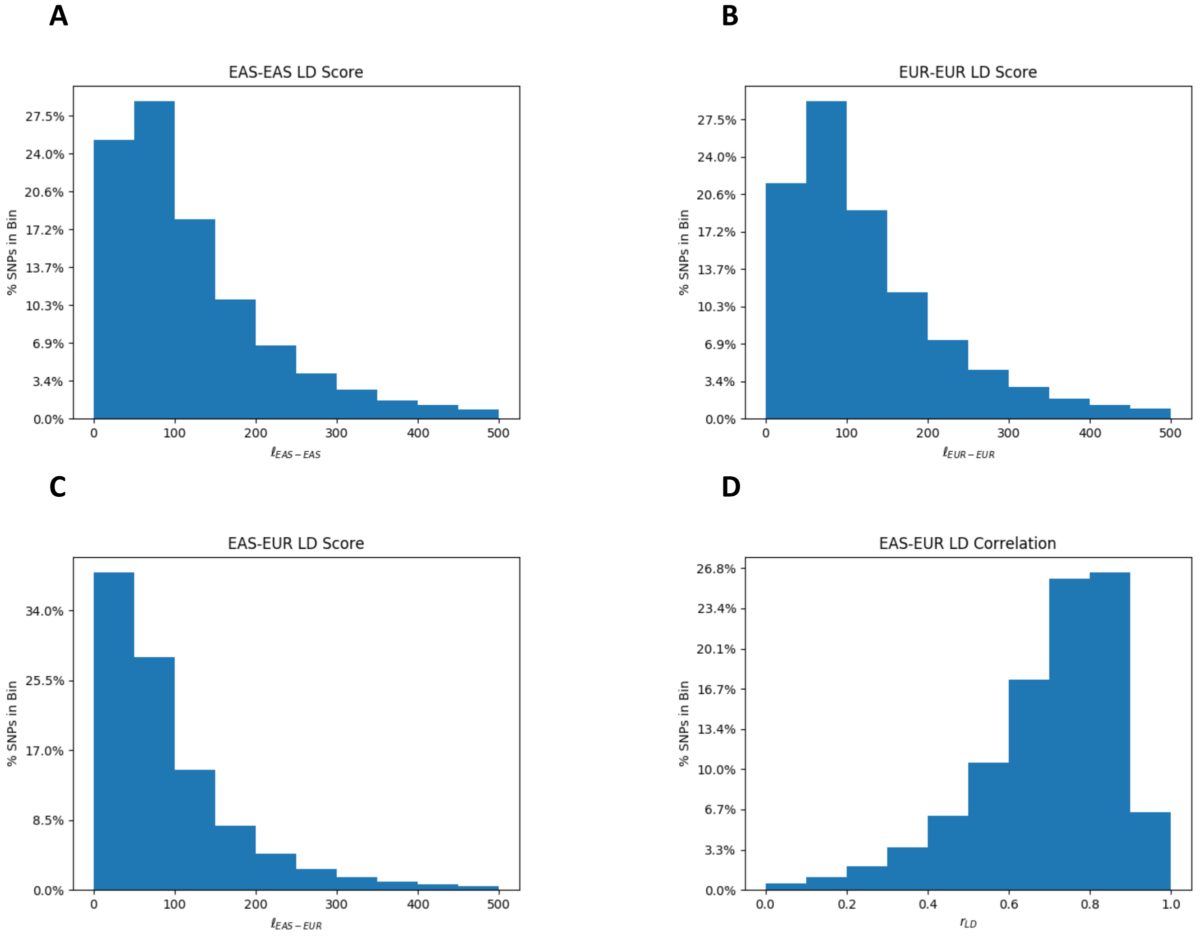


Note: Distribution of standardized unit LD scores in (a) EAS, (b) EUR, and (c) EAS-EUR. Distribution of EAS-EUR LD score correlation (d) defined as the EAS-EUR LD score over the square root of the product of the EAS LD score and the EUR LD score.
